# Establishment of Proximity-dependent Biotinylation Approaches in Different Plant Model Systems

**DOI:** 10.1101/701425

**Authors:** Deepanksha Arora, Nikolaj B. Abel, Chen Liu, Petra Van Damme, Klaas Yperman, Lam Dai Vu, Jie Wang, Anna Tornkvist, Francis Impens, Barbara Korbei, Dominique Eeckhout, Jelle Van Leene, Alain Goossens, Geert De Jaeger, Thomas Ott, Panagiotis Moschou, Daniël Van Damme

**Author notes:** joint first authors (Deepanksha Arora, Nikolaj B. Abel, Chen Liu, Petra Van Damme). Joint senior and corresponding authors. The author(s) responsible for distribution of materials integral to the findings presented in this article are: Geert De Jaeger, Thomas Ott, Panagiotis Moshou and Daniël Van Damme.

## Abstract

Proximity-dependent biotin labelling (PDL) uses a promiscuous biotin ligase (PBL) or a peroxidase fused to a protein of interest. This enables covalent biotin labelling of proteins and allows subsequent capture and identification of interacting and neighbouring proteins without the need for the protein complex to remain intact. To date, only few papers report on the use of PDL in plants. Here we present the results of a systematic study applying a variety of PDL approaches in several plant systems using various conditions and bait proteins. We show that TurboID is the most promiscuous variant in several plant model systems and establish protocols which combine Mass Spectrometry-based analysis with harsh extraction and washing conditions. We demonstrate the applicability of TurboID in capturing membrane-associated protein interactomes using *Lotus japonicus* symbiotically active receptor kinases as test-case. We further benchmark the efficiency of various PBLs in comparison with one-step affinity purification approaches. We identified both known as well as novel interactors of the endocytic TPLATE complex. We furthermore present a straightforward strategy to identify both non-biotinylated as well as biotinylated peptides in a single experimental setup. Finally, we provide initial evidence that our approach has the potential to infer structural information of protein complexes.

## INTRODUCTION

Protein-protein interaction (PPI) studies often fail to capture low-affinity interactions as these are usually not maintained following cell lysis, protein extraction and protein complex purification. Particularly, this is the case for PPI’s of integral membrane proteins because of the harsh conditions during protein extraction and purification. Proximity-dependent biotin labelling (PDL) on the contrary, uses covalent biotinylation of proteins that are interactors or near-neighbours of a bait protein of interest *in vivo* (Varnaite and MacNeill, 2016). Hence, to identify interactions, they do not need to remain intact during purification. Although biotin is an essential cofactor for a small number of omnipresent biotin-dependent enzymes involved mainly in the transfer of CO_2_ during HCO_3_^-^-dependent carboxylation reactions, biotinylation is a relatively rare *in vivo* protein modification. Moreover, biotinylated proteins can be selectively isolated with high affinity using streptavidin-biotin pairing. PDL, therefore, permits the identification of both high and low-affinity *in vivo* interactions.

Analogues to DamID in which a prokaryotic *Dam* methylase is fused to a protein of interest to monitor DNA-protein interactions in eukaryotes (van Steensel and Henikoff, 2000), the principle of PDL allows the capture of PPIs. More specifically, PDL is based on the fact that native biotin ligases, e.g. the *Escherichia coli* BirA catalyzes a two-step reaction: first, the generation of reactive biotinyl-AMP (biotinoyl-5’-AMP or bioAMP) from biotin and ATP, and second, the attachment of that bioAMP to a specific lysine of the target protein. Engineered PBLs have a significantly reduced affinity for the reactive bioAMP intermediate (Choi-Rhee et al., 2004; Kim and Roux, 2016). This intermediate is prematurely released and, due to its high reactivity, will interact with neighbouring primary amines (e.g. lysine). Therefore, these variants lead to promiscuous labelling despite their lower affinity for biotin compared to native biotin ligases.

There are several variations of PDL. The first-generation enzymes used for PDL are based on the *E. coli* biotin ligase BirA (Roux et al., 2012). The mutant BirA, designated BirA* (R118G) (Kwon and Beckett, 2000), referred hereafter as BioID, represents a monomeric protein of 35.3 kDa, and was the first PBL variant used for PDL (Choi-Rhee et al., 2004; Cronan, 2005; Kim and Roux, 2016). A second-generation PBL, called BioID2, was derived from the *Aquifex aeolicus* biotin ligase (Kim and Roux, 2016). BioID2, which naturally lacks a DNA-binding domain that is present in the larger BirA, is approximately one-third smaller than BioID, potentially reducing sterical hindrance of the bait protein (Kim et al., 2016). The third-generation PBLs, called TurboID and mini-Turbo (mTurbo), are derived from the directed evolution of BirA in yeast. These two variants showed maximal activity at 30°C, whereas the previous variants show maximal activity at higher temperatures (Branon et al., 2018). TurboID has the same size as the original BioID tag, albeit with 14 amino acid mutations that greatly increase its labelling efficiency. mTurbo has 12 out of the 14 mutations. The N-terminal DNA-binding domain was deleted to reduce its size (28 versus 35 kDa), which also slightly impacted on its labelling efficiency by reducing it ~2-fold. The first and second-generation PBLs required approximately 18 to 24 h of labelling or sometimes even much longer to produce detectable levels of protein biotinylation, while the TurboID variants required a labelling time in the range of 1 h or less in the various eukaryotic, non-plant systems tested so far (Branon et al., 2018).

PDL has its intrinsic advantages and limitations. In the presence of biotin, the bait-PBL fusion protein labels proximal proteins without the activation by a conditional trigger, thereby keeping track of all interactions that occurred over a time period. The ability for selective capture makes the method generally insensitive to protein solubility or protein complexation, with potential applicability for the interactomics studies of membrane proteins and cytoskeletal constituents, providing a major advantage over alternative approaches. Nevertheless, the identity of a candidate interactor does not immediately imply a direct or indirect interaction with the bait but reflects merely proximity [estimated to be ~10 to 15 nm (Kim et al., 2014)]. Furthermore, true interactors are missed (false negatives) if they lack accessible primary amines.

So far PBLs have successfully been used in yeast (Opitz et al., 2017b), protozoa (Opitz et al., 2017a), amoebae (Batsios et al., 2016), embryonic stem cells (Gu et al., 2017), and xenograft tumors (Dingar et al., 2015) to map a wide range of interactomes in both small-scale (i.e. using a single bait protein) and large-scale network mapping approaches (e.g. the protein interaction landscape of the centrosome-cilium interface or the organization of mRNA-associated granules and bodies (mRNP complexes) (Gupta et al., 2015; Youn et al., 2018).

In plants, the number of reports on the use of PBLs is slowly increasing. So far, four papers describe the application of the first generation of PDLs in plants (Conlan et al., 2018; Das et al., 2019; Khan et al., 2018; Lin et al., 2017). In these first trials, overexpression of BioID was combined with long labelling times, very high biotin levels and relatively poor labelling efficiencies. These results suggest that first-generation BioID variants do not achieve sufficient activity in plant tissues due to their temperature-activity profiles.

Recently, two studies evaluated several generations of PBLs in plants, including the third generation TurboID and mTurbo using *N. benthamiana* and Arabidopsis seedlings as model systems and concluded that TurboID outperforms the other PBLs in its capacity of both *cis*-as well as specific *trans*-biotinylation of both known as well as novel target proteins under conditions compatible with normal plant growth (Mair et al., 2019; Zhang et al., 2019).

Here, we expand our current knowledge on the use of PBL as an interactomics tool in plants by performing a systematic survey of different PDL approaches in various plant systems. We provide guidelines for the use of PDL in various frequently used plant models and highlight the most relevant shortcomings and contingencies. Furthermore, we benchmark different PDL methods at the proteomics level by studying the TPLATE protein complex and its interactors using harsh extraction and washing conditions to maximize the removal of false positives. We furthermore employ a strategy which allows the identification of both non-biotinylated as well as biotinylated peptides from a single experiment. Finally, we provide an extensive toolkit to perform PBL *in planta* and foresee that the methods, tools and materials herein will greatly benefit the research community.

## RESULTS

### PBL-mediated biotin labelling efficiency increases upon biotin administration in Solanum lycopersicum

In order to establish PDL in various plant systems, we first tested different PBLs in stable hairy root lines of *Solanum lycopersicum* (see **Figure 1** and **Materials and Methods**). More specifically, we compared the potential applicability of enzyme-catalyzed proximity labelling when using BioID (Kim et al., 2016; Roux et al., 2012), BioID2 (Kim et al., 2016), TurboID or mTurbo (Branon et al., 2018) as PBL. For this, we fused the engineered PBL to FLAG and enhanced green fluorescent protein (eGFP) tags under control of the constitutive cauliflower mosaic virus (CaMV) 35S promoter (**Supplemental Figure 1 and 2**). In all systems tested so far, supplementation of biotin is important for efficient proximity biotin ligation with all the PBLs tested. Plants synthesize biotin endogenously and thus, in certain systems, the intracellular pool of biotin might be high enough for the PBL. In fact, free biotin accumulates in plant mesophyll cells to a high concentration of ca. 11 μM (Alban et al., 2000), while for example in yeast this concentration is more than 10-fold lower (Pirner and Stolz, 2006). Considering that the *K*_*m*_ of BioID for biotin is 0.3 μM, this could, in theory, lead to efficient PDL even in the absence of exogenous biotin supplementation.

**Figure 1.**
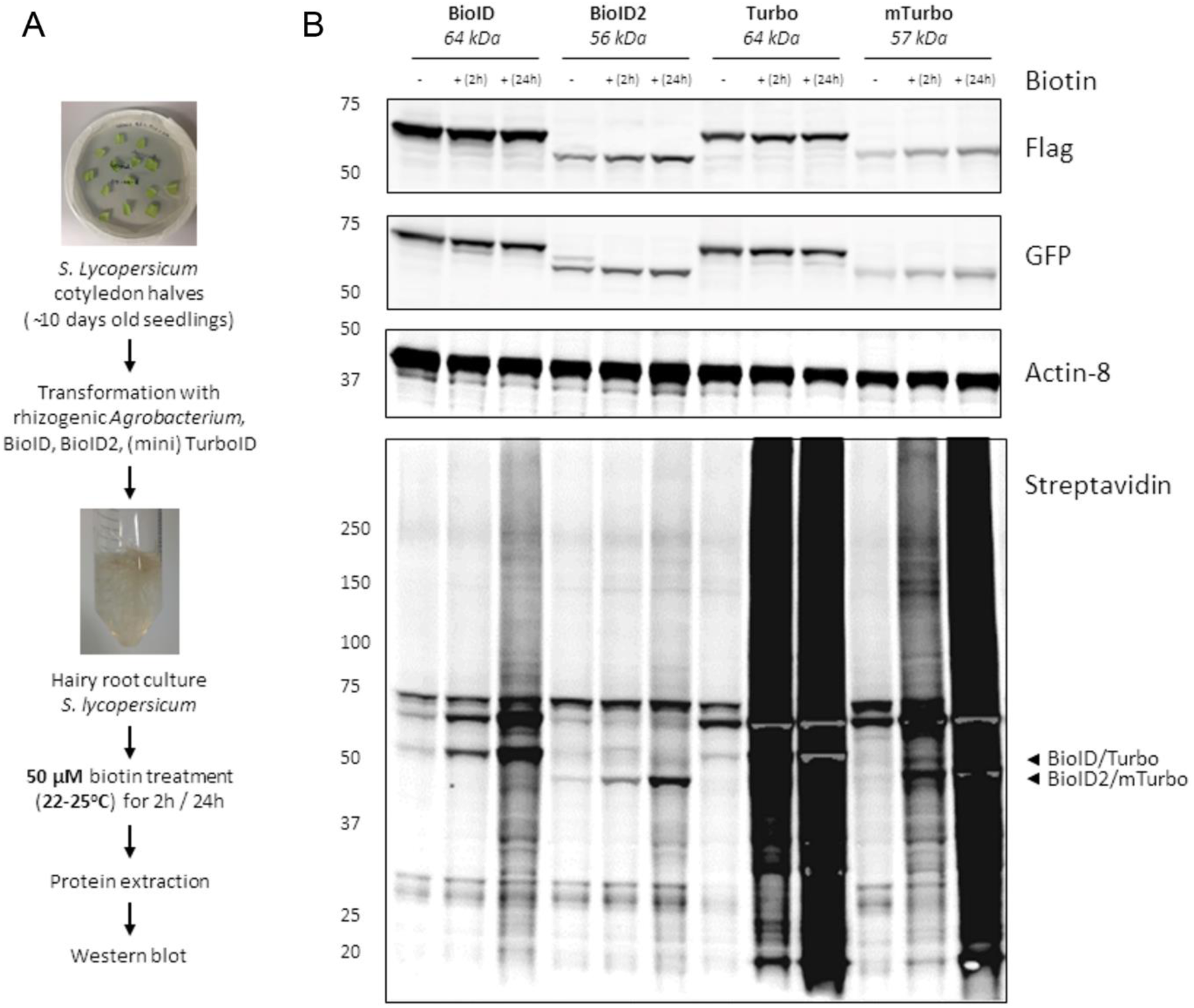
Characterization of enzyme-catalysed proximity labelling in hairy root cultures. Experimental setup. (B) Comparison of biotinylation activity in four PBL-expressing hairy root cultures. Addition of 50 μM exogenous biotin to two-weeks old hairy root cultures for 2 or 24 h was used for labelling. Arrowheads indicate the expected size of the *cis*-biotinylation signal. (B) Comparison of biotinylation activity in four PBL hairy root cultures from wild-type tomato expressing eGFP-BioID-Flag (~66 kDa), eGFP-BioID2-Flag (~56 kDa), eGFP-Turbo-Flag (~64 kDa) and eGFP-miniTurbo-Flag (~57 kDa). Gray regions in intense black areas represent saturation of the streptavidin-s680 signal and is most prominent in case of self-biotinylation activity. This is a representative experiment repeated twice and two independent root cultures were analyzed per combination.

We thus tested biotinylation efficiency in our hairy root system in the presence or absence of biotin using different tagged PBLs as fusion proteins, either codon-optimized for plants or non-codon optimized (**Supplemental Figure 1, Supplemental Table 1 and Supplemental sequences**).

As a test-case for non-bait specific biotinylation, PBL-fused eGFP was used. Biotinylation was evident as smears upon streptavidin-HRP-mediated Western blot detection. This smear depicts biotinylation of other proteins than PBLs, and will be referred to as “*trans*-biotinylation”. As a proxy of PBL activity, we used the *cis*-biotinylation efficiency (i.e. auto- or self-biotinylation level of PBL fusions) as readout (**Figure 1**). Manifold faster kinetics for TurboID and mTurbo over BioID and BioID2 could be observed (**Figure 1**). This is in line with the previously reported lower catalytic activities of the latter PBLs, especially at the growth conditions used (i.e. cultivation of hairy roots was performed at 22−25°C) (Branon et al., 2018). Noteworthy, only residual *trans*-biotinylation was observed when no exogenous biotin was added to the liquid grown hairy root cultures. Therefore, the addition of surplus (free) biotin seems also to function as a trigger of PDL in this system. This observation indicates that PDL in plants (to some extent) might also have the capacity to identify the spatiotemporal dynamics of interactome composition.

### PDL-efficiency depends on growth temperatures and PBL can facilitate trans-biotinylation in Nicotiana benthamiana

We used transient transformation of *Nicotiana benthamiana* leaf mesophyll cells to test the applicability of PDL in a second model system commonly used for protein expression *in planta* under various conditions. In this case, biotin was infiltrated directly into leaf tissue 24 h after transformation and harvested 24 h post-biotin infiltration (**Supplemental Figure 3A**). We confirmed that also in this system, the highest *cis*-biotinylation level was observed in case of TurboID, and supplementation of biotin was important for the efficient detection of *cis*-biotinylation (**Supplemental Figure 3B**). Furthermore, the overall biotinylation output signal in tobacco leaves was higher when biotin concentration was increased from 50 μM to 1 mM (**Supplemental Figure 3B**).

Evaluation of wild-type BirA showed no trans-biotinylation in the presence of 50 μM exogenous biotin (**Supplemental Figure 4A**), confirming that the R118G mutation is responsible for promiscuous labelling in plants. Furthermore, a temperature shift from 22°C to 28°C increased *cis*- and *trans*-biotinylation for both BioID and TurboID, suggesting that temperature control can be used to modulate PDL in plants (**Supplemental Figure 4A and B**, see also below).

Noteworthy, the effect of temperature on TurboID activity was less apparent compared to that of BioID, consistent with the temperature-activity profiles of the two enzymes (Branon et al., 2018). Interestingly, similar to GFP-TurboID expressed in the hairy root cultures, *cis-* biotinylation (**Figure 1**),was saturating already 2 h after biotin administration in *N. benthamiana* (**Supplemental Figure 4D**). TurboID and mTurbo were the only PBLs in plants with biotinylation efficiency occurring in the range of a few hours, as other PBLs did not show any visible sign of *trans*-biotinylation in that time frame **(Figure 1**).

### TurboID can be used for the efficient capture of plasma membrane interactomes in Nicotiana benthamiana

Next, we tested whether we could achieve biotinylation of protein interactors using PDL under the conditions established for *N. benthamiana*. We observed that the bait proteins used in plants for PDL so far were either membrane-anchored and small proteins [HopF2 (Khan et al., 2018) and AvrPto (Conlan et al., 2018)], or nuclear and/or cytoplasmic localized [OsFD2 (Lin et al., 2017), N (Zhang et al., 2019) and FAMA (Mair et al., 2019)].

We therefore tested our conditions for PDL using as test-cases, integral plasma membrane-localized protein complexes with components that reside within a range of a few nm. First, we used a known membrane receptor complex from *Lotus japonicus* comprising two symbiotically active receptor-like kinases (RLK): the LysM-type RLKs NOD FACTOR RECEPTOR 5 (NFR5) and the LRR-RLK SYMBIOTIC RECEPTOR-KINASE (SYMRK). These proteins assemble within the same complex in *L. japonicus* roots (Ried et al., 2014) as well as in *N. benthamiana* upon heterologous expression (Antolin-Llovera et al., 2014). In contrast, the brassinosteroid receptor BRASSINOSTEROID INSENSITIVE 1 (BRI1) did not co-immunoprecipitate with the symbiotic receptor complex indicating no or only weak interactions with these RLKs (Antolin-Llovera et al., 2014). However, using Bimolecular Fluoresce Complementation (BiFC), another study reported some interactions between NFR5 and BRI1 as well as with the *A. thaliana* innate immune pattern recognition receptors FLAGELLIN SENSING 2 (FLS2) (Madsen et al., 2011). To further extend the set of control proteins, we additionally included the EF-TU RECEPTOR (EFR), belonging to the LRR-family, as well as the LOW TEMPERATURE INDUCED PROTEIN LTI6b that is commonly used as a plasma membrane marker in plant cell biology (Grebe et al., 2003).

In a first experiment, we tested whether cytosolic TurboID would non-specifically *trans*-biotinylate the receptors at the plasma membrane. For this, we co-expressed a TurboID-GFP fusion protein with GFP-tagged receptors in *N. benthamiana* and immunoprecipitated (IP) all components using an anti-GFP nanotrap (**Supplemental Figure 5A**). While all co-expressed proteins could be detected before and after the IP, we only detected *cis*-biotinylation of TurboID-GFP but not of the receptors (**Supplemental Figure 5A**). This indicates the absence of non-specific *trans*-biotinylation of membrane resident receptors by a soluble TurboID itself. However, it should be clearly stated that prolonged reaction times and increased expression of TurboID will likely result in a certain degree of non-specificity due to the inherent features of the system.

To test biotinylation between membrane-resident receptors, we co-expressed a NFR5-TurboID (120 kDa) fusion protein with either the known NFR5-interacting RLK SYMRK or with BRI1 and FLS2 that may not be stable components of the NFR5/SYMRK receptor complex. As higher degrees of non-specificity are expected for proteins that reside in close proximity with each other, we tested *trans*-biotinylation 15 and 30 minutes after addition of exogenous biotin (**Figure 2 and Supplemental Figure 5**). As expected, we observed weak *trans*-biotinylation of SYMRK-GFP (150 kDa) by NFR5-TurboID after 15 minutes when SYMRK-GFP was immunoprecipitated using anti-GFP nanotrap beads. With 30 minutes labeling time, stronger *trans*-biotinylation of SYMRK5-GFP was detected (**Figure 2,** upper panel). When applying the same experimental conditions to plants co-expressing BRI1-GFP (157 kDa) and NFR5-TurboID, we detected no *trans*-biotinylation after 15 minutes and only very weak *trans*-biotinylation after 30 minutes of BRI1-GFP. These data show that temporal control during labelling experiments is crucial to maintain specificity in the system, and that BRI1 may reside in close proximity to the NFR5/SYMRK complex, despite a lack of a stable and physical interaction.

**Figure 2.**
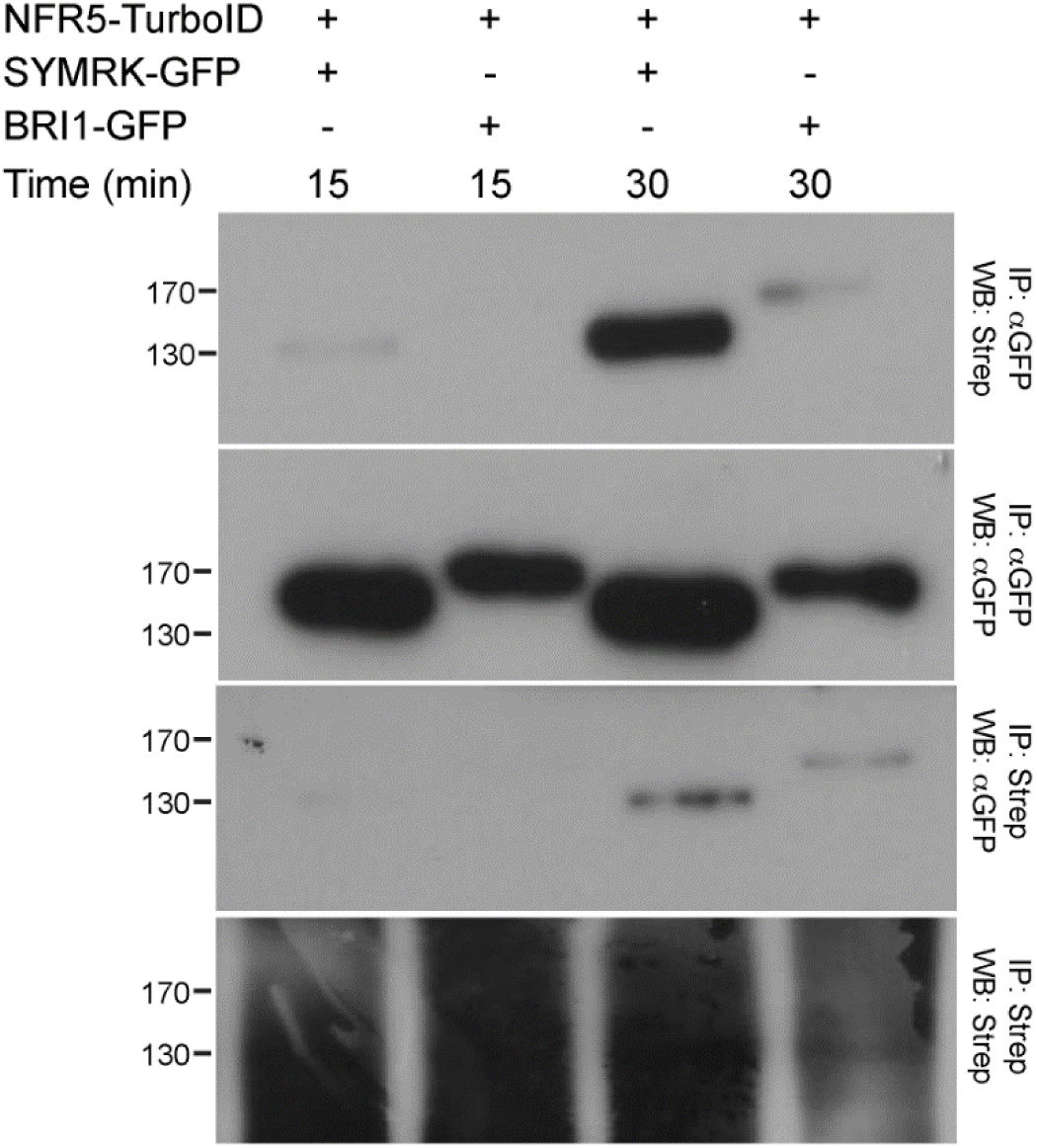
NFR5-TurboID shows strong biotinylation of known interactor SymRK-GFP. Pairwise combination of NFR5-TurboID (120 kDa) with either SYMRK-GFP (150 kDa) or BRI1-GFP (157 kDa) using transient expression in *N. benthamiana* leaves allowed time-dependent and prevalent biotinylation of SYMRK. 50 μM biotin was applied for 15 or 30 min. IP= immunoprecipitation; WB= Western Blot; Strep= Streptavidin.

Given these results, we sought to test a number of other membrane proteins to elucidate whether the observed levels of non-specificity are at least partially dependent on the target protein. We co-expressed NFR5-TurboID with the transmembrane proteins FLS2, EFR and LTI6b. While no *trans*-biotinylation of EFR and LTI6b was detected, we observed a weak signal for BRI1 as shown above as well as for FLS2, but again considerably lower compared to the levels found for SYMRK, indicating an important impact of the target proteins on the *trans*-biotinylation patterns (**Supplemental Figure 5B**). It should be noted that we were not able to detect *cis*-biotinylated NFR5 after immunoprecipitating SYMRK using GFP-nanotraps. This is most likely due to the stringent washing conditions and the possibility that only a fraction of NFR5-TurboID was co-immunoprecipitated together with SYMRK. Taken together, these data are in line with a previously published report (Madsen et al., 2011) and show that predominant *trans*-biotinylation of proximal membrane-resident proteins is possible, even under constitutive expression in heterologous systems. However, stringent control of experimental conditions such as expression levels and exposure time to biotin is greatly advised.

In summary, these data clearly show that TurboID-mediated PDL can be efficiently used for capturing interactors of membrane proteins. Furthermore, it can be advantageous over other methods such as co-immunoprecipitation as it does not require any optimization of the solubilization conditions and provides the possibility to detect transient protein complex constituents.

### Application of PDL in Arabidopsis thaliana cell cultures using the TPLATE complex as a case study

Next, we surveyed the efficiency of *trans*-biotinylation for a stable multi-subunit plant protein complex. As a test case, we selected the plasma membrane-associated octameric TPLATE complex (TPC) (Gadeyne et al., 2014) and used stably transformed *A. thaliana* cell suspension cultures as a third plant model system for PDL.

Given the higher biotinylation level observed in *N. benthamiana* at 28°C (**Supplemental Figure 4**), we started with evaluating different labelling conditions. To study the temperature effect in this system, we grew cells expressing TPLATE-BioID and GFP-BioID, i.e. proteins fused to the first generation PBL, at various temperatures in the presence of 50 μM biotin for 24 h. We subsequently isolated the complex under non-denaturing conditions using streptavidin affinity purification (see **Materials and Methods**), performed tryptic on-bead digest and analyzed the released non-biotinylated peptides using LC-MS/MS.

In order to evaluate the effect of temperature on the biotinylation efficiency and on the subsequent identification of the proteins from the isolated complexes, we focused on the other seven TPLATE complex members. We compared their abundances and fold changes to the control setup (35S::GFP-BioID) after streptavidin purification, taking into account label-free protein quantification (LFQ) intensities (Cox et al., 2014) (**Figure 3A**). In addition to the bait, all seven interacting subunits could be significantly detected at all tested temperatures (**Figure 3B, Supplemental Data Set 1**). The fold changes observed with respect to the control were however not dramatically different between the different temperatures. As we did not observe any major differences with respect to the efficiency of detecting TPC subunits at all tested temperatures, and given the increased efficiency observed in *N. benthamiana* at 28 °C and the likelynegative impact of increased temperature on the physiology of the plants, we opted for 28°C as an optimal trade-off to perform a series of follow-up experiments on the TPC in *A. thaliana* cultures.

**Figure 3.**
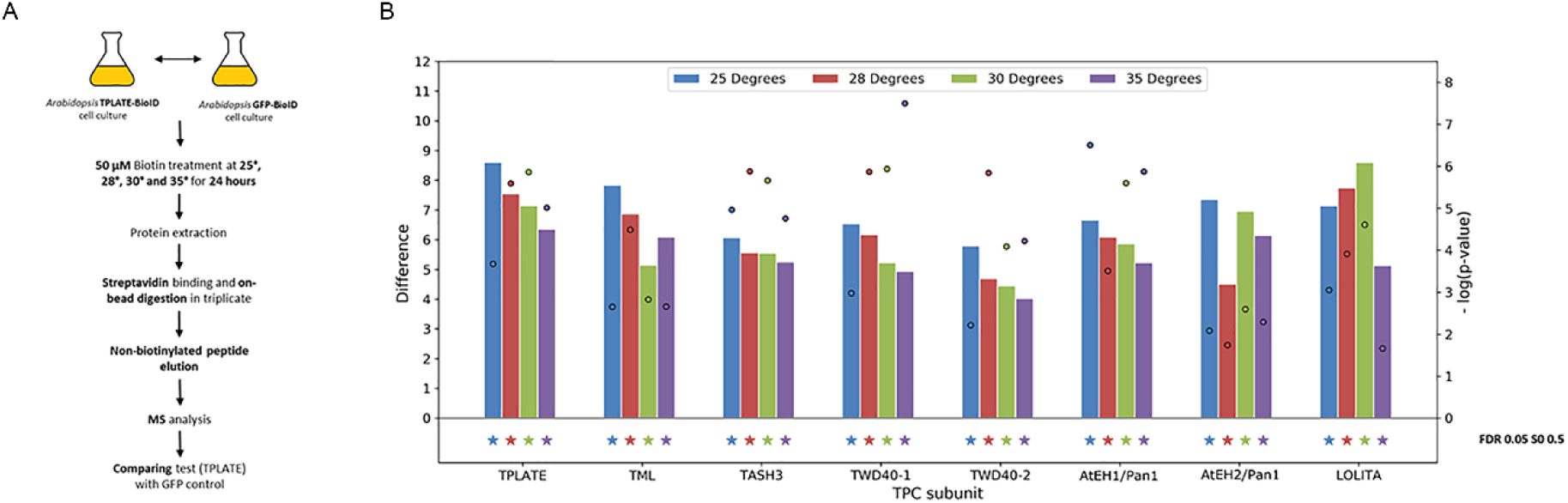
Detection of TPC subunits with TPLATE-BioID is optimal at 28° C. (A) Experimental setup to look for enriched TPC subunits in biotin treated transformed Arabidopsis cell cultures. Cell cultures were incubated with 50 μM biotin at 25°−35°C for 24 h before harvesting. (B) Comparison of the enrichment of the TPC subunits in the TPLATE-BioID samples at different temperatures compared to their respective GFP-BioID controls. Difference (bar charts) and −log(p-values) (dots) are derived from t-tests in Perseus software, using the average LFQ intensities of 3 technical replicates of TPLATE-BioID versus 3 technical replicates of GFP-BioID at similar temperature. All TPC subunits are detected at all 4 temperatures without major differences and all are significantly enriched with TPLATE-BioID (denoted by stars), as determined by permutation based FDR, with cut-offs FDR=0.05 and S0=0.5. The full list of significantly enriched identifications with TPLATE-BioID at all tested temperatures can be found in Supplemental Table 4.

### Various PBLs affect biotinylation of TPC subunits differently

The introduction of a flexible linker (Roux et al., 2012) has been successfully used to extend the labelling radius of PBLs (Kim et al., 2016) (Kim et al., 2016), which is estimated to be about 10 to 15 nm (Kim et al., 2014). This increased labelling radius may be desirable when the protein of interest is significantly larger than the labelling radius of the PBL alone, and/or when the goal is to map the constituency of a larger protein complex or a discrete subcellular region. We thus compared the efficiencies of various PBLs and assessed their biotinylation radius by inserting a 65 aa long flexible linker. *Arabidopsis* cultures expressing C-terminal fusions of TPLATE with BioID or BioID2 were assessed, with and without a 65 aa linker similar to the one that was reported before (Roux et al., 2012). As controls, we generated GFP fused to BioID or BioID2 without additional linker **(Supplemental Figure 6)**.

To test the effect of the linker and to further evaluate the activity of different PBLs in *Arabidopsis* cell culture, transgenic cultures were grown for 24h, with and without exogenous biotin at 28°C, and expression and biotinylation were assessed via Western blotting (**Supplemental Figure 6**). Protein abundance of the BioID and BioID2 constructs was comparable to their respective controls in our cell cultures and was not affected by the addition of biotin. Only TPLATE-BioID2 levels were rather lower. At the level of *cis*- and *trans*- biotinylation, we observed different patterns for each of the fusion proteins used. As several of the detected bands which increased significantly in the presence of biotin, did not correspond to bands in the control or GFP-BioID culture and varied between the different PBLs, they likely represent different *trans*-biotinylated interactors and suggest that the outcome of a BioID-based interaction assay might partially depend on the PBL used. TPLATE-linker PBL showed the most complex biotinylation pattern when comparing to the other setups expressing BioID and BioID2 fusions (**Supplemental Figure 6**), suggesting that the addition of a linker may be used to enhance proximity labelling. Consistent with the results described for tobacco, TurboID constructs showed some residual biotinylation without the addition of exogenous biotin, increased biotinylation after 1 h incubation with biotin and gave rise to an extensive biotinylation pattern after 24 h incubation with biotin in both control and bait cultures, suggesting it is highly promiscuous.

As observed in *N. benthamiana* (**Supplemental Figure 3**) using GFP as bait protein, BioID also outperformed BioID2 using TPLATE as bait in this system, although this might (in part) be skewed due to the lower expression levels of the latter. Adding a flexible linker increased *cis*-biotinylation levels of the bait compared to the constructs without linker (**Supplemental Figure 6A and C**). Overall, our results are consistent with previous observations in non-plant systems suggesting that linkers increase the biotinylation output (Kim et al., 2016).

Following the positive effect of exogenous biotin supplementation (**Supplemental Figures 3 and 4**), we tested the effect of increasing biotin concentrations on *cis*-biotinylation efficiency. Cell cultures expressing TPLATE-linkerBioID were grown at 28°C in the presence of increasing concentrations of biotin (50 μM to 4 mM) for 24 hours and analyzed by Western blotting (**Supplemental Figure 7A**). Supplementing the culture with biotin concentrations in the range of 50 μM to 2 mM increased *cis*-biotinylation output up to ~2-fold. Increasing biotin concentration >2 mM did not further increase *cis*-biotinylation efficiency (**Supplemental Figure 7B**).

We took advantage of the increased biotinylation observed by including a long linker sequence and generated *Arabidopsis* cultures expressing GFP-linkerTurboID and TPLATE-linkerTurboID. Similar to other reports, when sampling was done 24 h post-biotin addition, TurboID efficiency strongly outperformed all other PBLs tested as evident from the high biotinylation levels observed with and without the addition of exogenous biotin for both the control (GFP) as well as the TPLATE expressing cultures (**Supplemental Figure 6B and D**).

In order to compare the different PBL modules, we processed the isolated proteomes of our cell cultures for LC-MS/MS analysis and focused on the relative levels of the various TPC subunits compared to the control setup. Our first mass spectrometry results following streptavidin purification under non-denaturing conditions and on-bead digestion identified all known subunits of the TPC (**Figure 3**). Given that TPC is a robust multi-subunit complex (Gadeyne et al., 2014) and that we identify only non-biotinylated peptides with our on-bead digestion protocol, we assumed that the subunits we detect are a combination of direct biotinylation as well as co-immunoprecipitation of the complex as a whole under the non-denaturing conditions. To test this, we adapted our protocol (**Figure 4A**) and performed protein extraction and stringent washing steps under denaturing conditions using a buffer containing 8M urea and 2% SDS to unfold proteins before streptavidin immunoprecipitation and to remove non-specific, or indirect, non-biotinylated protein binders. We also included the TPLATE-linkerBioID setup treated with 2 mM biotin for 24 h to assess if increased biotin concentration improves TPC subunit detection.

**Figure 4.**
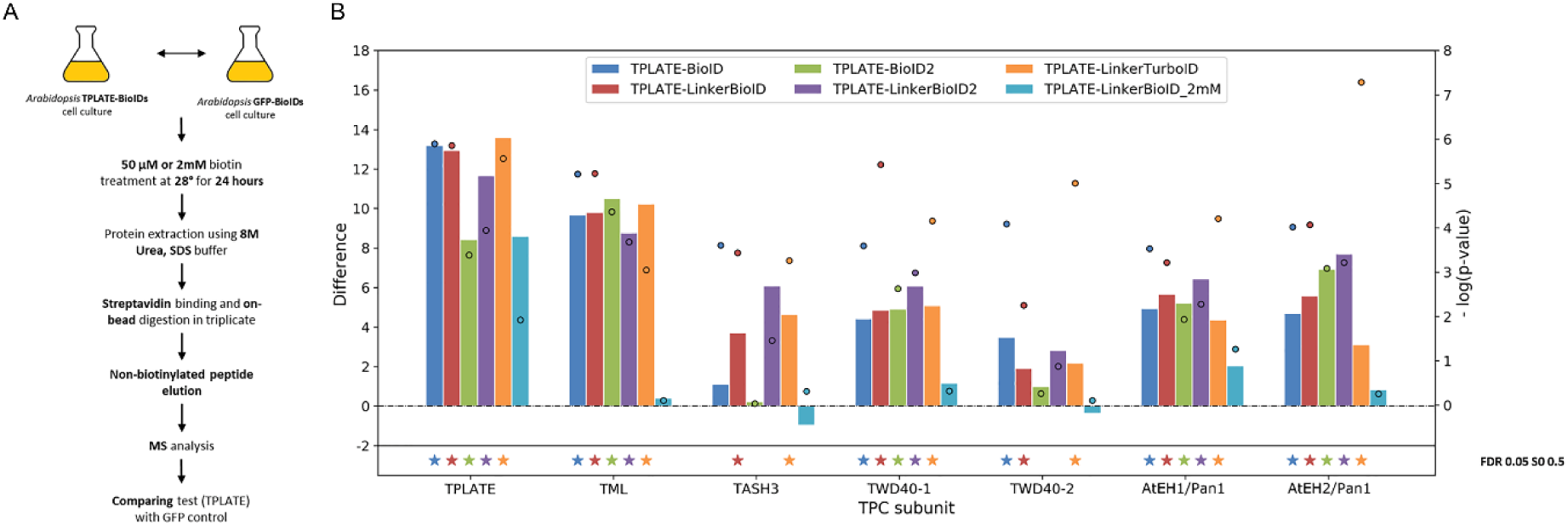
Different TPLATE-PBLs affect biotinylation of TPC subunits differently. (A) Experimental setup. Cell cultures were incubated with 50 μM biotin at 28° C for 24 h before harvesting. Protein extraction was performed under harsch conditions to exclude false postives Comparison of the enrichment of the TPC subunits with different TPLATE-PBLs versus their respective GFP-PBLs at 28° C. Difference (bar charts) and −log(p-value) (dots) are derived from t-tests in Perseus software, using LFQ intensities of 3 technical replicates of the test compared to 3 replicates of the respective control. The stars below the graph denote that proteins were found significantly different to the control by permutation based FDR, with cut-offs FDR = 0.05 and S0 = 0.5. The full list of significantly enriched identifications with different TPLATE PBLs at 28 degrees can be found in Supplemental Table 5.

In agreement with the higher stringency of the isolation procedure, the smallest TPC subunit, LOLITA, which was robustly detected using AP-MS (Gadeyne et al., 2014) and, as shown here, without being denatured before binding to streptavidin beads (**Figure 3**), was no longer detected (**Figure 4B, Supplemental Data Set 2**). LFQ revealed that the remaining seven TPC subunits, including the bait TPLATE, were detectable using BioID, linkerBioID, linkerBioID2 and linkerTurboID, although not all subunits were significantly enriched compared to the GFP PBL control using our statistical threshold criteria (FDR 0.05 and S0 of 0.5). The TASH3 and TWD40-2 subunits, for example, could not be confidently identified with all PBLs. For BioID2, this might be caused by the reduced expression level of the bait in these cultures (**Supplemental Figure 6**), yet this does not explain why this low level of detection is not observed for the other subunits as well (**Figure 4**). We also conclude that adding a long linker increased the robustness of prey identification. For example, using TPLATE-linkerBioID, the TASH3 subunit was detected with 15 peptides compared to only 2 peptides when using TPLATE-BioID (**Supplemental Table 3**). We did not identify TASH3 with TPLATE-BioID2, in contrast to TPLATE-linkerBioID2, where we identified TASH3 with 59 peptides (**Supplemental Table 3**).

Noteworthy, increasing the concentration of biotin from 50 μM to 2 mM adversely affected TPC subunit detection as only the bait itself could be identified. It is likely that increasing biotin concentrations causes residual free biotin to accumulate in the protein extract, even after protein desalting to deplete free biotin, thereby occupying the streptavidin binding sites on the beads which are saturated at >9 μM of biotin. We tested this “saturation hypothesis” using *N. benthamiana* leaves and protein precipitation to completely remove residual biotin, showing that even at low concentration, residual biotin can saturate the streptavidin beads and incapacitate detection (**Supplemental Figure 8**). Hence, special care should be taken to avoid an excess of residual free biotin during streptavidin-based capture. A similar conclusion was obtained in other studies combining PBL with MS analysis *in planta* (Mair et al., 2019; Zhang et al., 2019).

It should be noted that the fold change by which the other TPC subunits were detected with TurboID was comparable or sometimes even lower (e.g. AtEH2/Pan1) compared to the other BioID forms tested (**Figure 4**). This was caused by the fact that TPC subunits were identified with higher abundance in the TurboID control samples, resulting in lower relative fold changes. All individual TPC subunits were detected with more than 20 unique peptides using the GFP-linkerTurboID whereas TWD40-2 was the only TPC subunit detected in the other control GFP-PBLs, which explains its overall low fold change (**Supplemental Table 3**). Nevertheless, TurboID identified most of the TPC subunits more robustly compared to the other PBLs, as evidenced by the overall higher −log10p-values. So, although in our case, TurboID showed to be superior to all others in identifying the other TPC subunits, the lower signal/noise ratio of TurboID, due to its increased activity, might work as a disadvantage to observe differences between bait proteins and control samples, which might even be enhanced if the proteins are targeted to specific subcellular locations.

### The structural composition of protein complexes causes differences in detection between PDL and AP-MS

To further evaluate PDL, we compared the relative levels compared to the bait by which the different TPC subunits were detected using PDL using our stringent washing protocol with a one-step IgG-based pull-down (PD) protocol using the GS^rhino^ tandem affinity purification (TAP) tag (Van Leene et al., 2019). To do this, we used the Maxquant iBAQ value, which is the result of the summed intensity values of the identified peptides, divided by the number of theoretical peptides. We calculated these iBAQ values for each TPC subunit, normalized it to the value for the bait (TPLATE) to correct for differences in bait fusion expression levels, and compared the values of TPLATE-linkerBioID, TPLATE-linkerBioID2 and TPLATE-linkerTurboID with those from PD. When normalized to the bait protein (TPLATE), the other TPC subunits are detected by TurboID at similar levels as compared to PD (**Figure 5A, Supplemental Data Set 3**). The one exception is the subunit LOLITA, which could only be detected by PD. The six other TPC subunits could also be significantly detected by BioID and BioID2, however with less efficiency.

**Figure 5.**
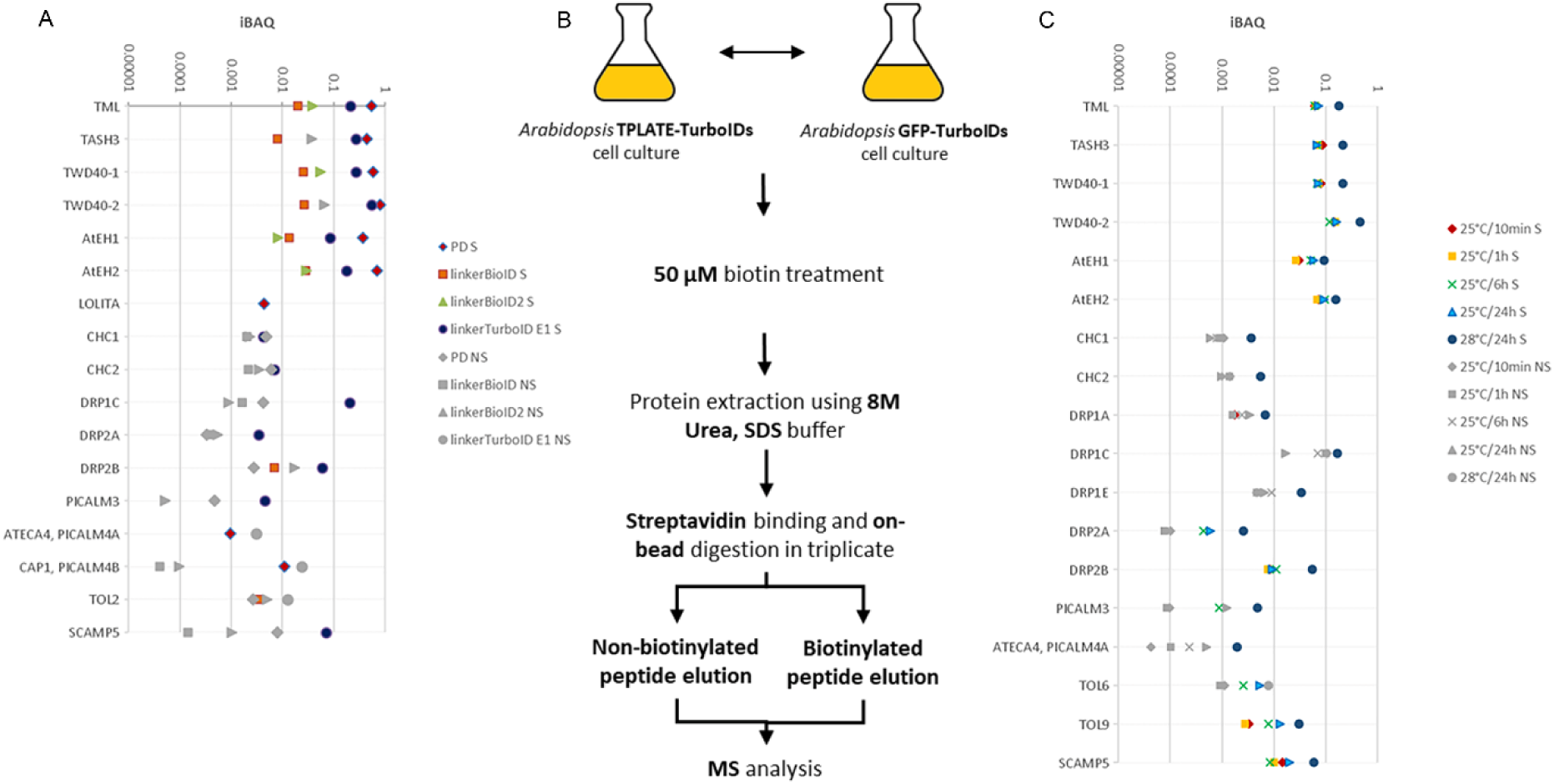
Comparing identification of a subset of proteins co-purified with TPLATE using GSrhino pull down (PD), linkerBioID, LinkerBioID2 or LinkerTurboID. (A) Pull down and proximity biotinylation comparison of a selection of TPLATE interactors. Experiments were performed in triplicate, using TPLATE as bait. Per set of experiments, MaxQuant iBAQ values, which are the summed intensity values divided by the number of theoretical peptides were calculated and normalized versus the bait in order to compare the relative abundance of the proteins between the four different approaches. Proteins that were identified significantly (S) in either method are represented with a colored shape. Proteins that were identified below the significance threshold (NS) for a given method are indicated with grey shapes. (B) Schematic overview of the experimental setup to detect biotinylated and non-biotinylated peptides. Following on-bead digestion, non-biotinylated and biotinylated peptides were separately analyzed using sequential elutions and all identified peptides were used for MS analysis. (C) Overview of a subset of the identified interactors, color coded according to their statistical significance in the different experiments (S=Significant, NS=Not significant) by combining MS data from both elution fractions. Arabidopsis cell cultures expressing TPLATE-linkerTurboID were grown at 25 degrees and supplemented with exogenous biotin for 10min, 6hours or 24hrs. Results were compared to the experiment from panel A where the culture was grown at 28 degrees in the presence of biotin for 24hrs. The complete list of significantly enriched identifications of the experiments shown in panel A and C, including their normalized average iBAQ values, can be found in Supplemental Table 6 and 7.

The fact that the smallest subunit, LOLITA, could only be identified via AP-MS, indicates that this subunit is not biotinylated although it harbors 11 lysine residues, possibly reflecting the structural composition of the TPC. Our results furthermore reveal that, except for LOLITA, all TPC subunits, which are part of a protein complex in the range of 1MDa can be identified using our stringent wash protocol as a proxy for biotinylation.

### TurboID allows broadening the interactome of protein complexes

We subsequently broadened the analysis towards other interactors and compared all proteins that were significantly enriched in one of the datasets (TPLATE-GS^rhino^, TPLATE-linkerBioID, TPLATE-linkerBioID2 and TPLATE-linkerTurboID) (**Supplemental Table 4A**). Whereas the overall number of significant interactors identified with the GS^rhino^ and linkerBioID tags was higher than the number of significant interactors found with linkerTurboID, the latter identified several known players in clathrin-mediated endocytosis (CME) with much stronger statistical significance (**Figure 5A**). These players included the two Clathrin Heavy Chains (CHC), and several Dynamin Related Proteins (DRP). Moreover, TPLATE-linkerTurboID allowed to significantly enrich for novel interactors with a clear link to CME such as the Secretory Carrier Membrane Protein 5 (SCAMP5) and an ANTH/ENTH protein, PICALM3. Integral membrane SCAMP proteins are hypothesized to act in both the exo- and endocytic pathways between the PM and TGN (Law et al., 2012). PICALM3 (Phosphatidylinositol binding clathrin assembly protein) was not identified before as a TPC iteractor, but PICALM4A (AtECA4) and 4B (CAP1), were previously found associated with TPC (Gadeyne et al., 2014) and also confirmed here using our PD approach (**Figure 5A**).

### Identification of biotinylated peptides enhances the identification power of PDL and allows identifying structural relationships between complex subunits

The interaction between biotin-streptavidin is strong enough to be maintained even under harsh conditions (**Supplemental Figure 8**). Thus, biotinylated peptides are expected to be retained on the streptavidin beads. Following stringent washing under denaturing conditions, on-bead digest will release non-biotinylated proteins, which can subsequently be identified using LC-MS/MS. This approach, however, does not provide direct evidence for biotinylation and it relies on the assumption that only biotinylated proteins remain bound to the beads after the washing steps. To acquire direct proof of biotinylation, and to further enhance the power of PDL to identify interactors, release of biotinylated peptides from the Streptavidin beads and their subsequent MS-based identification is required.

Thus, we expanded the protocol (**Figure 5B**) to also be able to identify biotinylated peptides. For this, we included a second elution step (see **Materials and Methods**) to release the biotinylated peptides from the beads using an adapted protocol based on previous work (Schiapparelli et al., 2014). This approach enables the detection of both non-biotinylated as well as biotinylated peptides in the same experimental setup.

As a previous report on TurboID describes no major changes in the activity of TurboID between 22 and 30°C and used biotin treatments of only a few hours (Mair et al., 2019), we tested whether we could improve the identification of novel TPC interactors by reducing the time of biotin addition to our cell cultures grown at normal growth temperatures. We, therefore, performed a series of experiments comparing short (10min and 1h), medium (6 h) and long (24 h) biotin treatments at the normal growth temperature (25°C) of our *Arabidopsis* cell culture. We compared the iBAQ values of all significant hits, using both elutions of each experiment at 25°C with those from our 24hrs experiment at 28°C (**Figure 5B** and **Supplemental Table 4B**). The robustness of detecting interactors clearly increased with longer biotin incubation times. Also, there was a positive effect of working at a slightly elevated temperature (**Figure 5C**). Combining both elution fractions also increased the robustness of interactor identification. More specifically, including the second elution allowed the identification of additional DRPs, AtECA4 as well as TOL6 and TOL9 (**Figure 5C**), compared to the results when only the first elution (on-bead digestion) was analysed (**Figure 5A**).

Out of the five TOL proteins studied so far, TOL6 and TOL9 localize strongly at the plasma membrane (Moulinier-Anzola et al., 2020). TOL proteins are part of the endosomal sorting complexes required for transport (ESCRT) pathway and act as gatekeepers for degradative protein sorting (Korbei et al., 2013). We confirmed the association between TPLATE and TOL6, TOL9 and SCAMP5. TOL6-Venus revealed a high degree of colocalization with TPLATE-TagRFP at endocytic foci on the PM (**Figure 6A**), which was severely reduced when the image of one channel was flipped horizontally (**Figure 6B**). Furthermore, quantitative analysis showed TPLATE interacting with TOL9 and SCAMP5 by ratiometric BiFC. The YFP/RFP ratio was significantly higher for all four independent combinations tested compared to a control set where we combined TPLATE with the shaggy-like kinase BIN2 (**Figure 6C** to **6H**). The identification and confirmation of these novel interactors shows that PDL can expand our knowledge on the interactomes of multisubunit complexes in plants beyond currently used AP-MS-based approaches.

**Figure 6.**
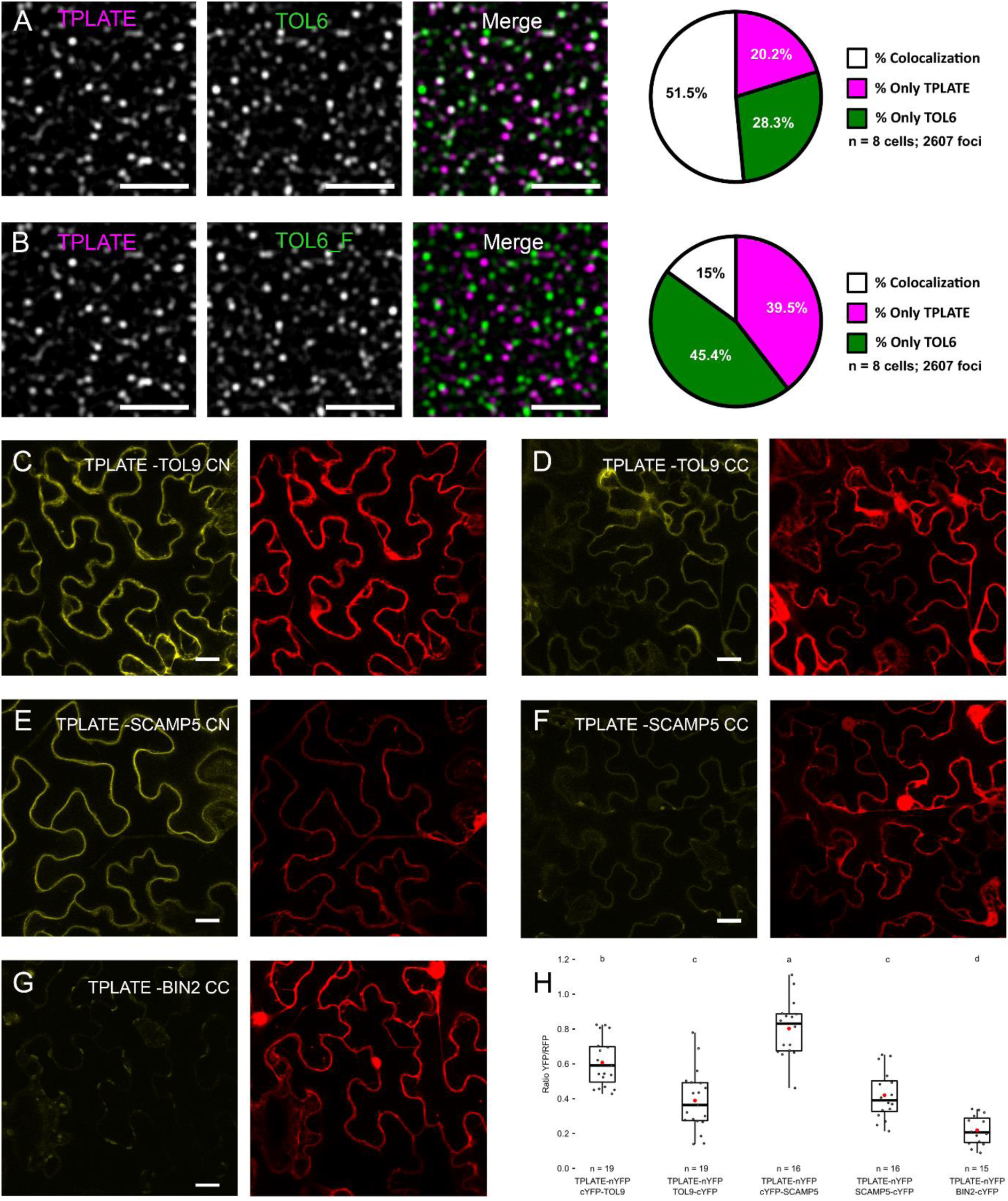
TOL6, TOL9 and SCAMP5 can be confirmed as novel TPC interactors. (A and B) Representative spinning disc dual-color images and corresponding quantification of colocalization (%) between TPLATE and TOL6. TPLATE-TagRFP endocytic foci at the PM were compared with TOL6-Venus foci (A) as well as horizontally flipped TOL6-Venus (TOL6_F) channel images as control (B). Eight movies from three individual plants, and in total 2607 foci were analyzed. (C to H) Ratiometric BiFC analysis confirming the interaction of TOL9 (C and D) and SCAMP5 (E and F) with TPLATE. BIN2 (G) was used as a control. CC and CN refer to the orientation of the nYFP adn cYFP, N-terminal cYFP is CN and C-terminal cYFP is annotated as CC. (H) Box plot and Jitter box representation of the quantification of the YFP/RFP fluorescence ratios (n ≥ 15). The black line represent the median and the diamonds represent the mean. Letters above the plots indicate statistical significance using a Welch-corrected ANOVA to account for heteroscedasticity.Scale bars represent 5 μm (A and B) or 20 μm (C to G).

Next to enhancing the robustness of TurboID to identify interactors, the identification of biotinylated peptides also provides direct proof of the proximity of specific domains of the prey proteins with respect to the bait. We, therefore, tested whether biotinylated peptides could reveal differential proximity between specific domains of TPC subunits using the TPLATE-linkerTurboID as bait (**Figure 7** and **Supplemental Data Set 4**). The highest number of biotinylated peptides were identified for TPLATE (44 biotinylated peptides), followed by TWD40-1 (18), AtEH2/Pan1 (16), AtEH1/Pan1 (12), TWD40-2 (9) and TML (3). No biotinylated peptides could be detected for LOLITA, correlating with our previous results. Mapping non-biotinylated and biotinylated peptides, taking into account their relative abundance, on the different TPC subunits revealed differences in the number of detected peptides as well as differences in the distribution of the biotinylated peptides along the length of the subunits. Whereas the bait, TPLATE, shows a relatively even distribution of biotinylated peptides along the protein sequence, there is a clear tendency of the AtEH1/Pan1, AtEH2/Pan1 and TML subunits towards increased biotinylation at their C-terminal parts **(**Figure 7**)**. It is tempting to speculate that the observed distribution of biotinylated peptides, as well as their absence, reflect the proximity of the domains as well as structural constraints with respect to the bait protein and that proximity biotinylation, next to providing topology information in case of transmembrane proteins (Kim et al., 2018), also harnesses the potential to help deduce structural insight into protein complexes.

**Figure 7.**
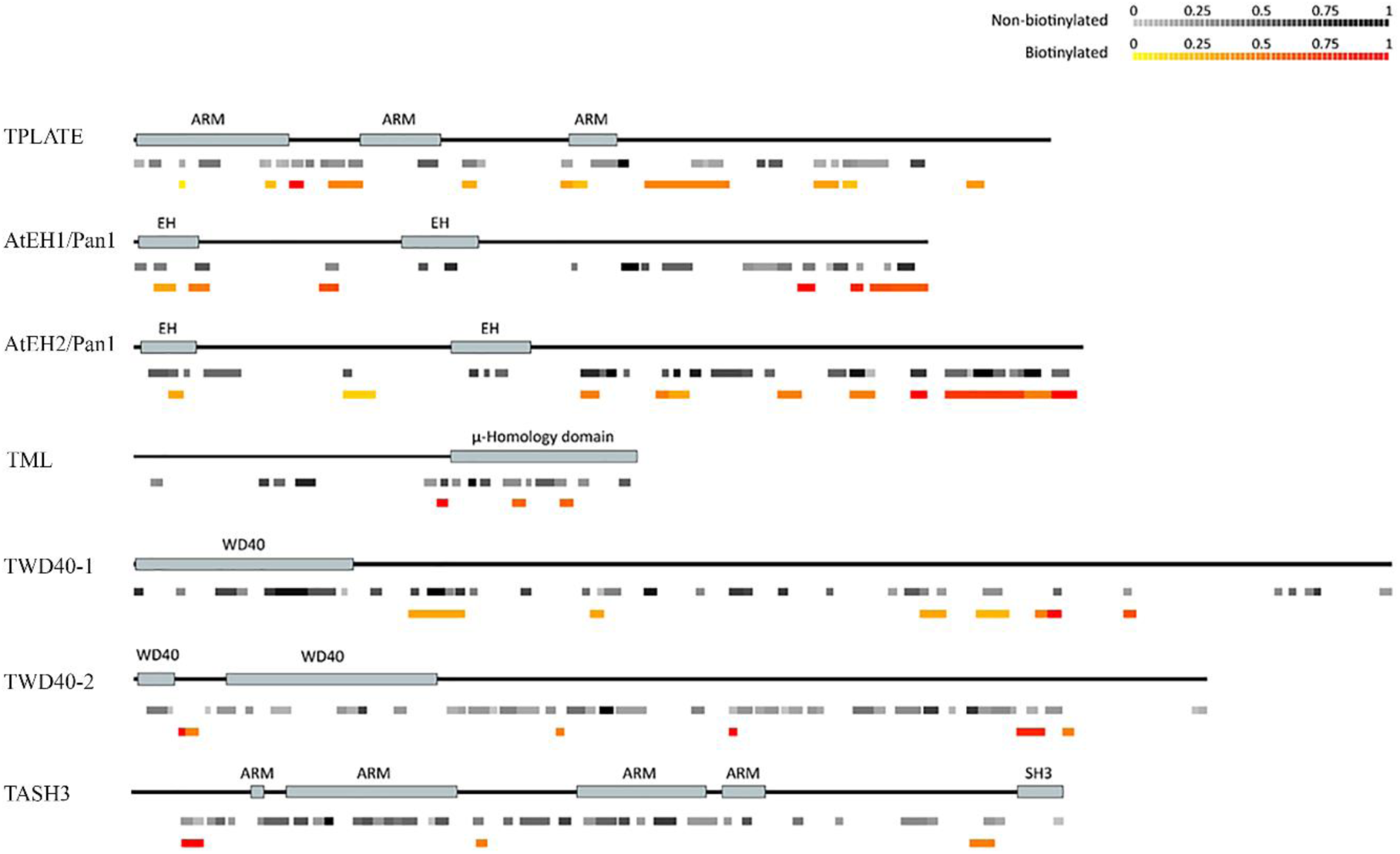
Mapping of biotinylated versus non-biotinylated peptides reveals differential proximity/accessibility of specific TPC subunit domains. Schematic representation of seven TPC subunits and their domains. Identified peptides, color-coded according to their abundance (in grey for non-biotinylated peptides and from yellow to red for biotinylated peptides), are mapped onto them.

## DISCUSSION

We provide a comprehensive comparison of various PBL based proximity labelling strategies in plants. We show that TurboID is the most promiscuous PBL, and that this sometimes leads to a lower signal to noise ratio. We also provide guidelines and approaches for interactome capture in various plant systems specifically focusing on proteins that are intrinsic or peripheral to the plasma membrane. Furthermore, we show that for each bait/system conditions might benefit from independent optimization.

We observed that in all three plant systems tested, the exogenous application of biotin enhances PDL output but might not be a strict requirement for the successful application of PDL. This result seems to contradict with what has been reported for a related method called INTACT (isolation of nuclei tagged in specific cell types) in plants. This method allows for affinity-based isolation of nuclei from individual cell types of tissue. INTACT relies on the endogenous pool of biotin as no exogenous supplementation is required (Deal and Henikoff, 2011). In INTACT, nuclei are affinity-labelled through transgenic expression of the wild-type variant of BirA which biotinylates a nuclear envelope protein carrying biotin ligase recognition peptide from ACC1. This tag acts as a native substrate for the *E. coli* biotin ligase BirA (Beckett et al., 1999). The use of wild-type BirA along with its preferable substrate could explain the higher affinity for the free biotin pool in INTACT, and the peptide used as fusion is an optimal substrate for the bioAMP intermediate. We assume that various proteins may show variability in functioning as acceptors of bioAMP (e.g. depending on the accessibility of lysine residues).

PDL utilizing bacterial enzymes poses the question of whether these enzymes could perform adequately in plants (Kim et al., 2016). The activity optimum for BioID2 is 50ºC, whereas for BioID this is 37ºC and thus BioID2 may be most adequate for use at higher temperature conditions. Both temperatures are however far-off from the usual growth temperatures of most plant species grown in temperate regions (e.g. several Arabidopsis sp.). Both BioID2 and BioID show reduced activity below 37ºC [(Kim et al., 2016) and our results herein]. Furthermore, the lower temperature optimum of TurboID (and mTurbo) (Branon et al., 2018) would imply that TurboID may function better at normal plant growth temperature. Previous work showed no enhanced activity of TurboID when using temperatures above normal plant growth conditions (Mair et al., 2019). We observed however that TurboID activity increases around 2-fold from 22°C to 28°C and that there is a beneficial effect of slightly increasing the growth temperature of our cell cultures on the identification of specific interactors of TPC. At all tested temperatures, we observed that TurboID (and mTurbo) outperforms other PBLs in terms of speed and promiscuity. Hence, TurboID might be preferred over other pBLs when it concerns the initial study of (transient) complex composition where the generation of as much as possible specific biotinylation output in a short time might be desirable.

However, the strong promiscuity of the control might also work as a disadvantage in revealing specific interactions in cases where the reaction cannot be controlled that easily in time or when both the bait and the control would be targeted to a confined intracellular space. Furthermore, controls may express at high levels and show increased diffusion due to their smaller hydrodynamic radius, further skewing results.

We provide evidence that our methods and conditions apply to plasma-membrane complexes. We showed that the interaction of the symbiotic RLKs NFR5 and SYMRK can be identified by exploiting PDL and particularly the PBL TurboID. Furthermore, the use of proper negative controls is imperative. However, even though the brassinosteroid receptor BRI1 was not co-immunoprecipitated with the symbiotic receptors in a previously published dataset (Antolin-Llovera et al., 2014), we detected weak biotinylation of this RLK and the immune-receptor FLS2. While it could be interpreted as unspecificity within the PBL system, it should also be considered, that PBL allows labelling of transient interactions or proximal proteins. As a consequence, continuous unstable interactions accumulate to detectable amounts of proteins and would thus allow their identification. As PDL using TurboID is capable of *trans*-biotinylation in the range of minutes (15 min under our experimental conditions), the enrichment of unstable interactions would thus be more prominent. Therefore, putative interactions identified by PBL still need to be verified using independent experimental systems but comparisons between the different experimental systems should always reflect the technical limitations of each approach.

By expanding our protocols and PBLs into *Arabidopsis* cell cultures, we could not only reproduce the composition of the TPC except for one subunit, but we could also robustly identify and confirm other CME players and novel interactors using the third generation PBL. We show that MS-based identification of interactors is more robust using prolonged biotin exposure of Arabidopsis cell cultures and that the use of linkers can be advantageous when it comes to identifying protein-protein interactions of multi-subunit complexes. Furthermore, TPLATE-linkerBioID2 shows reduced *cis*-biotinylation compared to TPLATE-linkerBioID in the presence of exogenous biotin but seems to function in the absence of biotin suggesting that in plants, BioID2 can function in tissues where exogenous supplementation of biotin may be less effective, e.g. the vasculature. Furthermore, increased biotin application can lead to serious impediments when it comes to the identification of interactors as this can interfere with biotinylated proteins binding on streptavidin slurries. Caution is warranted to assure sufficient capture-capacity of biotinylated proteins since the amount of beads needed for capture should be tested for each experimental model system/setup/protocol.

Complementary to the reports on TurboID *in planta* published so far (Mair et al., 2019; Zhang et al., 2019), we have established a strategy that uses much harsher conditions, with higher concentrations of SDS and urea for extraction and washing to remove as much as possible false positives (i.e. non-biotinylated proteins). Finally, we also provide a protocol for the simultaneous identification of biotinylated and non-biotinylated peptides. This approach allowed us to increase the robustness of interactor identification and provided evidence for the accessibility of different protein domains to PDL. We show that AtEH1/Pan1, AtEH2/Pan1 and TML subunits are preferentially biotinylated at their C-terminal parts, suggesting that their C-termini are in closer proximity to the C-terminal end of TPLATE and/or some domains (even complex subunits such as LOLITA) are not accessible for biotinylation. We thus provide evidence that PDL approaches in plants, combined with harsh extraction/washing conditions may be able to provide structural information of multi-subunit protein complexes and that this may be extended to the topology of membrane proteins.

Our results are complementary to the work deposited in BioRxiv reporting the use of TurboID to identify transient signalling components (Kim et al., 2019) and novel regulators of plant immunity (Zhang et al., 2019), as well as for the efficient capturing of cell- and subcellular compartment-specific interactomes (Mair et al., 2019). Taken together, these four studies provide a new arena for the identification of novel protein-protein interactions in plants.

## MATERIAL AND METHODS

### Bacterial strains

For cloning, *Escherichia coli* strains DH5α, DH10B or Top10 were used using standard chemical transformation protocols. Electrocompetent *Agrobacterium tumefaciens* C58C1 Rif^R^ (pMP90), AGL1 Rif^R^ or GV3101 Rif^R^ bacterial cells (i.e. a cured nopaline strain commonly used for tobacco infiltration (Ashby et al., 1988) were used for tobacco infiltration as well as Arabidopsis cell culture transformation. Electrocompetent rhizogenic *Agrobacterium* (RAB) ATCC15834 (ATCC® 15834™)(Kajala et al., 2014) bacterial cells were used for hairy root transformation.

### Cloning of constructs

For constructs used in hairy roots: Constructs encoding the full-length ORF of the PBL (e.g. BioID (pDEST-pcDNA5-BioID-Flag C-term, a kind gift from the Gingras laboratory (Couzens, Knight et al. 2013)), BioID2 (MCS-BioID2-HA, Addgene, Plasmid #74224 (Kim, Jensen et al. 2016)), TurboID (V5-TurboID-NES_pCDNA3, Addgene, Plasmid #107169 (Branon, Bosch et al. 2018)), mTurbo (V5-miniTurbo-NES_pCDNA3, Addgene, Plasmid #107170 (Branon et al., 2018) were PCR amplified using Q5® High-Fidelity DNA Polymerase (New England Biolabs, Cat n° M0491) with oligonucleotide primers containing attB recombination sequences. The forward and reverse primer additionally encoded the GGGGS linker and the Flag-tag (DYKDDDDK) followed by a stop codon, respectively. The primer sequences are depicted in **Table S2**. The resultant attB-flanked PCR products were used in a Gateway® BP recombination reaction with the pDONR™ P2r-P3 vector (Life Technologies, Carlsbad, CA, USA) according to the manufacturer’s instructions, thereby creating an entry clone. The construct was transformed in DH5α chemical competent cells and verified by sequencing (i.e. Sanger sequencing). Using a standard multisite (3-fragment) Gateway® LR cloning strategy as described by (Van Leene et al., 2007), the entry clones together with pEN-L1-F-L2 encoding eGFP (Karimi et al., 2007a) (https://gateway.psb.ugent.be/search) and pEN-L4-2-R1 encoding the constitutive cauliflower mosaic virus (CaMV) 35S promoter (Karimi et al., 2007a), were recombined with the multisite Gateway destination vector pKm43GW (Karimi et al., 2007a) to generate expression constructs. More specifically, the multisite LR Gateway reaction resulted in translational fusions between the eGFP and the proximity labels, driven by the 35S promoter. This way, the following expression constructs were created; Pro35S::eGFP-BioID, Pro35S::eGFP-BioID2, Pro35S::eGFP-TurboID and Pro35S::eGFP-miniTurbo and Pro35S::eGFP-BioID construct (in pKm43GW), with a C-terminally triple HA-tagged BioID fused to eGFP.

For constructs used in *N. benthamiana*: original BioID, BioID2 and TurboID DNA sequences were taken from (Branon et al., 2018; Kim et al., 2014; Roux et al., 2012), codon-optimized to *Arabidopsis*. The GOLDENGATE compatible BirA, BioID, BioID2 and TurboID were synthesized and codon-optimized using the codon optimization tool of Integrated DNA Technologies, Inc. The ORFs were synthesized with BsaI overhands and were ligated to the Level1/2 vector pICSL86900 and pICSL86922, as previously described (Patron et al., 2015). The following expression vectors were used: Pro35S::BirA-Myc, Pro35S::BioID-myc, Pro35S::HF-BioID2-HA and Pro35S::superfolderGFP-TurboID-FLAG.

The genomic sequence of NFR5 and the coding sequence of BRI1 was synthesized with BsaI overhangs for Golden Gate as Level1 vector (Binder et al., 2014). Pro35S::NFR5-TurboID and Pro35S::BRI1-GFP were created by Golden Gate cloning in Xpre2-S (pCAMBIA) vectors (Binder et al 2014). Pro35S::FLS2-GFP was kindly provided by Hemsley lab, University of Dundee, Scotland. Pro35S::EFR-GFP (Schwessinger et al., 2011) and Pro35S::SymRK-GFP/ Pro35S::NFR5-GFP (Madsen et al., 2011; Wong et al., 2019) were kindly provided by Cyril Zipfel (University of Zurich, Switzerland) and Jens Stougaard (Aarhus University, Denmark).

BiFC constructs were created in the 2in1 BiFC vectors (Grefen and Blatt, 2012). The entry clones were generated by a Gateway® BP recombination reaction using coding sequences of SCAMP5 and TOL9 (BioXP/gBlocks, IDT). TPLATE was amplified from the pDONR plasmid described before (Gadeyne et al., 2014). All entry clones were sequence verified. The BIN2 entry plasmid was kindly provided by Jenny Russinova (Houbaert et al., 2018). Entry clones were combined in a Gateway® LR recombination reaction with an empty BiFC destination vector and selected using LB containing spectinomycin and Xgal. Final BiFC vectors were checked by restriction digest and sequencing of the recombination borders.

For constructs used in *A. thaliana*: BioID and BioID2 DNA sequences were taken from (Kim et al., 2014; Roux et al., 2012), codon-optimized for *Arabidopsis* using the codon optimization tool of Integrated DNA Technologies, Inc. The BioID and BioID2 with and without linker (GGGGS)_13_ with stop codon, flanked by attB2 and attB3 sites (Karimi et al., 2005) were synthesized by Gen9 in the Gm9-2 plasmid. The TurboID sequence (Tess et al., 2018) was codon-optimized to *Arabidopsis* using the codon optimization tool of Integrated DNA Technologies, Inc. TurboID with linker (GGGGS)_13_ with stop codons, flanked by attB2 and attB3 sites (Karimi et al., 2005), was synthesized by GenScript in the pUC57 plasmid. Entry clones of eGFP (Mylle et al., 2013), and TPLATE (At3g01780) (Van Damme et al., 2006) without stop codon were used in a triple Gateway LR reaction, combining pK7m34GW or pH7m34GW (Karimi et al., 2005), pDONRP4-P1R-Pro35 and pDONRP2-P3R-BioID/BioID2/(GGGGS)_13_BioID/(GGGGS)_13_ BioID2/(GGGGS)_13_ TurboID to yield pK7m34GW, Pro35S::GFP/TPLATE-BioID, pK7m34GW, Pro35S::GFP, pH7m34GW, Pro35S::TPLATE-BioID2, pK7m34GW, Pro35S∷TPLATE-(GGGGS)_13_ BioID/BioID2 and pK7m34GW, Pro35S∷GFP/TPLATE-(GGGGS)_13_ TurboID. Sequences of these constructs can be found in supplementary data.

ProTOL6p∷TOL6:Ven was obtained by replacing mCherry in ProTOL6∷TOL6:mCherry (Korbei et al., 2013) with the Venus-tag (Ven), which was PCR amplified with the primer pair: NotImcherryu/NotImcherryd from proPIN2∷PIN2:VEN (Leitner et al., 2012).

### Plant transformations

Hairy roots: Seeds of tomato (*Solanum* spp.) cv. Moneymaker were surface-sterilized in 70% ethanol for 10 min and in 3% NaOCl for 20 min (rinsing with sterile deionized water was performed in between the two sterilization steps), and then rinsed 3 times 5 min each with sterile deionized water. The seeds were germinated on Murashige and Skoog (MS) tissue culture medium (Murashige and Skoog, 1962) containing 4.3 g/L MS medium (Duchefa; catalog no. M0221.0050), 0.5 g/L MES, 20 g/L sucrose, pH 5.8, and 8 g/L agar (Difco; catalog no. 214530) in magenta boxes (~50 ml). The pH of the medium was adjusted to 5.8 with KOH and autoclaved at 121°C for 20 min. The boxes were covered and placed in the dark at 4°C in a cold room for two days. Subsequently, the boxes were transferred to a 24°C growth chamber (16 h light/8 h photoperiod) for ~10 days until cotyledons were fully expanded and the true leaves just emerged. Rhizogenic *Agrobacterium* (RAB) transformation was essentially performed as described previously (Harvey et al., 2008) with some minor modifications. More specifically, competent rhizogenic *Agrobacterium* cells were transformed by electroporation (Shen and Forde 1989) with the desired binary vector, plated on YEB medium plates with the appropriate antibiotics (100 mg/L spectinomycin), and incubated for 3 to 4 d at 28°C. A transformed rhizogenic *Agrobacterium* culture was inoculated from fresh plates into YEB liquid medium with the appropriate antibiotics added and grown overnight at 28°C with shaking at 200 rpm. The RAB culture was used to transform 20 to 40 tomato cotyledon halves. Using a scalpel, the cotyledons were cut in half from ~10 days old tomato seedlings, transferred (adaxial side down) onto MS liquid medium. The MS liquid was subsequently removed and the cotyledon halves immediately immersed in a bacterial suspension at an optical density at 600 nm of 0.3 in MS liquid medium for 20 min, then blotted on sterile Whatman filter paper and transferred (adaxial side down) onto MS agar plates without antibiotics (4.3 g/L MS medium, 0.5 g/L MES, 30 g/L sucrose, pH 5.8, and 8 g/L agar). The co-cultivation culture plates were sealed with aeropore tape. After 3 to 4 days of incubation at 22−25°C in the dark (Oberpichler, Rosen et al. 2008), the cotyledons were transferred to MS agar plates with 200 mg/L cefotaxime (Duchefa; catalogue no. c0111.0025) and 50 mg/L kanamycin and returned to 22−25°C. Typically, three to five independent roots arise from each cotyledon. The expression of the eGFP marker of antibiotic-resistant roots that emerged was monitored by using fluorescent microscopic imaging (Leica stereomicroscope and imaging DFC7000 T Leica microscope camera) and four to ten independent roots showing expression of the marker were subcloned for each construct. These roots were subsequently transferred to new selection plates with the same antibiotic concentration for 3 rounds of subcultivation (~6 weeks) before antibiotics-free cultivation of the hairy root cultures in liquid MS (in 50 ml Falcon tubes containing 10 to 30 ml MS medium at 22−25°C and shaking at 300 rpm) and downstream analysis. After 3 rounds of cultivation, root cultures were maintained and grown in antibiotics-free half-strength (½) MS medium supplemented with 3% sucrose at 22−25°C.

#### N. benthamiana

Wild-type tobacco (*Nicotiana benthamiana*) plants were grown under normal light and dark regime at 25°C and 70% relative humidity. 3-to 4-weeks old *N. benthamiana* plants were watered from the bottom ~2h prior infiltration. Transformed *Agrobacterium tumefaciens* strain C58C1 Rif^R^ (pMP90), AGL1 Rif^R^) or GV3101 Rif^R^ harbouring the constructs of interest were used to infiltrate tobacco leaves and used for transient expression of binary constructs by *Agrobacterium tumefaciens*-mediated transient transformation of lower epidermal leaf cells essentially as described previously (Boruc et al., 2010). Transformed *Agrobacterium tumefaciens* were grown for ~20h in a shaking incubator (200 rpm) at 28°C in 5 mL of LB-medium (Luria/Miller) (Carl Roth) or YEB medium, supplemented with appropriate antibiotics (i.e. 100 g/L spectinomycin). After incubation, the bacterial culture was transferred to 15 ml Falcon tubes and centrifuged (10 min, 5,000 rpm). The pellets were washed with 5 mL of the infiltration buffer (10 mM MgCl_2_, 10 mM MES pH 5.7) and the final pellet resuspended in the infiltration buffer supplemented with 100-150 μM acetosyringone. The bacterial suspension was diluted with supplemented infiltration buffer to adjust the inoculum concentration to a OD600 value of 0.025-1.0. The inoculum was incubated for 2-3 h at room temperature before injecting and delivered to tobacco by gentle pressure infiltration of the lower epidermis leaves (fourth and older true leaves were used and about 4/5-1/1 of their full size) with a 1-mL hypodermic syringe without needle (Moschou et al., 2016).

Arabidopsis cell suspension: The PSB-D Arabidopsis thaliana cell suspension cultures were transformed with the POI: Pro35S∷GFP/TPLATE/TML-BioID/BioID2, Pro35S∷TPLATE/TML-(GGGGS)13 BioID/BioID2 and Pro35S∷GFP/TPLATE-(GGGGS)13 TurboID and selected without callus screening, grown and subcultured as described by (Van Leene et al., 2007).

Arabidopsis plants to express TOL6-Venus: Flowering *tol2-1/tol2-1 tol5-1/tol5-1 tol6-1/tol6-1 tol9-1/tol9-1* plants were transformed with *Agrobacterium tumefaciens* using the floral dip method (Clough and Bent, 1998). Resulting T2 lines were confirmed for single-transgene insertion sites and propagated for further analysis. At least three independent transformants were characterized for each line. Homozygous plants were confirmed by PCR genotyping for the mutant alleles (Korbei et al., 2013).

### Biotin treatments

#### Hairy roots

For assessing self-biotinylation, 2 weeks old 25 ml liquid cultures were added 5 ml fresh MS medium with or w/o supplemented biotin (i.e. 50 μM f.c.; stock solution dissolved in water) for 2h or 24h and samples collected. Two independent root cultures were analyzed per combination and the experiment repeated twice with similar results.

#### *N. benthamiana* leaves

Plants were kept under normal growing conditions 22°C, re-infiltrated with infiltration buffer (no biotin) or alternatively, infiltration buffer supplemented with biotin (stock solution dissolved in DMSO or water) and samples collected at the indicated times points. Two infiltrated tobacco leaf segments/leaves were analyzed per combination.

#### *Arabidopsis* cell cultures

were grown under normal conditions, at 25°C at 130 rpm in the dark. 48 h after subculturing, the required amount of biotin was added and the cell culture was transferred to the desired temperature for the required time at 130 rpm shaking in the dark in an INCLU-line IL56 (VWR) incubator. After the needed time, cell cultures were harvested and flash-frozen in liquid nitrogen and stored at −70° till used.

### Protein extractions

#### Hairy roots

The tissue samples were flash-frozen and crushed using a liquid cooled mortar and pestle and the crushed material was transferred to a 1.5 ml Eppendorf in homogenization buffer (25 mM Tris-HCl pH 7.6, 15 mM MgCl_2_, 5 mM EGTA, 150 mM NaCl, 15 mM pNO_2_PhenylPO_4_, 15 mM β-glycerolphosphate, 1 mM DTT, 0.1% NP-40, 0.1 mM Na_3_VO_4_, 1 mM NaF, 1 mM PMSF, 10 μg/ml leupeptin, 10 μg/ml aprotinin, 10 μg/ml SBTI, 0.1 mM benzamidine, 5 μg/ml antipain, 5 μg/ml pepstatin, 5 μg/ml chymostatin, 1μM E64, 5% ethylene glycol) was added with volumes according to the dry weight of the recovered material (1/1 w/v) and protein material extracted by three repetitive freeze-thaw cycles in liquid nitrogen and the lysate transferred to a 1.5 ml Eppendorf. The lysates were cleared by centrifugation for 15 min at 16,100 x g (4 °C) and the supernatant transferred to a new 1.5 ml Eppendorf. This step was repeated two times and the protein concentration was determined by the DC Protein Assay Kit (Bio-Rad, Munich, Germany) according to the manufacturer’s instructions.

#### *N. benthamiana* leaves

The tissue samples were crushed using a liquid cooled mortar and pestle and the crushed material transferred to a 1.5 ml Eppendorf in homogenization buffer. Leaves were harvested and directly frozen in liquid nitrogen. Proteins were extracted with buffer containing 50 mM Tris-HCl (pH 7.5), 150 mM NaCl, 10 % glycerol, 2 mM EDTA, 5 mM DTT, 1 mM PMSF, Protease inhibitor Cocktail (Roche) and 1 % (v/v) IGEPAL CA-630 (Sigma-Aldrich). Extraction buffer was added at 2 ml/g tissue. Extracts were incubated at 4 °C for 1 h and then centrifuged at 4 °C, 13000 rpm for 30min. Supernatants were used directly or filtered through PD-10 columns (GE Healthcare) and incubated with streptavidin (Roche) or GFP (Chromotek) beads for 1 h. For ammonium acetate protein precipitation, supernatants were precipitated using 5x v/v pre-cold 0.1 M ammonium acetate in methanol at −20 °C for 2h and then centrifuged at 4 °C, 13,000 rpm for 15min. The pellet was washed with pre-cold 0.1 M ammonium acetate and dissolved in the same extraction buffer plus 1% SDS. Magnetic separation was done using Dynabeads™ M-280 Streptavidin (Thermo Fisher Scientific) followed by 5 times washing in buffer containing 50 mM Tris-HCl (pH 7.5), 150 mM NaCl, 10 % glycerol, 2 mM EDTA, Protease inhibitor Cocktail (Roche) and 0.5 % (v/v) IGEPAL CA-630 (Sigma-Aldrich) and one time in buffer containing 50 mM Tris-HCl (pH 7.5), 1M NaCl, 10 % glycerol, 2 mM EDTA, Protease inhibitor Cocktail (Roche) and 0.5 % (v/v) IGEPAL CA-630 (Sigma-Aldrich) at 4°C. To release the proteins, 100 μl 2x NuPAGE LDS sample buffer (Invitrogen) was added and samples were heated for 5 min at 95 °C

#### Arabidopsis cell cultures

Total protein extracts were obtained from biotin treated, harvested and liquid nitrogen retched (20 Hz, 1 min), *Arabidopsis* cell suspension cultures using double the volume (w/2v) of extraction buffer containing 150 mM Tris-HCl pH 7.5; 150 mM NaCl; 10 % glycerol; 10 mM EDTA; 1mM sodium molybdate; 1 mM NaF and freshly added 10 mM DTT; 1 % (v/v) protease inhibitor cocktail (P9599, Sigma (1 tablet/10ml elution buffer) and 1 % (v/v) NP-40. Cell debris was removed by two rounds of centrifugation at 14,000 rpm for 20 min at 4°C and the supernatant was collected.

### SDS-PAGE and Western blots

#### Hairy roots

Sample loading buffer was added and equivalent amounts of protein (~ 30 μg) separated by SDS-PAGE (1.0 mm thick 4 to 12% polyacrylamide Criterion Bis-Tris XT-gels, Bio-Rad or equivalent) in MOPS buffer (Bio-Rad) at 150 V. Subsequently, proteins were transferred onto PVDF membranes with 0.2 um porous size. Membranes were blocked for 30 min in a 1:1 Tris-buffered saline (TBS)/Odyssey Blocking solution (cat n° 927-40003, LI-COR, Lincoln, NE, USA) and probed by Western blotting. Following overnight incubation of primary antibody in TBS-T/Odyssey blocking buffer and three 10 min washes in TBS-T (0.1% Tween-20), membranes were incubated with secondary antibody for 30 min in TBS-T/Odyssey blocking buffer followed by 3 washes in TBS-T or TBS (last wash step). The following antibodies were used: streptavidin-S680 (Invitrogen, S32358, 1/10000), mouse anti-Flag (Sigma, F3165; 1/5000), mouse anti-actin (plant) (Sigma, A0480, 1/2000), rabbit anti-GFP (Invitrogen, A11122, 1/1000), anti-mouse (IRDye 800 CW goat anti-mouse antibody IgG, LI-COR, cat n° 926-32210, 1/10000) and anti-rabbit (IRDye 800 CW goat anti-rabbit IgG, LI-COR, cat n° 926-3221, 1/10000). The bands were visualized using an Odyssey infrared imaging system (LI-COR) and the intensity of bands assessed using the LICOR Odyssey software for Western Blot image processing.

#### N. benthamiana

Extracted proteins were loaded to 12% SDS-PAGE gels and separated for 2 h at 90-110V. SDS-PAGE gels were blotted via wet transfer on PVDF membranes (Carl Roth) overnight at 30V. Membrane blocking was performed with 3%BSA in PBS-t buffer for 1 h at room temperature followed by incubation with Mouse-anti-GFP (TaKaRa) (1/5,000) for 2 h followed by Anti-Mouse-HRP (Sigma-Aldrich) (1/5,000) for 2 h or directly Strep-Tactin-HRP (iba-Life Sciences) (1/5,000) for 2 h. Chemiluminescence was detected with Clarity Western ECL (Bio-rad).

#### N. benthamiana

Input and eluted proteins were loaded to 12% SDS-PAGE gels and separated for 1-2 h at 120 V. SDS-PAGE gels were blotted via wet transfer on PVDF membranes (Bio-rad) 3h at 300 mA in a cool room. The membrane was blocked with 3% BSA in PBS-T buffer for 1 h at room temperature followed by incubation with Streptavidin-HRP (Sigma-Aldrich) (1/25,000) for 2 h. Chemiluminescence was detected with ECL Prime Western Blotting Detection Reagent (GE healthcare).

#### Arabidopsis cell cultures

The total protein extracts were heated in sample buffer for 10 min at 70°C and loaded in equal amounts (20 μg, protein concentration was measured using a qubit system, ThermoFischer) on a 4-20% SDS-PAGE gel. SDS-PAGE separated proteins were blotted on PVDF membrane (Thermo Fisher). Membranes were blocked overnight at RT in 5% (v/v) BSA dissolved in 25 mM Tris-HCl, pH 8, 150 mM NaCl and 0.1% Tween20. The blots were then incubated at room temperature with the Pierce High Sensitivity Streptavidin-HRP Thermo Fisher scientific (1/2,000) or Abcam Anti-HA-HRP tag antibody (ab1190) (1/5,000) in 1% BSA made as mentioned above for 2 h. Antigen-antibody complexes were detected using chemiluminescence (Perkin-Elmer).

### Imaging analysis

A *tplate* mutant complemented line expressing proLAT52∷TPLATE-TagRFP (Wang et al., BioRxiv 948109) was crossed with a quadruple tol (*tol2/tol2 tol5/tol5 tol6/tol6 tol9/tol9*) mutant line expressing proTOL6∷TOL6-Venus. F1 seedlings were imaged using spinning dics microscopy. Etiolated hypocotyl cells of 4-day old seedlings expressing TPLATE-TagRFP and TOL6-Venus were imaged with a Nikon Ti microscope equipped with an Ultraview spinning-disk system (PerkinElmer) and a 512 × 512 Hamamatsu ImagEM C9100-13 EMccd camera. Images of hypocotyl epidermal cells were acquired with a 100x oil immersion objective (Plan Apo, NA = 1.45). TOl6-Venus was imaged with 514 nm excitation light and an emission window between 525 nm and 575 nm. TPLATE-TagRFP was imaged with 561 nm excitation light and an emission window between 570nm and 625nm. Dual-color images were acquired sequentially with an exposure time of 500 ms/frame.

Objects based co-localization was performed using the plugin Distance Analysis (DiAna) of ImageJ (Gilles et al., 2017). Prior to analysis with the DiAna plugin, images were processed with ImageJ. Each channel is processed using a Walking Average of 4 and then merged (also rotated if required). Regions of interest within each image were selected based on that they excluded the border of the cells and still contained a good number of objects. Z-projection images were generated using five frames with average intensity. Then, each channel of Z-projected images was processed with Morphological filters from the MorphoLibJ plugin (Legland et al., 2016), using the parameters white top-hat, disk element and a 2 pixel radius. Objects for each channel were segmented by selecting the 3D Spot segmentation tool. We adapted the calibration by changing the pixel size to 1.00001 for all dimensions. Both the noise and seed threshold value were obtained by averaging the maximum intensity of three regions covering only background signal. The spot was defined using a minimum value of 4 and maximum value of 36 pixels. The option to exclude objects on XY edges was activated. Default values were used for the other parameters. Results for number of total objects (Tot) or touching objects (Tou) in image A/B obtained from Diana were recorded. The colocalization ratio of objects was calculated as follows:

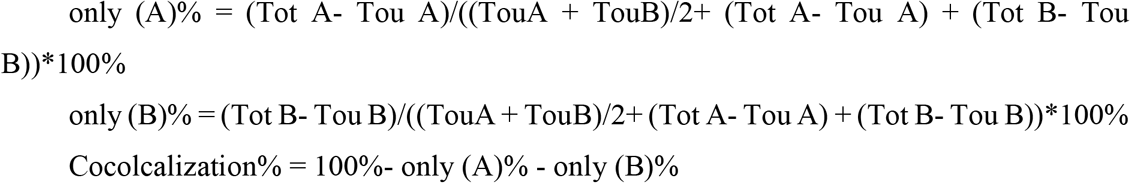

As a control, one of the channels was horizontally flipped, merged with the other channel and analyzed. 8 cells originating from 3 seedlings were analyzed.

### Bimolecular Fluorescence Complementation

Ratiometric BiFC images were obtained using an Olympus FV1000 inverted confocal microscope equipped with a UPLSAPO 60x water immersion objective (NA 1.2). Images were acquired in line sequential mode, using 515 nm excitation and an emission window between 530 nm to 548 nm for the YFP detection and using 559 nm excitation and an emission window between 580 nm to 615 nm for RFP detection. All images were taken using the exact same settings. The experiment was independently repeated twice with similar outcome.

For the quantification of the YFP/RFP ratio, only images with less than 1% saturation in the RFP or YFP channel were analysed. For each confocal image, parts of the cortical cytoplasm in the RFP channel were traced in ImageJ using the selection brush tool with a width of 15 pixels. Histogram analysis was performed to confirm that less than 1% saturated pixels were present in the ROI. The average intensity from the obtained ROI was calculated and divided by the average intensity of the same region in the YFP channel. Ratios were quantified for 15 to 19 individual cells.

Outliers were removed by iterative outlier removal (Leys et al., 2013). Data were analyzed using Rstudio (RStudio Team (2015). RStudio: Integrated Development for R. RStudio, Inc., Boston, MA URL http://www.rstudio.com/) with Welch-corrected ANOVA to account for heteroscedasticity. Post hoc pairwise comparison was performed with the package MULTCOMP utilizing the Tukey contrasts (Herberich et al., 2010).

### Protein Extraction and Pull down for Mass Spectrometry analysis

#### For Figure 3

*Arabidopsis* cell cultures expressing different POI were ground in 0.67 volume of extraction buffer containing 150 mM Tris-HCl pH 7.5; 150 mM NaCl; 10 % glycerol; 10 mM EDTA; 1mM sodium molybdate; 1 mM NaF and freshly added 10 mM DTT; 1 % (v/v) protease inhibitor cocktail (P9599, sigma (1 tablet/10ml elution buffer), 1 % (v/v) digitonin and benzonase 0.1% (w/v). The extract was mixed using ultra-Turrax for 3×30” at 16000 rpm and sonicated for 15”x 3 with 30” interval. The extract was incubated on a rotating wheel for 1 hour at 4°C. Cell debris was removed by two rounds of centrifugation at 20,000 rpm for 20 mins at 4°C and the supernatant was buffer exchanged using pre-equilibrated PD-10 columns and eluted in binding buffer (extraction buffer without digitonin and benzonase) at 4° C. Pull-downs were performed in triplicate. For each pull-down, 1/3 of the soluble protein extract was incubated with 200 μl slurry of streptavidin sepharose high-performance beads (Amersham) (pre-equilibrated with binding buffer) overnight on a rotating wheel at 4°C. The unbound fraction or supernatant, was removed after centrifugation at 1,500 rpm for 1 min. Beads were transferred to a mobicol column and washed with 2.5 ml binding buffer followed by wash with 2.5 ml of wash buffer-1 containing 25mM Tris-HCl (pH7.5); 150mM NaCl; digitonin 0.1% (w/v). The beads were washed once with wash buffer-2 containing 25mM Tris-HCl pH7.5; 150 mM NaCl and finally washed once with 50 mM ammonium bicarbonate pH 8.0. Proteins were digested on beads with Trypsin/LysC (Promega) overnight followed by zip-tip cleanup using C-18 Omix tips (Agilent). Digests containing the unbiotinylated peptides were dried in a speedvac and stored at −20 °C until LC-MS/MS analyses.

#### For Figure 4 and Figure 5

*Arabidopsis* cell cultures expressing different POI were ground in 0.67 volume of extraction buffer containing 100 mM Tris (pH 7.5), 2% SDS and 8M Urea. The extract was mechanically disrupted using three repetitive freeze-thaw cycles followed by 2 cycles of sonication at output level 4 with a 40% duty cycle for 50” with 35” interval. The extract was incubated at rotating wheel for 1 hour at RT. Cell debris was removed by two rounds of centrifugation at 20,000 rpm for 20 mins at RT and the supernatant was buffer exchanged using pre-equilibrated PD-10 columns and eluted in binding buffer containing 100 mM Tris (pH 7.5), 2% SDS and 7.5M Urea. Pull-downs were performed in triplicate. For each pull-down, 1/3 of the soluble protein extract was incubated with 200 μl slurry of streptavidin sepharose high-performance beads (Amersham) (pre-equilibrated with binding buffer) overnight on a rotating wheel at RT. The unbound fraction or supernatant, was removed after centrifugation at 1,500 rpm for 1 min. Beads were transferred to a mobicol column and washed with 4 ml binding buffer for 5 mins without agitation, followed by a wash with high salt buffer containing 1M NaCl, 100 mM Tris-HCl pH 7.5 and incubated for 30 mins. The beads were washed once with ultrapure water, incubated for 5 mins and finally washed with 3.2ml of 50 mM ammonium bicarbonate pH 8.0 incubating 5 mins. Proteins were digested on beads with Trypsin/LysC (Promega) overnight followed by zip-tip cleanup using C-18 Omix tips (Agilent). Digests containing the unbiotinylated peptides were dried in a Speedvac as elution-1 (E1) and stored at −20 °C until LC-MS/MS analyses.

After E1, for all linkerTurboID samples, biotinylated peptides were eluted from the beads, by adding 300μl of the elution buffer containing 0.2% TFA, 0.1% formic acid and 80% acetonitrile in water. The eluted peptides were collected by centrifugation at 1500 rpm for 1 min followed by an addition to the beads of 300μl of the elution buffer, after which the sample was heated at 95°C for 5 min to allow a maximal release of peptides. A short spin at 1,500 rpm for 1 min was done to collect the eluted peptides. The two elutes were pooled and dried in a speedvac. The dried peptides were dissolved in 1% TFA solution to perform zip-tip cleanup using C-18 Omix tips (Agilent). Digests were dried in a speedvac as elution-2 (E2) and stored at −20 °C until LC-MS/MS analysis.

TPLATE-CGSrhino pull-downs with home-made IgG beads were performed as described in (Van Leene et al., 2019).

### Mass Spectrometry and Data Analysis

Triplicate pull-down experiments were analyzed by LC-MSMS on Q Exactive (ThermoFisher Scientific) as previously reported (Nelissen et al., 2015).

For comparison of TPLATE-BioID at 25, 28, 30 and 35 degrees, raw data of GFP-BioID and TPLATE-BioID triplicates at the different incubation temperatures were searched together with MaxQuant (Tyanova et al., 2016a) using standard parameters (Supplemental Dataset 1). LFQ intensities were used in Perseus software (Tyanova et al., 2016b) to determine the significantly enriched proteins with TPLATE for each sample set, TPLATE versus GFP at respectively 25, 28, 30 and 35 degrees. Thereto the MaxQuant proteingroups file, with reverse, contaminant and only identified by site identifications already removed, was loaded in Perseus. Samples were grouped by the respective triplicates and filtered for minimal 2 valid values per triplicate. Missing LFQ values were imputated from normal distribution using standard settings in Perseus, width of 0.3 and down shift of 1.8. Next, ttests were performed and visualized in volcano plots, using permutation-based FDR to determine the significantly different proteins between TPLATE-BioID and GFP-BioID at the different incubation temperatures. As cut-off, FDR=0.05, S0=0.5 was applied. Protein lists significantly enriched with TPLATE can be found in Supplemental Table 4. For all TPC subunits, the values for Difference and −log(p-value) from the Perseus t-test were presented in Figure 3, in order to compare the different TPLATE-BioID samples and determine the optimal temperature for BioID.

For comparison of different TPLATE PBLs at 28 degrees, triplicate TPLATE-BioID, TPLATE-BioID2, TPLATE-linkerBioID, TPLATE-linkerBioID2, TPLATE-linkerTurboID and respective controls GFP-BioID, GFP-BioID2 and GFP-linkerTurboID raw data was searched together in MaxQuant with standard parameters (Supplemental Table 5). Datasets were further processed in the same way as described for the comparison of the different incubation temperatures. GFP-BioID served as control for TPLATE-BioID and TPLATE-linkerBioID, and GFP-BioID2 served as control for TPLATE-BioID2 and TPLATE-linkerBioID2. For linkerBioID, next to 50μM biotin incubation, also 2mM biotin incubation was tested. Pairwise comparisons were made between the different TPLATE PBLs and their respective controls, and a cut-off of FDR=0.05, S0=0.5 was applied. Protein lists significantly enriched with TPLATE can be found in Supplemental Table 5. For all TPC subunits, the values for Difference and −log(p-value) from the Perseus ttests were presented in Figure 4, in order to compare the different TPLATE PBLs.TPLATE-CGSrhino pull-downs were analyzed as described in Van Leene et al., 2019. Briefly, pull down triplicates were analyzed by LC-MSMS on Q Exactive (ThermoFisher Scientific), raw data were searched with the Mascot search engine (Matrix Science) and average Normalized Spectral Abundance Factors (NASF) for the identified proteins were compared in a t-test versus a large dataset of similar experiments consisting of non-related baits. Proteins not present in the background list or highly enriched versus the large dataset were kept as significant set. Thresholds used are NSAF ratio bait/large dataset >=10, and −log(p-value) >=10. In this case, 1 peptide identifications were retained, otherwise the small TPC subunit LOLITA would have fallen out of the data. The significant set can be found in Supplemental Table 6.

For comparison of the different PBLs versus GSrhino pull-down samples, a MaxQuant search was performed on all relevant TPLATE raw data together. Since LC-MSMS analysis is done the same for the GSrhino as for the Streptavidin pull downs, it’s also possible to include matching between runs. Next, resulting iBAQ values were used for comparison of the abundance of the identified proteins amongst the different TPLATE samples. For completeness, one peptide identifications were allowed, in order to also obtain iBAQ values for the one peptide identifications. The complete set of significant proteins as determined by previous analysis, for the PBLs and for CGSrhino, as described, with their iBAQ values can be found in Supplemental Table 6. In Figure 5, a subset of relevant endocytosis related proteins is presented.

In order to compare TPLATE-linkerTurboID with different incubation times and at different temperatures, triplicate Streptavidin pull-downs were performed with TPLATE-linkerTurboID and GFP-linkerTurboID at 25 degrees with different incubation times with 50μM biotin. Analysis to determine the significant identifications in each TPLATE-linkerTurboID set versus the respective GFP-linkerTurboID control was done as described before. Significant lists can be found in Supplemental Table 7. For a direct comparison of the different incubation times and temperatures with TPLATE-linkerTurboID, a MaxQuant search was performed on all relevant TPLATE raw data together, with matching between runs. Next, resulting iBAQ values were used for comparison of the abundance of the identified proteins amongst the different sample sets. Again, for completeness, one peptide identifications were allowed, in order to obtain iBAQ values also for the one peptide identifications. For this comparison between linkerTurboID samples only, both elutions were taken together for all sample sets, for the determination of the significant sets as well as for the comparison of the iBAQ values. The complete set of significant proteins with their iBAQ values can be found in Supplemental Table 6. In figure 6, a subset of proteins is presented.

For each of the TPC subunits, the identified peptides in the TPLATE-linkerTurboID replicates were mapped to the protein sequence, see Figure 7. Non-biotinylated and biotinylated peptides of TPC subunits were mapped to the protein sequence by using the Draw Map tool in the MSTools package (http://peterslab.org/MSTools/ (Kavan and Man, 2011) and put together using Inkscape v 0.92.4 (www.inkscape.org). Domain annotation of TPC subunits was retrieved using InterPro protein sequence analysis (https://www.ebi.ac.uk/interpro/) (Mitchell et al., 2018).

## SUPPLEMENTAL MATERIALS

Supplemental Figure 1. Overview of available constructs for proximity biotinylation in plants.

Supplemental Figure 2. GFP expression in tomato hairy root cultures produced with rhizogenic *Agrobacterium*.

Supplemental Figure 3. Characterization of PBL-catalysed proximity labelling in *N. benthamiana*.

Supplemental Figure 4. Biotinylation of BioID increases at elevated growth temperature and biotin concentration in Nicotiana benthamiana.

Supplemental Figure 5. *Trans*-biotinylation within membrane-resident receptor complexes.

Supplemental Figure 6. Different PBL cause different *cis*- and *trans*-biotinylation.

Supplemental Figure 7. *Cis*-biotinylation of TPLATE-linkerBioID increases at higher concentration of exogenous biotin.

Supplemental Figure 8. Exogenous application of biotin can exceed the binding capacity of streptavidin beads.

Supplemental Figure 9. The biotin-streptavidin interaction is retained under harsh conditions.

Supplemental Table 1. List of expression vectors used in this study.

Supplemental Table 2. list of primers used.

Supplemental Table 3. Cell cultures expressing different TPLATE-PBLs identifies TPC subunits with different amount of non-biotinylated peptides.

Supplemental Table 4. Full list of significantly enriched identifications with TPLATE-BioID versus GFP-BioID at different incubation temperatures for 24 hours with 50 μM biotin.

Supplemental Table 5. Full list of significantly enriched identifications with TPLATE as bait using BioID, BioID2, linkerBioID, linkerBioID2, linkerTurboID E1, linkerTurboID E2 and linkerTurboID E1+E2, versus the respective GFP PBLs.

Supplemental Table 6. Full list of significantly enriched hits with either TPLATE PBLs or GSrhino PD, including average iBAQ values, normalized versus TPLATE.

Supplemental Table 7. Full list of significantly enriched hits with TPLATE-linkerTurboID, including average iBAQ values, normalized versus TPLATE, at different incubation times at 25 degrees and 24-hour incubation time at 28 degrees, all with 50 μM biotin.

Supplemental Table 8. Supplemental sequences. List of all used PBL sequences.

## ACKNOWLEDGEMENTS

Part of this work was funded by the European Research Council T-Rex project number 682436 to D.V.D.) and by the National Science Foundation Flanders (FWO; G009415N to D.V.D. and G.D.J.). T.O. and N.B.A. were funded by the Deutsche Forschungsgemeinschaft (DFG, German Research Foundation) in frame of the Collaborative Research Center 924 (Sonderforschungsbereich 924, INST 95/1126-2, Project B4). P.N.M. funding in support of this work was from VR (no. 21679000) and FORMAS Research Councils (no. 22924-000), the Carl Tryggers Foundation (grant nos 15:336 and 17:317), and start-up grants from IMBB-FORTH.

## AUTHOR CONTRIBUTIONS

D.A., N.B.A., C.L., P.V.D., K.Y. J.W., B.K. and A.T. designed and/or performed research. L.D.V., B.K, F.I., D.E. contributed new analytic tools and analyzed data. P.V.D., A.G., G.D.J., T.O., P.M. and D.V.D. designed research, analyzed data and wrote the paper. All authors contributed to finalizing the manuscript text.

