## supplemental material for "Establishment of Proximity-dependent Biotinylation Approaches in Different Plant Model Systems"

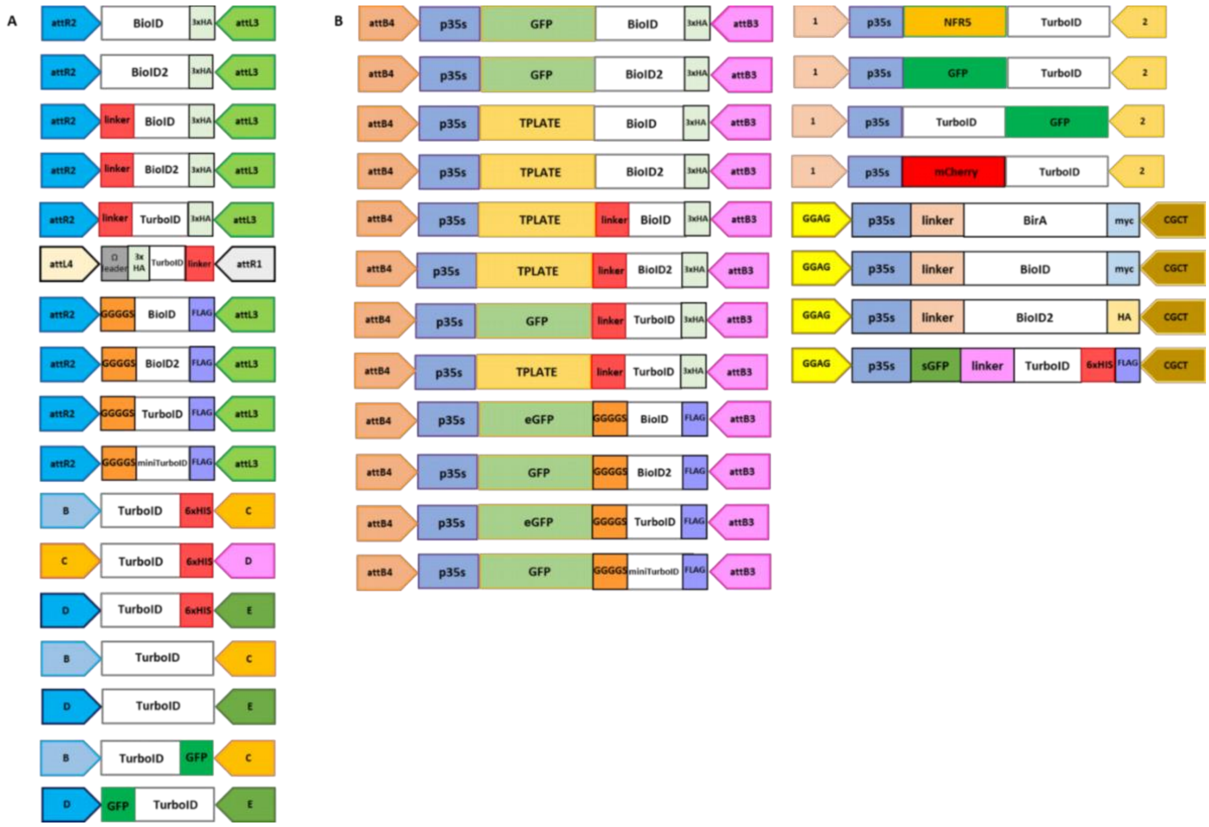


**Supplemental Figure 1. Overview of available constructs for proximity biotinylation in plants.** Diagrammatic representation of plasmids used in the study and/or available for the community. (A) various entry clones or building blocks used to make the destination vectors in panel B. (B) Destination vectors used in this study and/or available to the community. Gateway att sites are according to Karimi et al., 2007. Borders of golden gateway B-E and 1,2 are according to GoldenGate from Binder et al 2014, whereas the 4 nucleotide borders of the golden gateway system are according to GoldenGate from Patron et al., 2015. The linker (in red) is a 65 aa long $(GGGGS)_{13}$ repeat. The linker (in light orange) is a 213bp GS linker while the one in grey is 30 bp GS linker. BioID, BioID2, TurboID and miniTurbo all represent promiscuous versions of the respective biotin ligases. BirA is the non-promiscuous negative control version of the biotin ligase. Myc, HA, 6xHis and 1x or 3xFLAG are added tags for Western Blot detection. All corresponding sequences can be found in supplemental sequences.

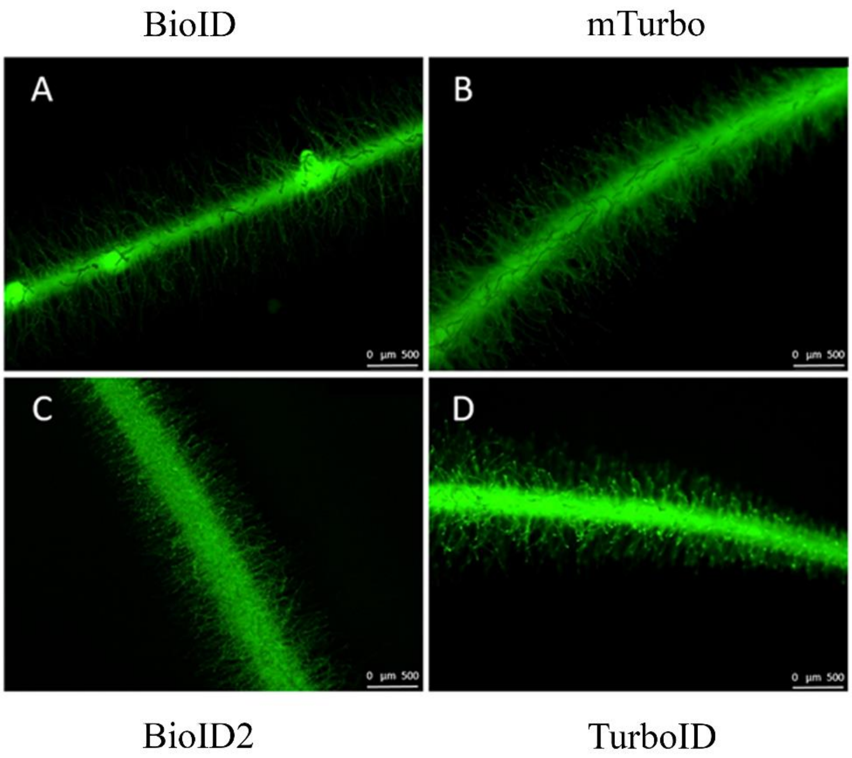


**Supplemental Figure 2. GFP expression in tomato hairy root cultures transformed with rhizogenic *Agrobacterium*.** Fluorescence micrograph images of eGFP expression were obtained from primary hairy roots (~14 days) transformed by rhizogenic *Agrobacterium* with the following expression constructs; Pro35S::eGFP-BioID, Pro35S::eGFP-BioID2, Pro35S::eGFP-TurboID, and Pro35S::eGFP-mTurbo. The scale bars are 500 μm. Images are representative for the four to ten independent roots selected for sub cultivation and showing expression of the marker per construct. This figure supports Figure 1.

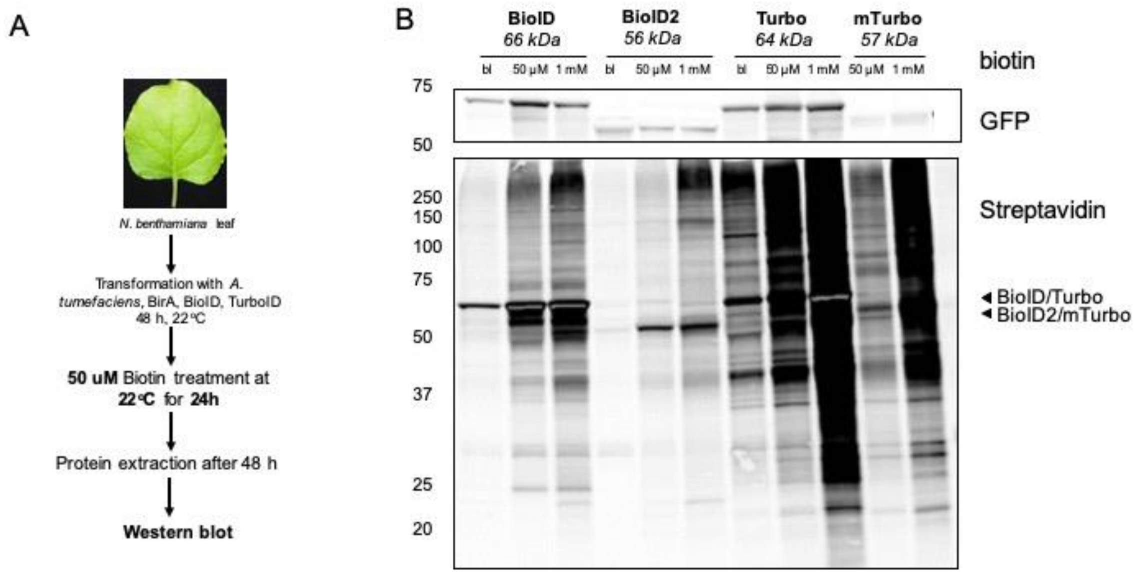


**Supplemental Figure 3. Characterization of PBL-catalysed proximity labelling in *N. benthamiana*.** (A) Experimental setup. (B) Comparison of biotinylation activity in *N. benthamiana* expressing eGFP-BioID (~66 kDa), eGPF-BioID2 (~56 kDa), eGFP-Turbo (~64 kDa) and eGPF-mTurbo (~57 kDa). Overlapping signal as indicated with a black arrow denote enzyme-catalysed *cis*-biotinylation. Gray bands in intense black areas represent saturation of the streptavidin-s680 signal and is most prominent in case of auto-biotinylation activity. Blank lanes (bl) depict endogenously biotinylated proteins. Two infiltrated tobacco leaf segments/leaves were analyzed per setup and the experiment was repeated twice with similar results. This figure supports Figure 1.

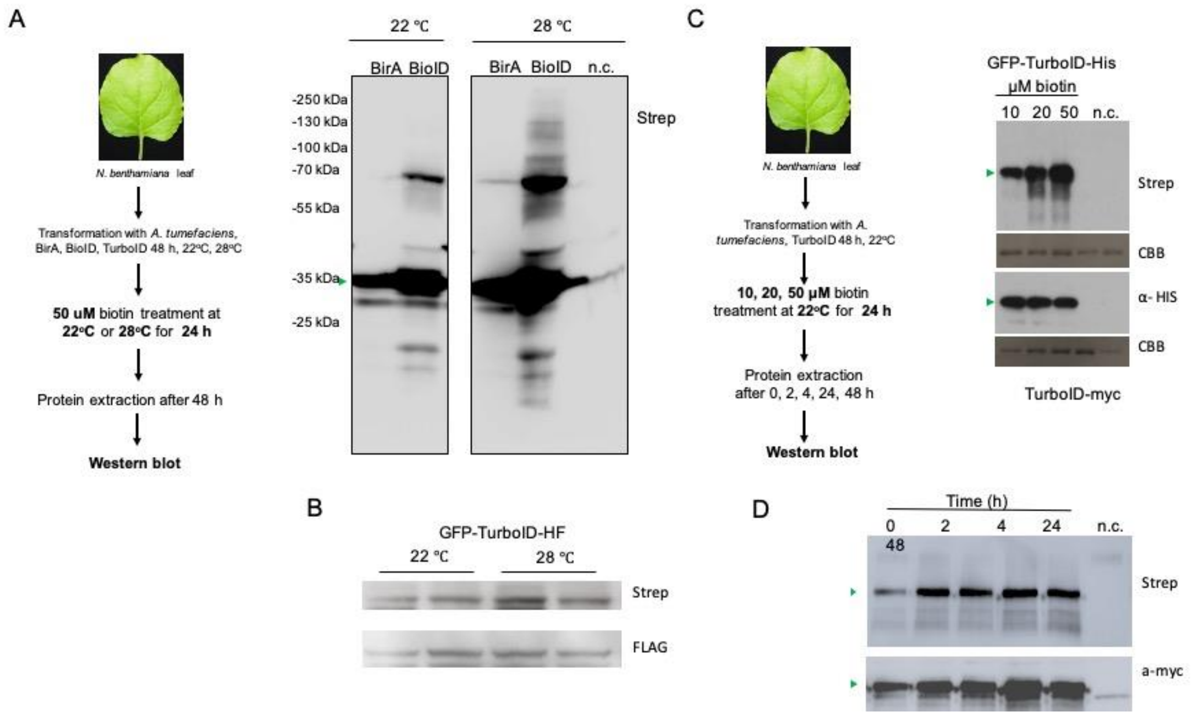


**Supplemental Figure 4. Biotinylation of BioID increases at elevated growth temperature and biotin concentration in *Nicotiana benthamiana*.** (A) The left panel shows the experimental setup. Western blot: The wild-type version of BioID (BirA) shows residual cis-biotinylation activity at both 22°C and 28°C. (B) BioID shows promiscuous biotinylation which increases with temperature. The samples were run on the same blot along with other ones. The blot was cropped to allow a direct comparison between the samples shown. GFP-TurboID-HF activity was ~2 fold increased at higher temperature (i.e. when going from 22°C to 28°C). Two biological replicates are depicted. (C) The left panel shows the experimental setup. Increasing biotin levels elevated biotinylation efficiency. (D) Efficient cis-biotinylation was observed already 2 h after biotin administration. For (C) and (D), green arrowheads show the corresponding BL. N.c. negative control (empty vector). This figure supports Figure 2.

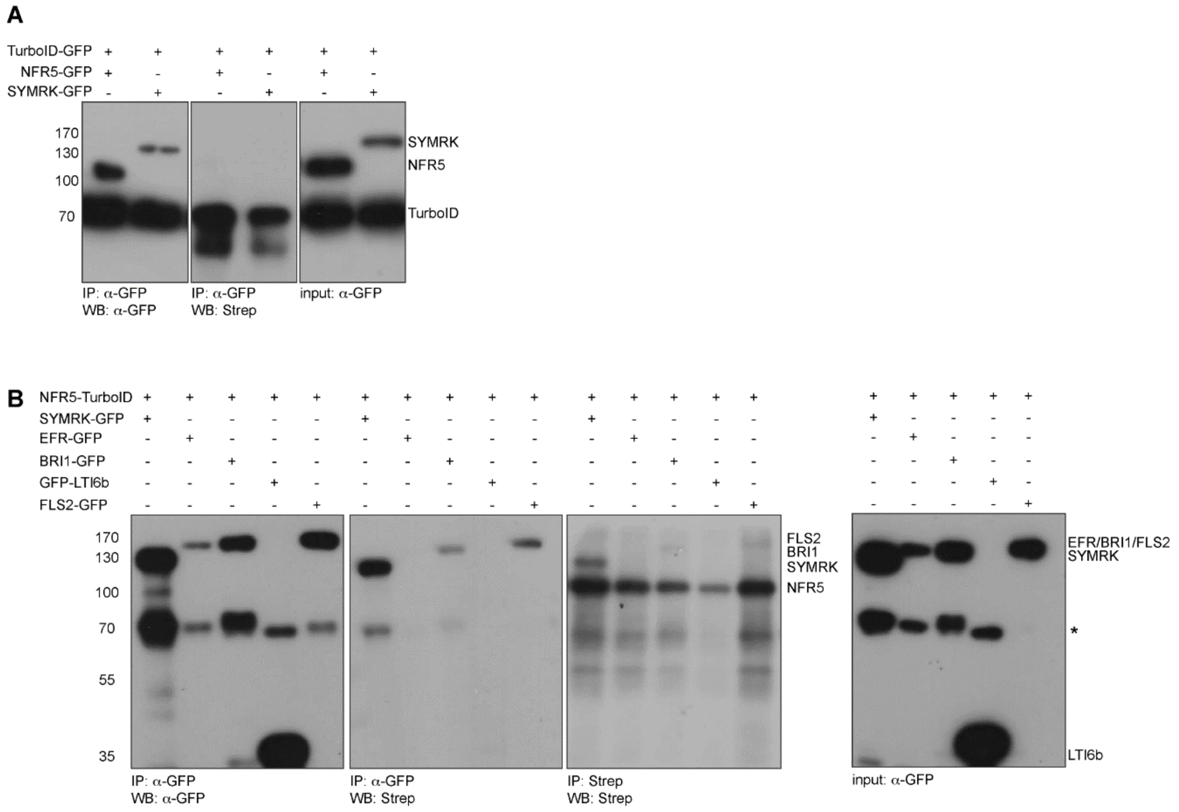


**Supplemental Figure 5. *Trans*-biotinylation within membrane-resident receptor complexes.** (A) TurboID-GFP was co-expressed with symbiotic RLKs to test for unspecific *trans*-biotinylation. While all proteins were detected before (right panel) and after immunoprecipitation (left panel), no *trans*-biotinylation of the receptors was observed under these conditions (middle panel). The activity of TurboID is indicated by *cis*-biotinylation of TurboID-GFP (70 kDa). (B) Fusing TurboID to NFR5 (120 kDa) resulted in strong *trans*-biotinylation of the known interaction partner SYMRK (150 kDa), weak signals were detected in case of the PM-resident receptors BRI1 and FLS2, while no *trans*-biotinylation of EFR and the PM-marker LTI6b were detected. Temporally limiting the reaction results in weak but specifically detectable bands in case of NFR5-TurboID and SYMRK-GFP. Biotin was applied for 2 hours. IP= immunoprecipitation; WB= Western Blot. *= unclassified band. Strep= Streptavidin. This figure supports Figure 2.

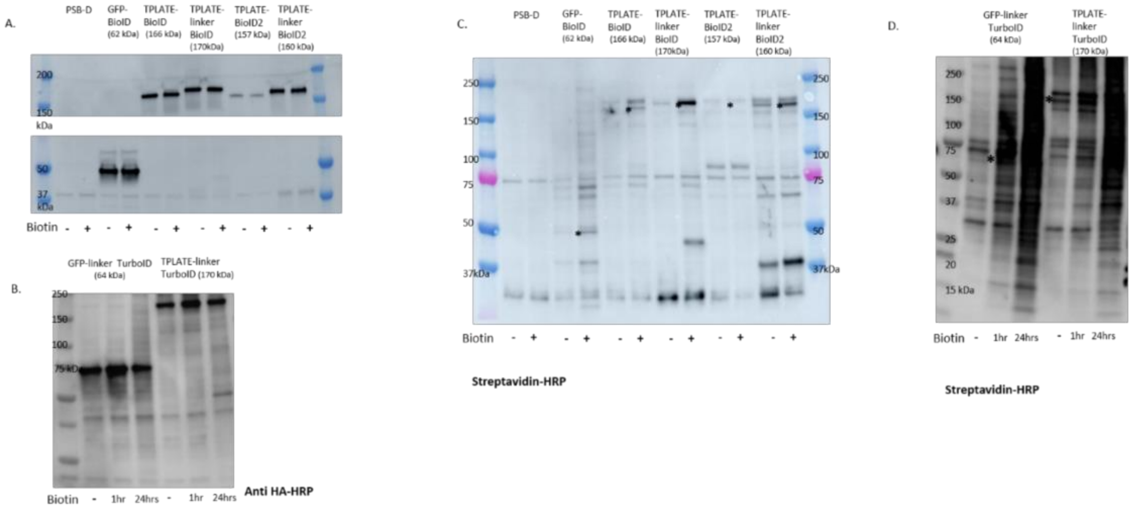


**Supplemental Figure 6. Different PBL cause different *cis*- and *trans*-biotinylation**. *Arabidopsis* cell cultures expressing different TPLATE-PBLs were incubated with 50µM biotin at 28°C for 24 h or for both 24 h and 1 h (linkerTurboID cultures) before harvesting. (A and B) Anti-HA HRP Western blotting was performed to visualize expression levels of the different cultures. (C and D) Streptavidin-HRP Western blotting of different TPLATE-PBLs, GFP-BioID and control cell cultures (PSB-D). *Cis*-biotinylation of the bait and *trans*-biotinylation can clearly be observed. * indicates *cis*-biotinylation of the bait. This figure supports Figure 4.

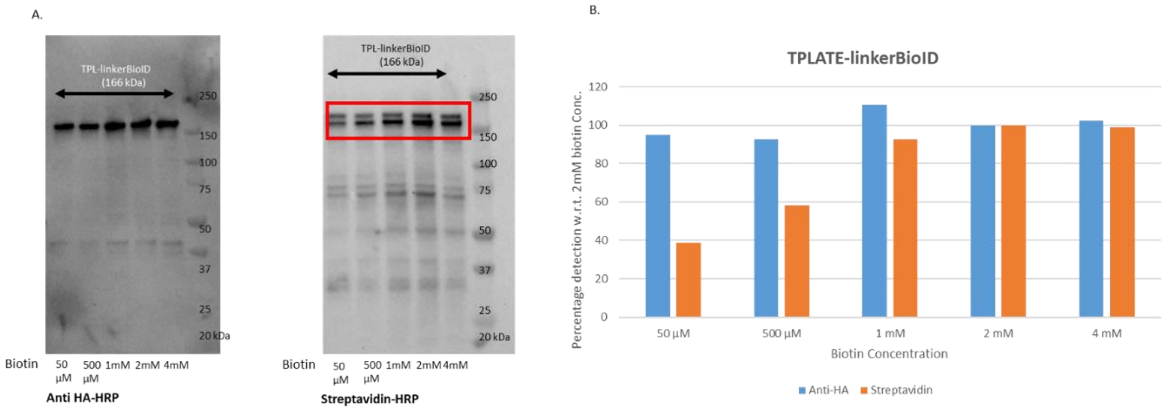


**Supplemental Figure 7. *Cis*-biotinylation of TPLATE-linkerBioID increases at higher concentration of exogenous biotin.** Cell cultures expressing TPLATE-linkerBioID were incubated with biotin (50µM to 4mM) at 28°C for 24 h. (A) Anti-HA HRP Western immunostaining was performed to check protein expression while streptavidin-HRP Western immunostaining was used to assess the biotinylation levels of TPLATE-BioID. (B) Quantification of the percentage of biotinylation (orange) as well as the expression (blue) for each biotin concentration normalized to the maximum biotinylation efficiency (2mM) using ImageJ. This figure supports Figure 4.

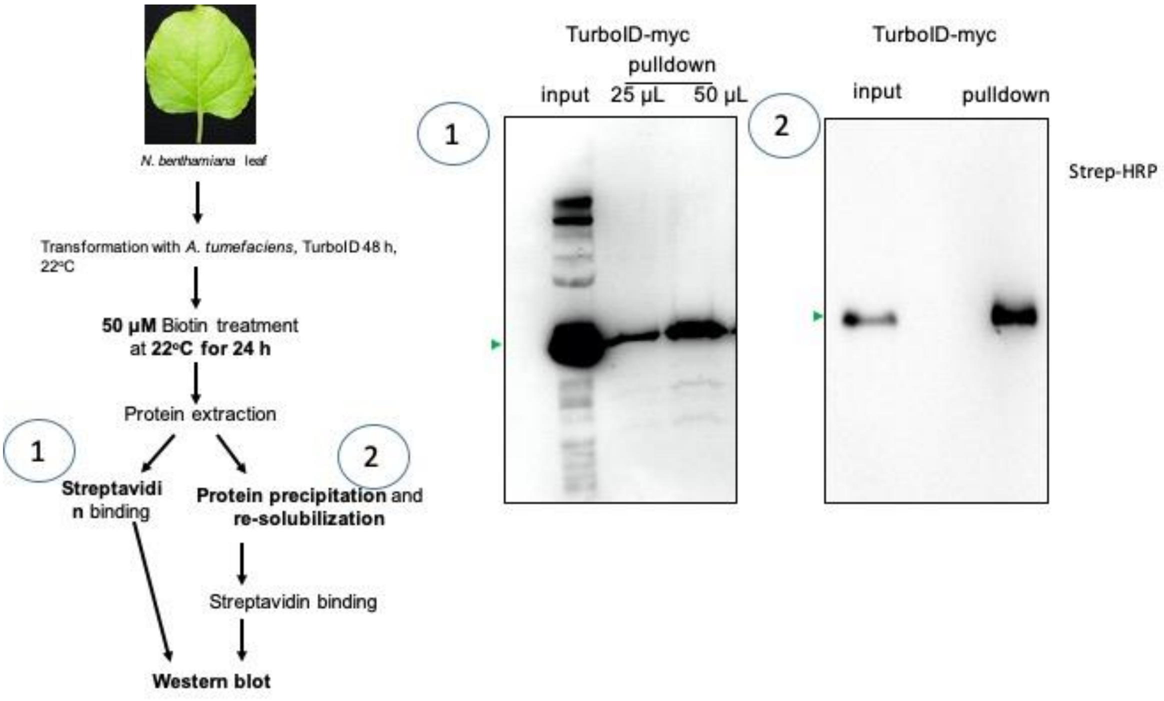


**Supplemental Figure 8. Exogenous application of biotin can exceed the binding capacity of streptavidin beads.** Blot on the left: input and IP with streptavidin using 25 or 50 µl of beads. Note that 2x more beads increased the recovery of the input signal, suggesting that the beads are saturated. Blot on the right: IP with 25 µl of streptavidin beads but in this case the supernatant was precipitated using ammonium acetate to remove excess biotin (in this case, the pulldown is much more than the input). Green arrowheads mark the position of the BL. This figure supports Figure 4.

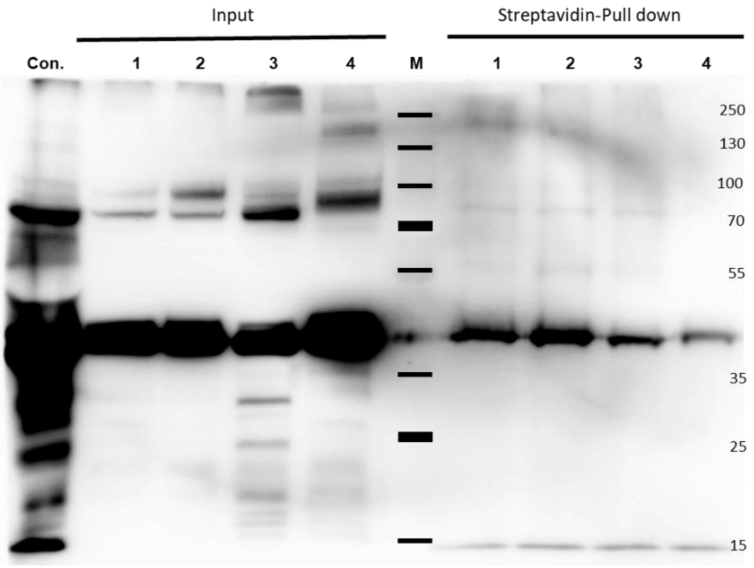


**Supplemental Figure 9. The biotin-streptavidin interaction is retained under harsh conditions.** Different extraction buffers were used for testing the binding affinity of biotin-labelled proteins with streptavidin from equal amounts of plant protein material: 1. 50mM HEPES, 150 mM NaCl, 0.5% NP40,

10% Glycerol, 1mM PMSF; 2. 50mM Tris/HCl, 150 mM NaCl, 0.5% NP40, 10% Glycerol, 1mM PMSF; 3. 1x PBST, 1mM PMSF; 4. 2x Laemmli sample buffer (65 mM Tris-HCl, pH 6.8, 20% (w/v) glycerol, 2% SDS, 0.01% bromophenol blue, 10mM DTT). Con: control. M: marker. Note that the lack of enrichment in the pulldown (right panel) is due to the presence of biotin (see also Supplemental Figure 8). This figure supports Figure 4, Figure 5 and Figure 6.

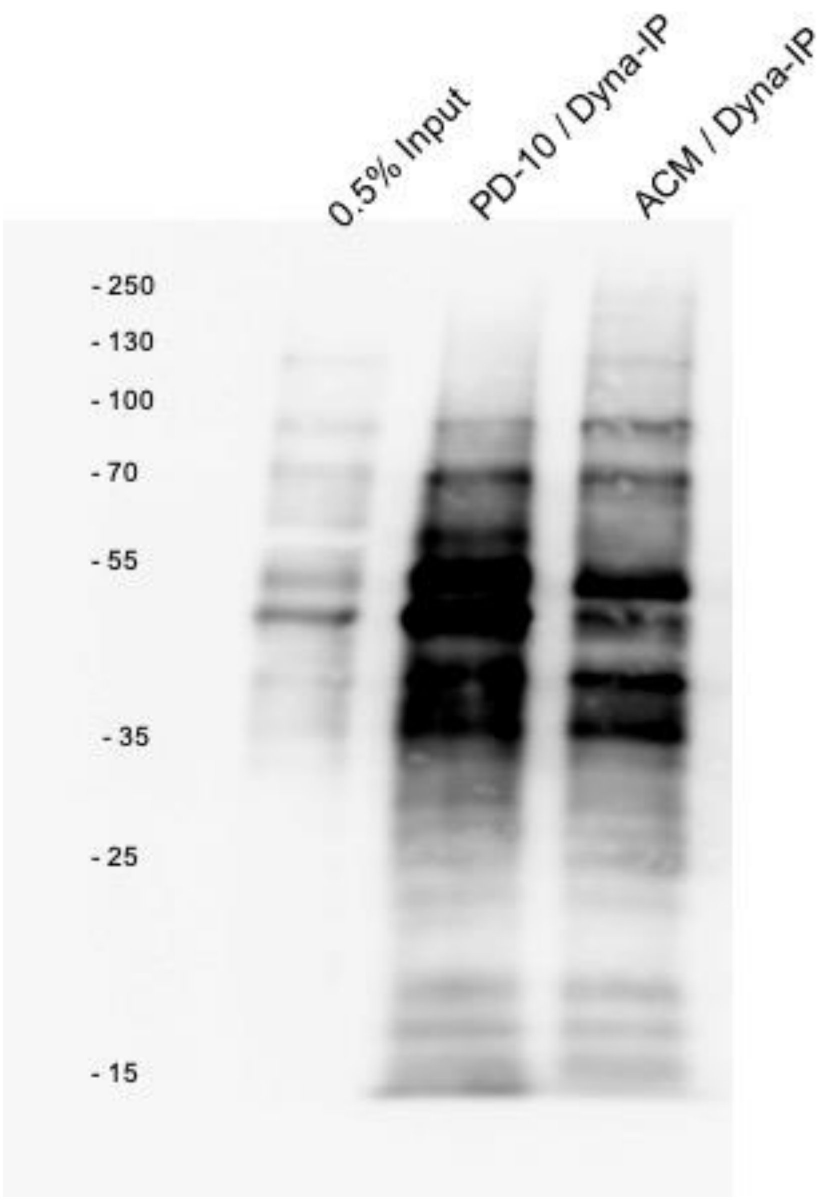


**Supplemental Figure 10. Comparing PD10 versus ammonium acetate for biotin removal.** To test the best way for removing free biotin following sample extraction, the same amount of supernatant (2.5 mL) was either de-salted using PD-10 columns or precipitated by ammonium acetate method (ACM). Next, the PD10-cleared lysate and the ammonium acetate re-dissolved precipitate were incubated with Dynabeads (Dyna-IP) to capture biotinylated proteins. Western blot analysis using anti-streptavidin showed that majority of biotinylated proteins (ca. 15 to 70 kDa) were better enriched in the supernatant by using PD-10 columns. Larger proteins (ca. >100 kDa) were better enriched using ACM.

**Supplemental Table1. List of expression vectors used in this study.**

| **Name** | **Vector** | **Promoter** | **Gene** | **PBL used** | **Material used** |
|---|---|---|---|---|---|
| GFP-BioID | pK7m34Gw | p35s | GFP | BioID | *Arabidopsis* cell culture |
| GFP-BioID2 | pK7m34Gw | p35s | GFP | BioID2 | *Arabidopsis* cell culture |
| TPLATE-BioID | pK7m34Gw | p35s | TPLATE | BioID | *Arabidopsis* cell culture |
| TPLATE-BioID2 | pH7m34Gw-R | p35s | TPLATE | BioID2 | *Arabidopsis* cell culture |
| TPLATE-linkerBioID | pH7m34Gw-R | p35s | TPLATE | $(GGGGS)_{13}$ BioID | *Arabidopsis* cell culture |
| TPLATE-linkerBioID2 | pH7m34Gw-R | p35s | TPLATE | $(GGGGS)_{13}$ BioID | *Arabidopsis* cell culture |
| GFP linkerTurboID | pK7m34Gw | p35s | GFP | $(GGGGS)_{13}$ TurboID | *Arabidopsis* cell culture |
| TPLATE-linkerTurboID | pK7m34Gw | p35s | TPLATE | $(GGGGS)_{13}$ TurboID | *Arabidopsis* cell culture |
| BirA-myc | pICSL86900 | p35s | BirA-myc | BirA-myc | *N.benthamiana* |
| BioID-myc | pICSL86900 | p35s | BioID-myc | BioID-myc | *N.benthamiana* |
| HF-BioID2-HA | pICSL86900 | p35s | BioID2-HA | BioID2-HA | *N.benthamiana* |
| GFP-TurboID-HF | pICSL86922 | p35s | TurboID-HF | TurboID | *N.benthamiana* |
| GFP-TurboID-His | Xpre2-S (pCAMBIA) | p35s | GFP | TurboID | *N.benthamiana* |
| NFR5-TurboID | Xpre2-S (pCAMBIA) | p35s | NFR5 | TurboID | *N.benthamiana* |
| GFP-LTI6b | Xpre2-S (pCAMBIA) | p35s | LTI6b | | N.benthamiana |
| BiFC_CC_TPLATE_BIN2 | BiFC_CC | p35s | TPLATE/BIN2 | NA | *N.benthamiana* |
| BiFC_CC_TPLATE_TOL9 | BiFC_CC | p35s | TPLATE/TOL9 | NA | *N.benthamiana* |
| BiFC_CN_TPLATE_TOL9 | BiFC_CN | p35s | TPLATE/TOL9 | NA | *N.benthamiana* |
| BiFC_CC_TPLATE_SCAMP5 | BiFC_CC | p35s | TPLATE/SCAMP5 | NA | *N.benthamiana* |
| BiFC_CN_TPLATE_SCAMP5 | BiFC_CN | p35s | TPLATE/SCAMP5 | NA | *N.benthamiana* |
| GFP-GGGS-BioID-FLAG | pK7m34Gw | p35s | GFP | BioID | *Solanum lycopersicum* stable hairy root lines & *N. Benthamiana* |
| GFP-GGGS-BioID2-FLAG | pK7m34Gw | p35s | GFP | BioID2 | *Solanum lycopersicum* stable hairy root lines & *N. Benthamiana* |
| GFP-GGGS-TurboID-FLAG | pK7m34Gw | p35s | GFP | TurboID | *Solanum lycopersicum* stable hairy root lines & *N. Benthamiana* |
| GFP-GGGGS-mTurbo-FLAG | pK7m34Gw | p35s | GFP | mTurbo | *Solanum lycopersicum* stable hairy root lines & *N. Benthamiana* |

**Supplemental Table 2: list of primers used.**

| Name | Primer |
|---|---|
| BirA | GTGGTCTC A T TCG GGA AAC GCGGCT ATT AGA TCA AAG GAT AAC ACC GTG CCA CTT A |
| BirA | GTGGTCTC A AAGC CTA CAG ATC CTC TTC TGA GAT GAG TTT TTG TTC TTT TTC TGC ACT ACG AAG G |
| BirA | GTGGTCTC A CGTAGAGGTCGTAAATGGTTTTC |
| BirA | GTGGTCTC C TACGTCCACGACCAGCCTGCT |
| HF-Module-Fw | GTGGTCTC ACCATGGGTTCCGGAAGAGGATCGCA |
| HF-Module-Rv | GTGGTCTC ACATTCCCTTGTCATCGTCATCCTTG |
| BioID2 with long-linker-Module-Fw | agGAAGACaaTTCGGGATCTGGAGGTGGCGGAAG |
| BioID2 without linker-Module-Fw | agGAAGACaaTTCGTTTAAGAACTTGATATGGCTGAAAG |
| ScFv-superfolder-GFP-Module-Fw | GTGGTCTC A ATGG GCCCCGACATCGT |
| ScFv-superfolder-GFP-Module-Fw | GTGGTCTC A CGAA CCACCTTTGTAGAGCTC |
| NotImcherryu | GCGGCCGCATGGTGAGCAAGGGCGAGGAG |
| NotImcherryd | GCGGCCGCTTACTTGTACAGCTCGTCC ATG |
| attB1_SCAMP5_FWD | GGGGACAAGTTTGTACAAAAAAGCAGGCTTAATGAATCGCCACCACGATCCCAATCC |
| attB4_SCAMP5_REV | GGGGACAACTTTGTATAGAAAAGTTGGGTGCTTGTTTCCCCTAAAGTAGAGGTAAATTTTCTG |
| attB3_TPLATE_FWD | GGGGACAACTTTGTATAATAAAGTTGTAATGGACATTCTTTTTGCTCAGATCCAGGC |
| attB2-TPLATE_REV | GGGGACCACTTTGTACAAGAAAGCTGGGTTGTTAACTTTGGTATATTTTCTATCTTTGCACAAGATG |
| attB1_TOL9_FWD | GGGGACAAGTTTGTACAAAAAAGCAGGCTTAATGGTGAACGCTATGG |
| attB4_TOL9_REV | GGGGACAACTTTGTATAGAAAAGTTGGGTGTCACATGGTACCAGCTC |

**Supplemental Table 3: Cell cultures expressing different TPLATE-PBLs identifies TPC subunits with different amount of non-biotinylated peptides.** The table shows the amount of non-biotinylated peptides identified for TPC subunits in GFP and TPLATE-BioIDs PBL cell cultures. Preparation of samples for LC-MS/MS involved the use of a buffer containing 8M Urea and 2% SDS. Cell cultures were incubated with either 50µM or 2mM biotin for 24 hours before harvesting. This table shows higher amount of peptides detected for all TPC subunits in case of linkerTurboID constructs than other BioIDs PBL. Also, it shows higher amounts of peptides detected in TurboID control experiments (GFP-linkerTurboID vs. GFP-BioID and GFP-BioID2) suggesting increased promiscuity of TurboID. This table supports Figure 4.

| Total number of unique non-biotinylated peptides identified for each TPC subunit | | | | | | | |
|---|---|---|---|---|---|---|---|
| | AT1G20760 | AT1G21630 | AT2G07360 | AT3G01780 | AT3G50590 | AT5G24710 | AT5G57460 |
| | AtEH1 | AtEH2 | TASH3 | TPLATE | TWD40-1 | TWD40-2 | TML |
| **TPLATE linkerTurboID** | 73 | 155 | 205 | 384 | 228 | 374 | 116 |
| **GFP linkerTurboID** | 40 | 98 | 122 | 27 | 102 | 271 | 21 |
| **TPLATE BioID** | 20 | 32 | 2 | 428 | 68 | 95 | 37 |
| **TPLATE linkerBioID** | 19 | 37 | 15 | 540 | 35 | 34 | 16 |
| **GFP BioID** | - | - | - | - | - | 26 | - |
| **TPLATE BioID2** | 9 | 30 | - | 157 | 42 | 38 | 51 |
| **TPLATE linkerBioID2** | 24 | 55 | 59 | 453 | 107 | 89 | 38 |
| **GFP BioID2** | - | - | 6 | - | - | 42 | - |
| **TPLATE linkerBioID 2mM** | - | - | - | 10 | - | - | - |

**Supplemental Table 4. Full list of significantly enriched identifications with TPLATE-BioID versus GFP-BioID at different incubation temperatures for 24 hours with 50 µM biotin.**
The table shows the amount of non-biotinylated peptides identified for TPC subunits in GFP and TPLATE-BioID PBLs at different temperatures (25°, 28°, 30° and 35°). Cell cultures were incubated with 50µM biotin for 24 hours. This table supports figure 3.

**Supplemental Table 5. Full list of significantly enriched identifications with TPLATE as bait using BioID, BioID2, linkerBioID, linkerBioID2, linkerTurboID E1, linkerTurboID E2 and linkerTurboID E1+E2, versus the respective GFP PBLs.** This table shows the amount of non-biotinylated peptides identified for TPC subunits in GFP and various TPLATE-PBLs. Cell cultures were treated with either 50 µM or 2mM biotin at 28°C for 24 hours. This table supports Figure 4.

**Supplemental Table 6. Full list of significantly enriched hits with either TPLATE PBLs or GSrhino PD, including average iBAQ values normalized versus TPLATE.** The table shows the normalized MaxQuant iBAQ values of all proteins that were significantly enriched in TPLATE-GSRhino (PD), TPLATE-linkerBioID (LBioID), TPLATE-linkerBioID2 (LBioID2) or TPLATE-linkerTurboID (LTurboID) compared to the respective control samples. Colored cells (red for PD, orange for LBioID, green for

LBioID2 and blue for LTurboID) are proteins that are significantly enriched for those experiments. This table supports Figure 5.

**Supplemental Table 7. Full list of significantly enriched hits with TPLATE-linkerTurboID, including average iBAQ values normalized versus TPLATE, at different incubation times at 25 degrees and 24-hour incubation time at 28 degrees, all with 50 µM biotin.** The table shows the normalized MaxQuant iBAQ values of all proteins that were significantly enriched in TPLATE-linkerTurboID (LTurboID) at different temperatures (25°C or 28°C) and biotin treatments (10 mins, 1 hour, 6 hours and 24 hours) compared to the respective control samples. Colored cells (red for 25°C,10 mins, orange for 25°C,1 hour, green for 25°C, 6 hours and blue triangle for 25°C, 24 hours and dark blue circle for 28°C, 24 hours) are proteins that are significantly enriched for those experiments. This table supports Figure 6.

**Supplemental Table 8. MaxQuant TPC subunits peptides in TPLATE-LTurboID.** The table shows the amount of biotinylated and non-biotinylated peptides identified for TPC subunits in the TPLATE-linkerTurboID culture. The cell culture was treated with 50 µM biotin at 28°C for 24 hours. This table supports Figure 7.

**Supplemental sequences. List of all used PBL sequences.**

AttR2_**BioID-**3xHA_attL3

AttR2_GACAAGGACAACACCGTACCCCTGAAGCTCATCGCACTACTTGCTAATGGTGAATTTCACTCCGGGGAA
CAGCTAGGAGAAACACTCGGCATGTCAAGAGCAGCAATAAACAAGCATATACAAACACTCCGTGACTGGGGCGTG
GATGTATTTACCGTCCCTGGTAAGGGATACTCCCTCCCGGAACCAATCCAACTTCTCAATGCCAAGCAGATTCTA
GGTCAACTCGATGGTGGCTCAGTGGCGGTTTTACCAGTAATAGACTCTACGAACCAGTATCTGCTCGACCGAATA
GGGGAGCTTAAATCCGGAGATGCTTGCATTGCGGAATATCAACAAGCCGGACGTGGTGGCAGGGGAAGGAAGTGG
TTCAGTCCCTTCGGCGCGAATCTCTATCTGAGCATGTTCTGGCGACTTGAACAGGGTCCCGCGGCCGCTATCGGC
CTATCCTTAGTTATAGGTATCGTCATGGCTGAGGTCCTCAGAAAGCTAGGGGCTGACAAGGTGAGGGTAAAATGG
CCTAATGACCTCTATCTTCAAGATAGAAAGTTGGCCGGTATCTTAGTGGAATTAACGGGAAAGACCGGCGATGCT
GCACAGATCGTGATCGGTGCGGGAATTAACATGGCCATGCGACGAGTGGAAGAAAGCGTTGTTAATCAGGGATGG
ATTACGCTACAAGAAGCCGGTATAAATCTTGACCGTAACACCCTAGCAGCCATGCTGATCCGTGAACTACGAGCG
GCGTTGGAACTTTTTGAACAGGAGGGTTTGGCACCTTACCTTAGCCGATGGGAAAAACTCGACAACTTCATTAAC
CGTCCCGTCAAATTGATTATTGGCGATAAGGAAATTTTCGGAATAAGTAGAGGGATCGATAAGCAGGGAGCGTTA
CTACTTGAGCAGGATGGTATCATAAAGCCGTGGATGGGAGGCGAAATTTCACTCCGTAGCGCCGAAAAGGCCTAC
CCATACGATGTTCCAGATTACGCTTACCCATACGATGTTCCAGATTACGCTTACCCATACGATGTTCCAGATTAC
GCTTGA_attL3.

AttR2_**BioID2-**3xHA_attL3

AttR2_GACTTCAAAAACTTAATTTGGTTAAAAGAAGTGGATTCAACACAGGAGCGACTTAAAGAATGGAATGTA
AGTTACGGCACGGCATTAGTTGCCGATCGTCAAACGAAGGGAAGAGGAGGGTTGGGGCGTAAGTGGCTAAGTCAG
GAAGGTGGGCTATACTTCAGTTTTCTTCTGAATCCGAAGGAGTTCGAGAACCTGCTACAATTACCGCTCGTACTG
GGCCTGAGTGTGAGCGAGGCTTTAGAAGAAATTACAGAAATCCCCTTCAGTTTGAAGTGGCCAAATGATGTCTAT
TTTCAGGAAAAGAAAGTCTCTGGCGTACTGTGCGAACTATCAAAGGATAAGCTGATCGTAGGAATCGGTATAAAT
GTTAATCAACGAGAGATCCCGGAGGAAATCAAAGACAGAGCAACCACGTTATATGAGATTACCGGGAAGGACTGG
GATAGGAAAGAGGTACTATTAAAGGTACTAAAGCGAATATCAGAAAATCTCAAGAAATTCAAGGAAAAAAGCTTC
AAAGAATTCAAAGGCAAGATTGAAAGCAAGATGCTATACCTCGGGGAAGAGGTTAAGTTGCTTGGGGAAGGAAAA
ATTACTGGTAAGCTGGTTGGCCTGAGCGAAAAGGGCGGTGCTCTCATTTTAACTGAGGAAGGATCAAAGAAATC
TTGAGCGGAGAGTTCTCCTTGCGTCGAAGCGCCTACCCATACGATGTTCCAGATTACGCTTACCCATACGATGTT
CCAGATTACGCTTACCCATACGATGTTCCAGATTACGCTTGA_attL3.

AttR2_linker**BioID-**3xHA_attL3

AttR2_GGTGGCGGAGGTTCTGGCGGTGGAGGATCCGGAGGTGGCGGGTCCGGTGGTGGCGGATCAGGTGGAGGA
GGTTCGGGAGGCGGAGGTAGTGGAGGCGGTGGGAGCGGCGGAGGTGGGAGCGGTGGCGGAGGGAGTGGAGGTGGA
GGCTCGGGCGGAGGTGGCTCCGGCGGAGGTGGGTCTGGAGGCGGTGGCTCGGACAAGGACAACACCGTACCCCTG
AAGCTCATCGCACTACTTGCTAATGGTGAATTTCACTCCGGGGAACAGCTAGGAGAAACACTCGGCATGTCAAGA
GCAGCAATAAACAAGCATATACAAACACTCCGTGACTGGGGCGTGGATGTATTTACCGTCCCTGGTAAGGGATAC
TCCCTCCCGGAACCAATCCAACTTCTCAATGCCAAGCAGATTCTAGGTCAACTCGATGGTGGCTCAGTGGCGGTT
TTACCAGTAATAGACTCTACGAACCAGTATCTGCTCGACCGAATAGGGGAGCTTAAATCCGGAGATGCTTGCATT
GCGGAATATCAACAAGCCGGACGTGGTGGCAGGGGAAGGAAGTGGTTCAGTCCCTTCGGCGCGAATCTCTATCTG
AGCATGTTCTGGCGACTTGAACAGGGTCCCGCGGCCGCTATCGGCCTATCCTTAGTTATAGGTATCGTCATGGCT
GAGGTCCTCAGAAAGCTAGGGGCTGACAAGGTGAGGGTAAAATGGCCTAATGACCTCTATCTTCAAGATAGAAAG
TTGGCCGGTATCTTAGTGGAATTAACGGGAAAGACCGGCGATGCTGCACAGATCGTGATCGGTGCGGGAATTAAC
ATGGCCATGCGACGAGTGGAAGAAAGCGTTGTTAATCAGGGATGGATTACGCTACAAGAAGCCGGTATAAATCTT
GACCGTAACACCCTAGCAGCCATGCTGATCCGTGAACTACGAGCGGCGTTGGAACTTTTTGAACAGGAGGGTTTG
GCACCTTACCTTAGCCGATGGGAAAAACTCGACAACTTCATTAACCGTCCCGTCAAATTGATTATTGGCGATAAG
GAAATTTTCGGAATAAGTAGAGGGATCGATAAGCAGGGAGCGTTACTACTTGAGCAGGATGGTATCATAAAGCCG
TGGATGGGAGGCGAAATTTCACTCCGTAGCGCCGAAAAGGCCTACCCATACGATGTTCCAGATTACGCTTACCCA
TACGATGTTCCAGATTACGCTTACCCATACGATGTTCCAGATTACGCTTGA_attL3.

AttR2_linker**BioID2-**3xHA_attL3

AttR2_GGTGGCGGAGGTTCTGGCGGTGGAGGATCCGGAGGTGGCGGGTCCGGTGGTGGCGGATCAGGTGGAGGA
GGTTCGGGAGGCGGAGGTAGTGGAGGCGGTGGGAGCGGCGGAGGTGGGAGCGGTGGCGGAGGGAGTGGAGGTGGA
GGCTCGGGCGGAGGTGGCTCCGGCGGAGGTGGGTCTGGAGGCGGTGGCTCGGACTTCAAAAACTTAATTTGGTTA
AAAGAAGTGGATTCAACACAGGAGCGACTTAAAGAATGGAATGTAAGTTACGGCACGGCATTAGTTGCCGATCGT
CAAACGAAGGGAAGAGGAGGGTTGGGGCGTAAGTGGCTAAGTCAGGAAGGTGGGCTATACTTCAGTTTTCTTCTG
AATCCGAAGGAGTTCGAGAACCTGCTACAATTACCGCTCGTACTGGGCCTGAGTGTGAGCGAGGCTTTAGAAGAA
ATTACAGAAATCCCCTTCAGTTTGAAGTGGCCAAATGATGTCTATTTTCAGGAAAAGAAAGTCTCTGGCGTACTG
TGCGAACTATCAAAGGATAAGCTGATCGTAGGAATCGGTATAAATGTTAATCAACGAGAGATCCCGGAGGAAATC
AAAGACAGAGCAACCACGTTATATGAGATTACCGGGAAGGACTGGGATAGGAAAGAGGTACTATTAAAGGTACTA
AAGCGAATATCAGAAAATCTCAAGAAATTCAAGGAAAAAGCTTCAAAGAATTCAAAGGCAAGATTGAAAGCAAG
ATGCTATACCTCGGGGAAGAGGTTAAGTTGCTTGGGGAAGGAAAAATTACTGGTAAGCTGGTTGGCCTGAGCGAA
AAGGGCGGTGCTCTCATTTTAACTGAGGAAGGGATCAAAGAAATCTTGAGCGGAGAGTTCTCCTTGCGTCGAAGC
GCCTACCCATACGATGTTCCAGATTACGCTTACCCATACGATGTTCCAGATTACGCTTACCCATACGATGTTCCA
GATTACGCTTGA_attL3.

AttR2_linker**TurboID-**3xHA _attL3

AttR2_GGTGGCGGAGGTTCTGGCGGTGGAGGATCCGGAGGTGGCGGGTCCGGTGGTGGCGGATCAGGTGGAGGA
GGTTCGGGAGGCGGAGGTAGTGGAGGCGGTGGGAGCGGCGGAGGTGGGAGCGGTGGCGGAGGGAGTGGAGGTGGA
GGCTCGGGCGGAGGTGGCTCCGGCGGAGGTGGGTCTGGAGGCGGTGGCTCGGACAAAGACAATACCGTGCCCCTC
AAACTAATTGCATTGCTTGCCAATGGTGAGTTCCATTCTGGGGAACAACTTGGGGAGACTCTGGGCATGTCCCGA
GCGGCCATTAATAAGCACATACAGACTTTGAGGGACTGGGGTGTCGATGTGTTTACAGTACCGGGTAAGGGATAT
TCTCTGCCGGAGCCGATACCTTTACTCAATGCAAAGCAGATTTTAGGACAGCTGGATGGCGGGAGCGTGGCTGTC
CTCCCGGTGGTCGATAGTACAAATCAATATCTTTTGGACAGGATCGGTGAGCTGAAATCTGGAGATGCTTGCATA
GCGGAATACCAACAAGCTGGTAGGGGATCTCGAGGCCGAAAGTGGTTTTCACCCTTTGGGGCCAACTTATACTTA
TCCATGTTCTGGCGTCTGAAACGTGGCCCGGCAGCCATAGGGCTAGGGCCCGTAATTGGGATCGTTATGGCGGAA
GCGTTACGAAAGTTGGGAGCCGATAAAGTCAGAGTAAAGTGGCCAAATGATCTTTATTTACAAGATAGAAAGCTG
GCGGGCATTCTTGTTGAGCTTGCCGGAATAACAGGCGATGCCGCACAGATCGTTATAGGGGCCGGGATTAATGTG
GCTATGAGGAGGGTCGAAGAAAGTGTTGTGAATCAAGGATGGATTACTCTCCAAGAGGCGGGTATAAATCTTGAT
CGAAATACTCTAGCTGCAACCTTGATACGAGAGCTAAGAGCAGCACTAGAATTGTTTGAACAAGAAGGGCTGGCC
CCATATCTCCCTCGATGGGAGAAGTTGGACAATTTTATAAACAGACCGGTGAAGTTAATCATTGGAGACAAAGAA
ATTTTCGGTATAAGCCGTGGGATCGATAAACAGGGTGCTCTTCTCCTTGAGCAGGACGGTGTCATCAAGCCTTGG
ATGGGAGGCGAAATCTCTCTCCGAAGCGCCGAGAAGAAGCTTGCCTACCCATACGATGTTCCAGATTACGCTTAC
CCATACGATGTTCCAGATTACGCTTACCCATACGATGTTCCAGATTACGCT TGA_attL3.

attL4-*omegaleader*-start-3xHA-TurboID-(GGGGS) $_{13}$ linker-attR1

AAATAATGATTTTATTTTGACTGATAGTGACCTGTTCGTTGCAACAAATTGATAAGCAATGCTTTTTTATAATGC
CAACTTTGTATAGAAAAGTTG*AACAACAAACAACAAACAACATTACAATTACA*ATGTACCCATACGATGTTCCAG
ATTACGCTTACCCATACGATGTTCCAGATTACGCTTACCCATACGATGTTCCAGATTACGCTGCCAAAGACAATA
CCGTGCCCCTCAAACTAATTGCATTGCTTGCCAATGGTGAGTTCCATTCTGGGGAACAACTTGGGGAGACTCTGG
GCATGTCCCGAGCGGCCATTAATAAGCACATACAGACTTTGAGGGACTGGGGTGTCGATGTGTTTACAGTACCGG
GTAAGGGATATTCTCTGCCGGAGCCGATACCTTTACTCAATGCAAAGCAGATTTTAGGACAGCTGGATGGCGGGA
GCGTGGCTGTCCTCCCGGTGGTCGATAGTACAAATCAATATCTTTTGGACAGGATCGGTGAGCTGAAATCTGGAG
ATGCTTGCATAGCGGAATACCAACAAGCTGGTAGGGGATCTCGAGGCCGAAAGTGGTTTTCACCCTTTGGGGCCA
ACTTATACTTATCCATGTTCTGGCGTCTGAAACGTGGCCCGGCAGCCATAGGGCTAGGGCCCGTAATTGGGATCG
TTATGGCGGAAGCGTTACGAAAGTTGGGAGCCGATAAAGTCAGAGTAAAGTGGCCAAATGATCTTTATTTACAAG
ATAGAAAGCTGGCGGGCATTCTTGTTGAGCTTGCCGGAATAACAGGCGATGCCGCACAGATCGTTATAGGGGCCG
GGATTAATGTGGCTATGAGGAGGGTCGAAGAAAGTGTTGTGAATCAAGGATGGATTACTCTCCAAGAGGCGGGTA
TAAATCTTGATCGAAATACTCTAGCTGCAACCTTGATACGAGAGCTAAGAGCAGCACTAGAATTGTTTGAACAAG
AAGGGCTGGCCCCATATCTCCCTCGATGGGAGAAGTTGGACAATTTTATAAACAGACCGGTGAAGTTAATCATTG
GAGACAAAGAAATTTTCGGTATAAGCCGTGGGATCGATAAACAGGGTGCTCTTCTCCTTGAGCAGGACGGTGTCA
TCAAGCCTTGGATGGGAGGCGAAATCTCTCTCCGAAGCGCCGAGAAGAAGCTTGACGGTGGCGGAGGTTCTGGCG
GTGGAGGATCCGGAGGTGGCGGGTCCGGTGGTGGCGGATCAGGTGGAGGAGGTTCGGGAGGCGGAGGTAGTGGAG
GCGGTGGGAGCGGCGGAGGTGGGAGCGGTGGCGGAGGGAGTGGAGGTGGAGGCTCGGGCGGAGGTGGCTCCGGCG
GAGGTGGGTCTGGAGGCGGTGGCTCGGCAAGTTTGTACAAAAAAGTTGAACGAGAAACGTAAAATGATATAAATA
TCAATATATTAAATTAGATTTTGCATAAAAAACAGACTACATAATACTGTAAAACACAACATATGCAGTCACTAT
G

AttR2_GGGGS-**BioID**-FLAG_attL3

AttR2_GGAGGCGGTGGATCGAAGGACAACACCGTGCCCCTGAAGCTGATCGCCCTGCTGGCCAACGGCGAGTTC
CACAGCGGCGAGCAGCTGGGCGAGACCCTGGGCATGAGCCGGGCCGCCATCAACAAGCACATCCAGACCCTGCGG
GACTGGGGCGTGGACGTGTTCACCGTGCCCGGCAAGGGCTACAGCCTGCCCGAGCCCATCCAGCTGCTGAACGCC
AAGCAGATCCTGGGCCAGCTGGACGGCGGCAGCGTGGCCGTGCTGCCCGTGATCGACAGCACCAACCAGTACCTG
CTGGACCGGATCGGCGAGCTGAAGAGCGGCGACGCCTGCATCGCCGAGTACCAGCAGGCCGGCCGGGGCGGCCGG
GGCCGGAAGTGGTTCAGCCCCTTCGGCGCCAACCTGTACCTGAGCATGTTCTGGCGGCTGGAGCAGGGCCCCGCC
GCCGCCATCGGCCTGAGCCTGGTGATCGGCATCGTGATGGCCGAGGTGCTGCGGAAGCTGGGCGCCGACAAGGTG
CGGGTGAAGTGGCCCAACGACCTGTACCTGCAGGACCGGAAGCTGGCCGGCATCCTGGTGGAGCTGACCGGCAAG
ACCGGCGACGCCGCCCAGATCGTGATCGGCGCCGGCATCAACATGGCCATGCGGCGGGTGGAGGAGAGCGTGGTG
AACCAGGGCTGGATCACCCTGCAGGAGGCCGGCATCAACCTGGACCGGAACACCCTGGCCGCCATGCTGATCCGG
GAGCTGCGGGCCGCCCTGGAGCTGTTCGAGCAGGAGGGCCTGGCCCCCTACCTGAGCCGGTGGGAGAAGCTGGAC
AACTTCATCAACCGGCCCGTGAAGCTGATCATCGGCGACAAGGAGATCTTCGGCATCAGCCGGGGCATCGACAAG
CAGGGCGCCCTGCTGCTGGAGCAGGACGGCATCATCAAGCCCTGGATGGGCGGCGAGATCAGCCTGCGGAGCGCC
GAGAAGGACTACAAAGACGATGACGATAAATGA_attL3

AttR2_GGGGS-**BioID2**-FLAG_attL3

AttR2_GGAGGCGGTGGATCGTTCAAGAACCTGATCTGGCTGAAGGAGGTGGACAGCACCCAGGAGAGACTGAAG
GAGTGGAACGTGAGCTACGGCACCGCCCTGGTGGCCGACAGACAGACCAAGGGCAGAGGCGGCCTGGGCAGAAAG
TGGCTGAGCCAGGAGGGCGGCCTGTACTTCAGCTTCCTGCTGAACCCCAAGGAGTTCGAGAACCTGCTGCAGCTG
CCCCTGGTGCTGGGCCTGAGCGTGAGCGAGGCCCTGGAGGAGATCACCGAGATCCCCTTCAGCCTGAAGTGGCCC
AACGACGTGTACTTCCAGGAGAAGAAGGTGAGCGGCGTGCTGTGCGAGCTGAGCAAGGACAAGCTGATCGTGGGC
ATCGGCATCAACGTGAACCAGAGAGAGATCCCCGAGGAGATCAAGGACAGAGCCACCACCCTGTACGAGATCACC
GGCAAGGACTGGGACAGAAAGGAGGTGCTGCTGAAGGTGCTGAAGAGAATCAGCGAGAACCTGAAGAAGTTCAAG
GAGAAGAGCTTCAAGGAGTTCAAGGGCAAGATCGAGAGCAAGATGCTGTACCTGGGCGAGGAGGTGAAGCTGCTG
GGCGAGGGCAAGATCACCGGCAAGCTGGTGGGCCTGAGCGAGAAGGGCGGCGCCCTGATCCTGACCGAGGAGGGC
ATCAAGGAGATCCTGAGCGGCGAGTTCAGCCTGAGAAGAAGCGACTACAAAGACGATGACGATAAATGA_attL3

AttR2_GGGGS-**TurboID**-FLAG_attL3

AttR2_GGAGGCGGTGGATCGAAAGACAATACTGTGCCTCTGAAGCTGATCGCTCTCCTGGCTAATGGCGAGTTC
CATAGTGGCGAACAGCTGGGAGAAACCCTGGGCATGTCCAGGGCCGCTATCAACAAGCACATTCAGACTCTGCGC
GACTGGGGCGTGGACGTGTTCACCGTGCCCGGAAAGGGCTACTCTCTGCCCGAGCCTATCCCGCTGCTGAACGCT
AAACAGATTCTGGGACAGCTGGACGGCGGGAGCGTGGCAGTCCTGCCTGTGGTCGACTCCACCAATCAGTACCTG
CTGGATCGAATCGGCGAGCTGAAGAGTGGGGATGCTTGCATTGCAGAATATCAGCAGGCAGGGAGAGGAAGCAGA
GGGAGGAAATGGTTCTCTCCTTTTGGAGCTAACCTGTACCTGAGTATGTTTTGGCGCCTGAAGCGGGGACCAGCA
GCAATCGGCCTGGGCCCGGTCATCGGAATTGTCATGGCAGAAGCGCTGCGAAAGCTGGGAGCAGACAAGGTGCGA

GTCAAATGGCCCAATGACCTGTATCTGCAGGATAGAAAGCTGGCAGGCATCCTGGTGGAGCTGGCCGGAATAACA
GGCGATGCTGCACAGATCGTCATTGGCGCCGGGATTAACGTGGCTATGAGGCGCGTGGAGGAAAGCGTGGTCAAT
CAGGGCTGGATCACACTGCAGGAAGCAGGGATTAACCTGGACAGGAATACTCTGGCCGCTACGCTGATCCGAGAG
CTGCGGGCAGCCCTGGAACTGTTCGAGCAGGAAGGCCTGGCTCCATATCTGCCACGGTGGGAGAAGCTGGATAAC
TTCATCAATAGACCCGTGAAGCTGATCATTGGGGACAAAGAGATTTTCGGGATTAGCCGGGGGATTGATAAACAG
GGAGCCCTGCTGCTGGAACAGGACGGAGTTATCAAACCCTGGATGGGCGGAGAAATCAGTCTGCGGTCTGCCGAA
AAGGACTACAAAGACGATGACGATAAATGA_attL3

AttR2_GGGGS-**miniTurboID**-FLAG_attL3

AttR2_GGAGGCGGTGGATCGATCCCGCTGCTGAACGCTAAACAGATTCTGGGACAGCTGGACGGCGGGAGCGTG
GCAGTCCTGCCTGTGGTCGACTCCACCAATCAGTACCTGCTGGATCGAATCGGCGAGCTGAAGAGTGGGGATGCT
TGCATTGCAGAATATCAGCAGGCAGGGAGAGGAAGCAGAGGGAGGAAATGGTTCTCTCCTTTTGGAGCTAACCTG
TACCTGAGTATGTTTTGGCGCCTGAAGCGGGGACCAGCAGCAATCGGCCTGGGCCCGGTCATCGGAATTGTCATG
GCAGAAGCGCTGCGAAAGCTGGGAGCAGACAAGGTGCGAGTCAAATGGCCCAATGACCTGTATCTGCAGGATAGA
AAGCTGGCAGGCATCCTGGTGGAGCTGGCCGGAATAACAGGCGATGCTGCACAGATCGTCATTGGCGCCGGGATT
AACGTGGCTATGAGGCGCGTGGAGGAAAGCGTGGTCAATCAGGGCTGGATCACACTGCAGGAAGCAGGGATTAAC
CTGGACAGGAATACTCTGGCCGCTATGCTGATCCGAGAGCTGCGGGCAGCCCTGGAACTGTTCGAGCAGGAAGGC
CTGGCTCCATATCTGTCACGGTGGGAGAAGCTGGATAACTTCATCAATAGACCCGTGAAGCTGATCATTGGGGAC
AAAGAGATTTTCGGGATTAGCCGGGGGATTGATAAACAGGGAGCCCTGCTGCTGGAACAGGACGGAGTTATCAAA
CCCTGGATGGGCGGAGAAATCAGTCTGCGGTCTGCCGAAAAGGACTACAAAGACGATGACGATAAATGA_attL3

B_TurboID_6xHIS_C

AAGAAGACTATACGGGTCTCATCTGAACAATGAAAGACAATACTGTGCCTCTGAAGCTGATCGCTCTCCTGGCTA
ATGGCGAGTTCCATAGTGGCGAACAGCTGGGAGAAACCCTGGGCATGTCCAGGGCCGCTATCAACAAGCACATTC
AGACTCTGCGCGACTGGGGCGTGGACGTGTTCACCGTGCCCGGAAAGGGCTACTCTCTGCCCGAGCCTATCCCGC
TGCTGAACGCTAAACAGATTCTGGGACAGCTGGACGGCGGGAGCGTGGCAGTCCTGCCTGTGGTCGACTCCACCA
ATCAGTACCTGCTGGATCGAATCGGCGAGCTGAAGAGTGGGGATGCTTGCATTGCAGAATATCAGCAGGCAGGGA
GAGGAAGCAGAGGGAGGAAATGGTTCTCTCCTTTTGGAGCTAACCTGTACCTGAGTATGTTTTGGCGCCTGAAGC
GGGGACCAGCAGCAATCGGCCTGGGCCCGGTCATCGGAATTGTCATGGCAGAAGCGCTGCGAAAGCTGGGAGCAG
ACAAGGTGCGAGTCAAATGGCCCAATGACCTGTATCTGCAGGATAGAAAGCTGGCAGGCATCCTGGTGGAGCTGG
CCGGAATAACAGGCGATGCTGCACAGATCGTCATTGGCGCCGGGATTAACGTGGCTATGAGGCGCGTGGAGGAAA
GCGTGGTCAATCAGGGCTGGATCACACTGCAGGAAGCAGGGATTAACCTGGACAGGAATACTCTGGCCGCTACGC
TGATCCGAGAGCTGCGGGCAGCCCTGGAACTGTTCGAGCAGGAAGGCCTGGCTCCATATCTGCCACGGTGGGAGA
AGCTGGATAACTTCATCAATAGACCCGTGAAGCTGATCATTGGGGACAAAGAGATTTTCGGGATTAGCCGGGGGA
TTGATAAACAGGGAGCCCTGCTGCTGGAACAGGACGGAGTTATCAAACCCTGGATGGGCGGAGAAATCAGTCTGC
GGTCTGCCGAAAAGAAGCTTGCGGCCGCACTCGAGCACCACCACCACCACCACGGCACCTGAGACCTCTGTAGTC
TTCTT

C_TurboID_6xHIS_D

AAGAAGACTATACGGGTCTCACACCATGAAAGACAATACTGTGCCTCTGAAGCTGATCGCTCTCCTGGCTAATGG
CGAGTTCCATAGTGGCGAACAGCTGGGAGAAACCCTGGGCATGTCCAGGGCCGCTATCAACAAGCACATTCAGAC
TCTGCGCGACTGGGGCGTGGACGTGTTCACCGTGCCCGGAAAGGGCTACTCTCTGCCCGAGCCTATCCCGCTGCT
GAACGCTAAACAGATTCTGGGACAGCTGGACGGCGGGAGCGTGGCAGTCCTGCCTGTGGTCGACTCCACCAATCA
GTACCTGCTGGATCGAATCGGCGAGCTGAAGAGTGGGGATGCTTGCATTGCAGAATATCAGCAGGCAGGGAGAGG
AAGCAGAGGGAGGAAATGGTTCTCTCCTTTTGGAGCTAACCTGTACCTGAGTATGTTTTGGCGCCTGAAGCGGGG
ACCAGCAGCAATCGGCCTGGGCCCGGTCATCGGAATTGTCATGGCAGAAGCGCTGCGAAAGCTGGGAGCAGACAA
GGTGCGAGTCAAATGGCCCAATGACCTGTATCTGCAGGATAGAAAGCTGGCAGGCATCCTGGTGGAGCTGGCCGG
AATAACAGGCGATGCTGCACAGATCGTCATTGGCGCCGGGATTAACGTGGCTATGAGGCGCGTGGAGGAAAGCGT
GGTCAATCAGGGCTGGATCACACTGCAGGAAGCAGGGATTAACCTGGACAGGAATACTCTGGCCGCTACGCTGAT
CCGAGAGCTGCGGGCAGCCCTGGAACTGTTCGAGCAGGAAGGCCTGGCTCCATATCTGCCACGGTGGGAGAAGCT
GGATAACTTCATCAATAGACCCGTGAAGCTGATCATTGGGGACAAAGAGATTTTCGGGATTAGCCGGGGGATTGA
TAAACAGGGAGCCCTGCTGCTGGAACAGGACGGAGTTATCAAACCCTGGATGGGCGGAGAAATCAGTCTGCGGTC
TGCCGAAAAGAAGCTTGCGGCCGCACTCGAGCACCACCACCACCACCACAAGGtGAGACCTCTGTAGTCTTCTT

D_TurboID_6xHIS_E

AAGAAGACTATACGGGTCTCAAAGGGAATGAAAGACAATACTGTGCCTCTGAAGCTGATCGCTCTCCTGGCTAAT
GGCGAGTTCCATAGTGGCGAACAGCTGGGAGAAACCCTGGGCATGTCCAGGGCCGCTATCAACAAGCACATTCAG

ACTCTGCGCGACTGGGGCGTGGACGTGTTCACCGTGCCCGGAAAGGGCTACTCTCTGCCCGAGCCTATCCCGCTG
CTGAACGCTAAACAGATTCTGGGACAGCTGGACGGCGGGAGCGTGGCAGTCCTGCCTGTGGTCGACTCCACCAAT
CAGTACCTGCTGGATCGAATCGGCGAGCTGAAGAGTGGGGATGCTTGCATTGCAGAATATCAGCAGGCAGGGAGA
GGAAGCAGAGGGAGGAAATGGTTCTCTCCTTTTGGAGCTAACCTGTACCTGAGTATGTTTTGGCGCCTGAAGCGG
GGACCAGCAGCAATCGGCCTGGGCCCGGTCATCGGAATTGTCATGGCAGAAGCGCTGCGAAAGCTGGGAGCAGAC
AAGGTGCGAGTCAAATGGCCCAATGACCTGTATCTGCAGGATAGAAAGCTGGCAGGCATCCTGGTGGAGCTGGCC
GGAATAACAGGCGATGCTGCACAGATCGTCATTGGCGCCGGGATTAACGTGGCTATGAGGCGCGTGGAGGAAAGC
GTGGTCAATCAGGGCTGGATCACACTGCAGGAAGCAGGGATTAACCTGGACAGGAATACTCTGGCCGCTACGCTG
ATCCGAGAGCTGCGGGCAGCCCTGGAACTGTTCGAGCAGGAAGGCCTGGCTCCATATCTGCCACGGTGGGAGAAG
CTGGATAACTTCATCAATAGACCCGTGAAGCTGATCATTGGGGACAAAGAGATTTTCGGGATTAGCCGGGGGATT
GATAAACAGGGAGCCCTGCTGCTGGAACAGGACGGAGTTATCAAACCCTGGATGGGCGGAGAAATCAGTCTGCGG
TCTGCCGAAAAGAAGCTTGCGGCCGCACTCGAGCACCACCACCACCACCACAATCTGAGACCACGAAGTGGGTA

B_TurboID_C

AAGAAGACTATACGGGTCTCATCTGAACAATGAAAGACAATACTGTGCCTCTGAAGCTGATCGCTCTCCTGGCTA
ATGGCGAGTTCCATAGTGGCGAACAGCTGGGAGAAACCCTGGGCATGTCCAGGGCCGCTATCAACAAGCACATTC
AGACTCTGCGCGACTGGGGCGTGGACGTGTTCACCGTGCCCGGAAAGGGCTACTCTCTGCCCGAGCCTATCCCGC
TGCTGAACGCTAAACAGATTCTGGGACAGCTGGACGGCGGGAGCGTGGCAGTCCTGCCTGTGGTCGACTCCACCA
ATCAGTACCTGCTGGATCGAATCGGCGAGCTGAAGAGTGGGGATGCTTGCATTGCAGAATATCAGCAGGCAGGGA
GAGGAAGCAGAGGGAGGAAATGGTTCTCTCCTTTTGGAGCTAACCTGTACCTGAGTATGTTTTGGCGCCTGAAGC
GGGGACCAGCAGCAATCGGCCTGGGCCCGGTCATCGGAATTGTCATGGCAGAAGCGCTGCGAAAGCTGGGAGCAG
ACAAGGTGCGAGTCAAATGGCCCAATGACCTGTATCTGCAGGATAGAAAGCTGGCAGGCATCCTGGTGGAGCTGG
CCGGAATAACAGGCGATGCTGCACAGATCGTCATTGGCGCCGGGATTAACGTGGCTATGAGGCGCGTGGAGGAAA
GCGTGGTCAATCAGGGCTGGATCACACTGCAGGAAGCAGGGATTAACCTGGACAGGAATACTCTGGCCGCTACGC
TGATCCGAGAGCTGCGGGCAGCCCTGGAACTGTTCGAGCAGGAAGGCCTGGCTCCATATCTGCCACGGTGGGAGA
AGCTGGATAACTTCATCAATAGACCCGTGAAGCTGATCATTGGGGACAAAGAGATTTTCGGGATTAGCCGGGGGA
TTGATAAACAGGGAGCCCTGCTGCTGGAACAGGACGGAGTTATCAAACCCTGGATGGGCGGAGAAATCAGTCTGC
GGTCTGCCGAAAAGAAGCTTGCGGCCGCACTCGAGGGCACCTGAGACCTCTGTAGTCTTCTT

D_TurboID_E

AAGAAGACTATACGGGTCTCAAAGGGAATGAAAGACAATACTGTGCCTCTGAAGCTGATCGCTCTCCTGGCTAAT
GGCGAGTTCCATAGTGGCGAACAGCTGGGAGAAACCCTGGGCATGTCCAGGGCCGCTATCAACAAGCACATTCAG
ACTCTGCGCGACTGGGGCGTGGACGTGTTCACCGTGCCCGGAAAGGGCTACTCTCTGCCCGAGCCTATCCCGCTG
CTGAACGCTAAACAGATTCTGGGACAGCTGGACGGCGGGAGCGTGGCAGTCCTGCCTGTGGTCGACTCCACCAAT
CAGTACCTGCTGGATCGAATCGGCGAGCTGAAGAGTGGGGATGCTTGCATTGCAGAATATCAGCAGGCAGGGAGA
GGAAGCAGAGGGAGGAAATGGTTCTCTCCTTTTGGAGCTAACCTGTACCTGAGTATGTTTTGGCGCCTGAAGCGG
GGACCAGCAGCAATCGGCCTGGGCCCGGTCATCGGAATTGTCATGGCAGAAGCGCTGCGAAAGCTGGGAGCAGAC
AAGGTGCGAGTCAAATGGCCCAATGACCTGTATCTGCAGGATAGAAAGCTGGCAGGCATCCTGGTGGAGCTGGCC
GGAATAACAGGCGATGCTGCACAGATCGTCATTGGCGCCGGGATTAACGTGGCTATGAGGCGCGTGGAGGAAAGC
GTGGTCAATCAGGGCTGGATCACACTGCAGGAAGCAGGGATTAACCTGGACAGGAATACTCTGGCCGCTACGCTG
ATCCGAGAGCTGCGGGCAGCCCTGGAACTGTTCGAGCAGGAAGGCCTGGCTCCATATCTGCCACGGTGGGAGAAG
CTGGATAACTTCATCAATAGACCCGTGAAGCTGATCATTGGGGACAAAGAGATTTTCGGGATTAGCCGGGGGATT
GATAAACAGGGAGCCCTGCTGCTGGAACAGGACGGAGTTATCAAACCCTGGATGGGCGGAGAAATCAGTCTGCGG
TCTGCCGAAAAGAAGCTTGCGGCCGCACTCGAGAATCTGAGACCACGAAGTGGGTA

B_TurboID-sGFP_C

GGTCTCaTCTGAACAATGAAAGACAATACTGTGCCTCTGAAGCTGATCGCTCTCCTGGCTAATGGCGAGTTCCAT
AGTGGCGAACAGCTGGGAGAAACCCTGGGCATGTCCAGGGCCGCTATCAACAAGCACATTCAGACTCTGCGCGAC
TGGGGCGTGGACGTGTTCACCGTGCCCGGAAAGGGCTACTCTCTGCCCGAGCCTATCCCGCTGCTGAACGCTAAA
CAGATTCTGGGACAGCTGGACGGCGGGAGCGTGGCAGTCCTGCCTGTGGTCGACTCCACCAATCAGTACCTGCTG
GATCGAATCGGCGAGCTGAAGAGTGGGGATGCTTGCATTGCAGAATATCAGCAGGCAGGGAGAGGAAGCAGAGGG
AGGAAATGGTTCTCTCCTTTTGGAGCTAACCTGTACCTGAGTATGTTTTGGCGCCTGAAGCGGGGACCAGCAGCA
ATCGGCCTGGGCCCGGTCATCGGAATTGTCATGGCAGAAGCGCTGCGAAAGCTGGGAGCAGACAAGGTGCGAGTC
AAATGGCCCAATGACCTGTATCTGCAGGATAGAAAGCTGGCAGGCATCCTGGTGGAGCTGGCCGGAATAACAGGC
GATGCTGCACAGATCGTCATTGGCGCCGGGATTAACGTGGCTATGAGGCGCGTGGAGGAAAGCGTGGTCAATCAG
GGCTGGATCACACTGCAGGAAGCAGGGATTAACCTGGACAGGAATACTCTGGCCGCTACGCTGATCCGAGAGCTG
CGGGCAGCCCTGGAACTGTTCGAGCAGGAAGGCCTGGCTCCATATCTGCCACGGTGGGAGAAGCTGGATAACTTC

ATCAATAGACCCGTGAAGCTGATCATTGGGGACAAAGAGATTTTCGGGATTAGCCGGGGGATTGATAAACAGGGA
GCCCTGCTGCTGGAACAGGACGGAGTTATCAAACCCTGGATGGGCGGAGAAATCAGTCTGCGGTCTGCCGAAAAG
AAGCTTGCGGCCGCACTCGAGAAGGGAGGTGGAGGAGGTTCTGGAGGCGGTGGAAGTGGTGGCGGAGGTAGCGTG
AGCAAGGGCGAGGAGCTGTTCACCGGGGTGGTGCCCATCCTGGTCGAGCTGGACGGCGACGTAAACGGCCACAAG
TTCAGCGTGTCCGGCGAGGGCGAGGGCGATGCCACCTACGGCAAGCTGACCCTGAAGTTCATCTGCACCACCGGC
AAGCTGCCCGTGCCCTGGCCCACCCTCGTGACCACCTTCACCTACGGCGTGCAGTGCTTCAGCCGCTACCCCGAC
CACATGAAGCAGCACGACTTCTTCAAGTCCGCCATGCCCGAAGGCTACGTCCAGGAGCGCACCATCTTCTTCAAG
GACGACGGCAACTACAAGACCCGCGCCGAGGTGAAGTTCGAGGGCGACACCCTGGTGAACCGCATCGAGCTGAAG
GGCATCGACTTCAAGGAGGACGGCAACATCCTGGGGCACAAGCTGGAGTACAACTACAACAGCCACAACGTCTAT
ATCATGGCCGACAAGCAGAAGAACGGCATCAAGGTGAACTTCAAGATCCGCCACAACATCGAGGACGGCAGCGTG
CAGCTCGCCGACCACTACCAGCAGAACACCCCCATCGGCGACGGCCCCGTGCTGCTGCCCGACAACCACTACCTG
AGCACCCAGTCCGCCCTGAGCAAAGACCCCAACGAGAAGCGCGATCACATGGTCCTGCTGGAGTTCGTGACCGCC
GCCGGGATCACTCACGGCATGGACGAGCTGTACAAGGGCACCtGAGACC

D_sGFP-TurboID_E

AAGAAGACTATACGGGTCTCAAAGGGAATGGTGAGCAAGGGCGAGGAGCTGTTCACCGGGGTGGTGCCCATCCTG
GTCGAGCTGGACGGCGACGTAAACGGCCACAAGTTCAGCGTGTCCGGCGAGGGCGAGGGCGATGCCACCTACGGC
AAGCTGACCCTGAAGTTCATCTGCACCACCGGCAAGCTGCCCGTGCCCTGGCCCACCCTCGTGACCACCTTCACC
TACGGCGTGCAGTGCTTCAGCCGCTACCCCGACCACATGAAGCAGCACGACTTCTTCAAGTCCGCCATGCCCGAA
GGCTACGTCCAGGAGCGCACCATCTTCTTCAAGGACGACGGCAACTACAAGACCCGCGCCGAGGTGAAGTTCGAG
GGCGACACCCTGGTGAACCGCATCGAGCTGAAGGGCATCGACTTCAAGGAGGACGGCAACATCCTGGGGCACAAG
CTGGAGTACAACTACAACAGCCACAACGTCTATATCATGGCCGACAAGCAGAAGAACGGCATCAAGGTGAACTTC
AAGATCCGCCACAACATCGAGGACGGCAGCGTGCAGCTCGCCGACCACTACCAGCAGAACACCCCCATCGGCGAC
GGCCCCGTGCTGCTGCCCGACAACCACTACCTGAGCACCCAGTCCGCCCTGAGCAAAGACCCCAACGAGAAGCGC
GATCACATGGTCCTGCTGGAGTTCGTGACCGCCGCCGGGATCACTCACGGCATGGACGAGCTGTACAAGGGTGGA
GGAGGTTCTGGAGGCGGTGGAAGTGGTGGCGGAGGTAGCGGCACCATGAAAGACAATACTGTGCCTCTGAAGCTG
ATCGCTCTCCTGGCTAATGGCGAGTTCCATAGTGGCGAACAGCTGGGAGAAACCCTGGGCATGTCCAGGGCCGCT
ATCAACAAGCACATTCAGACTCTGCGCGACTGGGGCGTGGACGTGTTCACCGTGCCCGGAAAGGGCTACTCTCTG
CCCGAGCCTATCCCGCTGCTGAACGCTAAACAGATTCTGGGACAGCTGGACGGCGGGAGCGTGGCAGTCCTGCCT
GTGGTCGACTCCACCAATCAGTACCTGCTGGATCGAATCGGCGAGCTGAAGAGTGGGGATGCTTGCATTGCAGAA
TATCAGCAGGCAGGGAGAGGAAGCAGAGGGAGGAAATGGTTCTCTCCTTTTGGAGCTAACCTGTACCTGAGTATG
TTTTGGCGCCTGAAGCGGGGACCAGCAGCAATCGGCCTGGGCCCGGTCATCGGAATTGTCATGGCAGAAGCGCTG
CGAAAGCTGGGAGCAGACAAGGTGCGAGTCAAATGGCCCAATGACCTGTATCTGCAGGATAGAAAGCTGGCAGGC
ATCCTGGTGGAGCTGGCCGGAATAACAGGCGATGCTGCACAGATCGTCATTGGCGCCGGGATTAACGTGGCTATG
AGGCGCGTGGAGGAAAGCGTGGTCAATCAGGGCTGGATCACACTGCAGGAAGCAGGGATTAACCTGGACAGGAAT
ACTCTGGCCGCTACGCTGATCCGAGAGCTGCGGGCAGCCCTGGAACTGTTCGAGCAGGAAGGCCTGGCTCCATAT
CTGCCACGGTGGGAGAAGCTGGATAACTTCATCAATAGACCCGTGAAGCTGATCATTGGGGACAAAGAGATTTTC
GGGATTAGCCGGGGGATTGATAAACAGGGAGCCCTGCTGCTGGAACAGGACGGAGTTATCAAACCCTGGATGGGC
GGAGAAATCAGTCTGCGGTCTGCCGAAAAGAAGCTTGCGGCCGCACTCGAGTGAGACCACGAAGTGGGTA

GGAG_ p35S_Linker_**BirA**-Myc_CaMV poly(A)_CGCT

CCATTGAGACTTTTCAACAAAGGATAATTTCGGGAAACCTCCTCGGATTCCATTGCCCAGCTATCTGTCACTTCA
TCGAAAGGACAGTAGAAAAGGAAGGTGGCTCCTACAAATGCCATCATTGCGATAAAGGAAAGGCTATCATTCAAG
ATCTCTCTGCCGACAGTGGTCCCAAAGATGGACCCCCACCCACGAGGAGCATCGTGGAAAAAGAAGAGGTTCCAA
CCACGTCTACAAAGCAAGTGGATTGATGTGACATCTCCACTGACGTAAGGGATGACGCACAATCCCACTATCCTT
CGCAAGACCCTTCCTCTATATAAGGAAGTTCATTTCATTTGGAGAGGACACGCTCGAGTATAAGAGCTCATTTTT
ACAACAATTACCAACAACAACAAACAACAAACAACATTACAATTACATTTACAATTATCGATACCCATccatGGA
GAGTAAACGGAATAAGCCAGGGAAGGCGACAGGTAAAGGTAAACCAGTTGGTGATAAATGGCTGGATGATGCAGG
TAAAGATTCAGGAGCGCCAATTCCAGATCGCATTGCTGATAAGTTGCGTGATAAAGAATTTAAAAGCTTCGACGA
TTTTCGGAAGGCTGTATGGGAAGAGGTGTCGAAAGATCCTGAGCTTAGTAAAAATTTAAACCCAAGCAATAAGTC
TAGTGTTTCAAAAGGTTATTCTCCGTTTACTCCAAAGAATCAACAGGTCGGAGGGAGAAAAGTCTATGAACTTCA
TCATGACAAGCCAATTAGTCAAGGTGGTGAGGTTTATGACATGGATAATATCCGAGTGACTACACCTAAGCGACA
TATCGATATTCACCGAGGTAAGGTTTCGGGATCTGGAGGTGGCGGAAGCGGAGGCGGAGGATCCGGAGGTGGAGG
TAGCGGCGGTGGAGGCGGTAGTGGAGGAGGAGGGTCTGGTGGAGGTGGCTCAGGTGGTGGTGGGTCAGGCGGTGG
TGGTAGTGGCGGAGGAGGATCGGGAGGAGGTGGGTCAGGTGGCGGAGGATCTGGAGGGGGTGGTTCTGGTGGTGG
TGGTTCAGCAAGGAGATCTCATTCGGGAAACGCGGCTATTAGATCAAAGGATAACACCGTGCCACTTAAATTGAT
TGCCCTTTTAGCGAACGGTGAATTTCACTCTGGTGAGCAGTTGGGTGAAACGCTTGGAATGAGCAGAGCTGCTAT
TAATAAACACATTCAGACTCTTCGTGACTGGGGAGTTGATGTCTTTACCGTTCCAGGTAAAGGATACAGCCTGCC
TGAGCCTATCCAGTTACTTAATGCTAAACAGATATTGGGTCAGCTTGATGGAGGTAGTGTAGCTGTGCTGCCAGT

GATTGACTCCACGAATCAGTACCTTCTTGATCGTATCGGAGAGCTTAAATCTGGTGATGCTTGCATTGCAGAATA
CCAGCAGGCTGGTCGTGGAGGTAGAGGTCGTAAATGGTTTTCACCTTTTGGTGCAAACTTGTATTTGTCAATGTT
CTGGCGTCTTGAACAAGGTCCAGCTGCAGCTATTGGTTTAAGTCTTGTTATCGGTATCGTGATGGCGGAAGTATT
ACGTAAGCTTGGTGCAGATAAAGTTCGTGTTAAATGGCCTAATGACCTCTATCTTCAGGATCGTAAGCTTGCAGG
AATTCTTGTGGAGCTCACTGGCAAAACTGGTGATGCAGCTCAAATAGTCATTGGAGCTGGTATCAACATGGCAAT
GAGGCGTGTTGAAGAGAGTGTCGTTAATCAGGGATGGATCACGCTGCAGGAAGCTGGTATCAATCTCGATCGTAA
TACGTTGGCAGCTATGCTAATACGTGAATTACGTGCTGCGTTGGAACTCTTCGAACAAGAAGGATTGGCACCTTA
TCTTTCACGTTGGGAAAAGCTCGATAATTTTATTAATAGACCAGTGAAACTTATCATTGGTGATAAAGAAATATT
TGGTATTTCACGTGGAATAGACAAACAGGGTGCTTTGTTACTTGAGCAGGATGGAATAATAAAACCTTGGATGGG
AGGTGAAATATCCCTTCGTAGTGCAGAAAAAGAACAAAAACTCATCTCAGAAGAGGATCTGTAAGCTTCTCTAGC
TAGAGTCGATCGACAAGCTCGAGTTTCTCCATAATAATGTGTGAGTAGTTCCCAGATAAGGGAATTAGGGTTCCT
ATAGGGTTTCGCTCATGTGTTGAGCATATAAGAAACCCTTAGTATGTATTTGTATTTGTAAAATACTTCTATCAA
TAAAATTTCTAATTCCTAAAACCAAAATCCAGTACTAAAATCCAGATCGCTACGCT

GGAG_ p35S_Linker_**BioID**-Myc_CaMV poly(A)_CGCT

CCATTGAGACTTTTCAACAAAGGATAATTTCGGGAAACCTCCTCGGATTCCATTGCCCAGCTATCTGTCACTTCA
TCGAAAGGACAGTAGAAAAGGAAGGTGGCTCCTACAAATGCCATCATTGCGATAAAGGAAAGGCTATCATTCAAG
ATCTCTCTGCCGACAGTGGTCCCAAAGATGGACCCCCACCCACGAGGAGCATCGTGGAAAAAGAAGAGGTTCCAA
CCACGTCTACAAAGCAAGTGGATTGATGTGACATCTCCACTGACGTAAGGGATGACGCACAATCCCACTATCCTT
CGCAAGACCCTTCCTCTATATAAGGAAGTTCATTTCATTTGGAGAGGACACGCTCGAGTATAAGAGCTCATTTTT
ACAACAATTACCAACAACAACAAACAACAAACAACATTACAATTACATTTACAATTATCGATACCCATCCATGGA
GAGTAAACGGAATAAGCCAGGGAAGGCGACAGGTAAAGGTAAACCAGTTGGTGATAAATGGCTGGATGATGCAGG
TAAAGATTCAGGAGCGCCAATTCCAGATCGCATTGCTGATAAGTTGCGTGATAAAGAATTTAAAAGCTTCGACGA
TTTTCGGAAGGCTGTATGGGAAGAGGTGTCGAAAGATCCTGAGCTTAGTAAAAATTTAAACCCAAGCAATAAGTC
TAGTGTTTCAAAAGGTTATTCTCCGTTTACTCCAAAGAATCAACAGGTCGGAGGGAGAAAAGTCTATGAACTTCA
TCATGACAAGCCAATTAGTCAAGGTGGTGAGGTTTATGACATGGATAATATCCGAGTGACTACACCTAAGCGACA
TATCGATATTCACCGAGGTAAGGTTTCGGGATCTGGAGGTGGCGGAAGCGGAGGCGGAGGATCCGGAGGTGGAGG
TAGCGGCGGTGGAGGCGGTAGTGGAGGAGGAGGGTCTGGTGGAGGTGGCTCAGGTGGTGGTGGGTCAGGCGGTGG
TGGTAGTGGCGGAGGAGGATCGGGAGGAGGTGGGTCAGGTGGCGGAGGATCTGGAGGGGGTGGTTCTGGTGGTGG
TGGTTCAGCAAGGAGAAACCATGGAACAAAAACTCATCTCAGAAGAGGATCTGAAGGATAACACCGTGCCACTTA
AATTGATTGCCCTTTTAGCGAACGGTGAATTTCACTCTGGTGAGCAGTTGGGTGAAACGCTTGGAATGAGCAGAG
CTGCTATTAATAAACACATTCAGACTCTTCGTGACTGGGGAGTTGATGTCTTTACCGTTCCAGGTAAAGGATACA
GCCTGCCTGAGCCTATCCAGTTACTTAATGCTAAACAGATATTGGGTCAGCTTGATGGAGGTAGTGTAGCTGTGC
TGCCAGTGATTGACTCCACGAATCAGTACCTTCTTGATCGTATCGGAGAGCTTAAATCTGGTGATGCTTGCATTG
CAGAATACCAGCAGGCTGGTCGTGGACGTAGAGGTCGTAAATGGTTTTCACCTTTTGGTGCAAACTTGTATTTGT
CAATGTTCTGGCGTCTTGAACAAGGTCCAGCTGCAGCTATTGGTTTAAGTCTTGTTATCGGTATCGTGATGGCGG
AAGTATTACGTAAGCTTGGTGCAGATAAAGTTCGTGTTAAATGGCCTAATGACCTCTATCTTCAGGATCGTAAGC
TTGCAGGAATTCTTGTGGAGCTCACTGGCAAAACTGGTGATGCAGCTCAAATAGTCATTGGAGCTGGTATCAACA
TGGCAATGAGGCGTGTTGAAGAGAGTGTCGTTAATCAGGGATGGATCACGCTGCAGGAAGCTGGTATCAATCTCG
ATCGTAATACGTTGGCAGCTATGCTAATACGTGAATTACGTGCTGCGTTGGAACTCTTCGAACAAGAAGGATTGG
CACCTTATCTTTCACGTTGGGAAAAGCTCGATAATTTTATTAATAGACCAGTGAAACTTATCATTGGTGATAAAG
AAATATTTGGTATTTCACGTGGAATAGACAAACAGGGTGCTTTGTTACTTGAGCAGGATGGAATAATAAAACCTT
GGATGGGAGGTGAAATATCCCTTCGTAGTGCAGAAAAAGGAAACGCGGCTATTAGATCAGGAATGGAACAAAAAC
TCATCTCAGAAGAGGATCTGTAAGCTTCTCTAGCTAGAGTCGATCGACAAGCTCGAGTTTCTCCATAATAATGTG
TGAGTAGTTCCCAGATAAGGGAATTAGGGTTCCTATAGGGTTTCGCTCATGTGTTGAGCATATAAGAAACCCTTA
GTATGTATTTGTATTTGTAAAATACTTCTATCAATAAAATTTCTAATTCCTAAAACCAAAATCCAGTACTAAAAT
CCAGATCGCTACGCT

GGAG_ p35S_Linker_**BioID2**-HA_CaMV poly(A)_CGCT

GGAGTGAGACTTTTCAACAAAGGATAATTTCGGGAAACCTCCTCGGATTCCATTGCCCAGCTATCTGTCACTTCA
TCGAAAGGACAGTAGAAAAGGAAGGTGGCTCCTACAAATGCCATCATTGCGATAAAGGAAAGGCTATCATTCAAG
ATCTCTCTGCCGACAGTGGTCCCAAAGATGGACCCCCACCCACGAGGAGCATCGTGGAAAAAGAAGAGGTTCCAA
CCACGTCTACAAAGCAAGTGGATTGATGTGACATCTCCACTGACGTAAGGGATGACGCACAATCCCACTATCCTT
CGCAAGACCCTTCCTCTATATAAGGAAGTTCATTTCATTTGGAGAGGACACGCTCGAGTATAAGAGCTCATTTTT
ACAACAATTACCAACAACAACAAACAACAAACAACATTACAATTACATTTACAATTATCGATACCCATccatGGA
GAGTAAACGGAATAAGCCAGGGAAGGCGACAGGTAAAGGTAAACCAGTTGGTGATAAATGGCTGGATGATGCAGG
TAAAGATTCAGGAGCGCCAATTCCAGATCGCATTGCTGATAAGTTGCGTGATAAAGAATTTAAAAGCTTCGACGA
TTTTCGGAAGGCTGTATGGGAAGAGGTGTCGAAAGATCCTGAGCTTAGTAAAAATTTAAACCCAAGCAATAAGTC
TAGTGTTTCAAAAGGTTATTCTCCGTTTACTCCAAAGAATCAACAGGTCGGAGGGAGAAAAGTCTATGAACTTCA

TCATGACAAGCCAATTAGTCAAGGTGGTGAGGTTTATGACATGGATAATATCCGAGTGACTACACCTAAGCGACA
TATCGATATTCACCGAGGTAAGGTTTCGGGATCTGGAGGTGGCGGAAGCGGAGGCGGAGGATCCGGAGGTGGAGG
TAGCGGCGGTGGAGGCGGTAGTGGAGGAGGAGGGTCTGGTGGAGGTGGCTCAGGTGGTGGTGGGTCAGGCGGTGG
TGGTAGTGGCGGAGGAGGATCGGGAGGAGGTGGGTCAGGTGGCGGAGGATCTGGAGGGGGTGGTTCTGGTGGTGG
TGGTTCAGCAAGGAGATTTAAGAACTTGATATGGCTGAAAGAAGTAGATTCCACACAAGAGAGACTTAAAGAATG
GAATGTCTCCTATGGGACTGCACTTGTTGCCGATAGGCAAACCAAAGGAAGAGGCGGGTTAGGACGTAAATGGTT
ATCTCAAGAAGGCGGTCTATACTTCAGTTTTCTTCTGAATCCTAAAGAGTTTGAGAACTTGCTTCAGCTTCCTTT
GGTGTTGGGTTTATCGGTTTCGGAGGCTCTGGAGGAAATCACTGAGATCCCATTTTCTCTCAAATGGCCTAATGA
CGTTTACTTCCAGGAGAAGAAAGTATCTGGTGTGCTTTGTGAACTCTCAAAGGACAAGCTAATTGTTGGAATCGG
TATTAACGTCAATCAGAGAGAAATTCCGGAAGAGATCAAAGATCGAGCTACAACACTATACGAGATTACCGGTAA
GGATTGGGACAGAAAGGAGGTCCTACTCAAAGTGTTAAAACGGATATCAGAGAACTTGAAGAAGTTCAAGGAAAA
GAGCTTCAAAGAATTCAAAGGGAAGATAGAATCCAAAATGTTGTACCTTGGGGAAGAAGTGAAGCTTTTGGGAGA
AGGAAAGATAACGGGAAAGCTCGTTGGGCTTAGTGAGAAAGGGGGGGCTTTGATTCTCACTGAAGAGGGAATTAA
GGAGATCTTATCTGGGGAATTTAGTCTCAGGCGTAGCTATCCCTATGATGTTCCAGATTATGCGTAAGCTTCTCT
AGCTAGAGTCGATCGACAAGCTCGAGTTTCTCCATAATAATGTGTGAGTAGTTCCCAGATAAGGGAATTAGGGTT
CCTATAGGGTTTCGCTCATGTGTTGAGCATATAAGAAACCCTTAGTATGTATTTGTATTTGTAAAATACTTCTAT
CAATAAAATTTCTAATTCCTAAAACCAAAATCCAGTACTAAAATCCAGATCGCT

GGAG_ p35S_sGFP_Linker_**TurboID**_6xHIS-3xFLAG_CaMV poly(A)_CGCT

GGAGGAATTCCAATCCCACAAAAATCTGAGCTTAACAGCACAGTTGCTCCTCTCAGAGCAGAATCGGGTATTCAA
CACCCTCATATCAACTACTACGTTGTGTATAACGGTCCACATGCCGGTATATACGATGACTGGGGTTGTACAAAG
GCGGCAACAAACGGCGTTCCCGGAGTTGCACACAAGAAATTTGCCACTATTACAGAGGCAAGAGCAGCAGCTGAC
GCGTACACAACAAGTCAGCAAACAGACAGGTTGAACTTCATCCCCAAAGGAGAAGCTCAACTCAAGCCCAAGAGC
TTTGCTAAGGCCCTAACAAGCCCACCAAAGCAAAAAGCCCACTGGCTCACGCTAGGAACCAAAAGGCCCAGCAGT
GATCCAGCCCCAAAAGAGATCTCCTTTGCCCCGGAGATTACAATGGACGATTTCCTCTATCTTTACGATCTAGGA
AGGAAGTTCGAAGGTGAAGGTGACGACACTATGTTCACCACTGATAATGAGAAGGTTAGCCTCTTCAATTTCAGA
AAGAATGCTGACCCACAGATGGTTAGAGAGGCCTACGCAGCAAGTCTCATCAAGACGATCTACCCGAGTAACAAT
CTCCAGGAGATCAAATACCTTCCCAAGAAGGTTAAAGATGCAGTCAAAAGATTCAGGACTAATTGCATCAAGAAC
ACAGAGAAAGACATATTTCTCAAGATCAGAAGTACTATTCCAGTATGGACGATTCAAGGCTTGCTTCATAAACCA
AGGCAAGTAATAGAGATTGGAGTCTCTAAAAAGGTAGTTCCTACTGAATCTAAGGCCATGCATGGAGTCTAAGAT
TCAAATCGAGGATCTAACAGAACTCGCCGTCAAGACTGGCGAACAGTTCATACAGAGTCTTTTACGACTCAATGA
CAAGAAGAAAATCTTCGTCAACATGGTGGAGCACGACACTCTGGTCTACTCCAAAAATGTCAAAGATACAGTCTC
AGAAGATCAAAGGGCTATTGAGACTTTTCAACAAAGGATAATTTCGGGAAACCTCCTCGGATTCCATTGCCCAGC
TATCTGTCACTTCATCGAAAGGACAGTAGAAAAGGAAGGTGGCTCCTACAAATGCCATCATTGCGATAAAGGAAA
GGCTATCATTCAAGATCTCTCTGCCGACAGTGGTCCCAAAGATGGACCCCCACCCACGAGGAGCATCGTGGAAAA
AGAAGAGGTTCCAACCACGTCTACAAAGCAAGTGGATTGATGTGACATCTCCACTGACGTAAGGGATGACGCACA
ATCCCACTATCCTTCGCAAGACCCTTCCTCTATATAAGGAAGTTCATTTCATTTGGAGAGGACACGCAATGGGCC
CCGACATCGTGATGACCCAGAGCCCCAGCAGCCTGAGCGCCAGCGTGGGCGACCGCGTGACCATCACCTGCCGCA
GCAGCACCGGCGCCGTGACCACCAGCAACTACGCCAGCTGGGTGCAGGAGAAGCCCGGCAAGCTGTTCAAGGGCC
TGATCGGCGGCACCAACAACCGCGCCCCCGGCGTGCCCAGCCGCTTCAGCGGCAGCCTGATCGGCGACAAGGCCA
CCCTGACCATCAGCAGCCTGCAGCCCGAGGACTTCGCCACCTACTTCTGCGCCCTGTGGTACAGCAACCACTGGG
TGTTCGGCCAGGGCACCAAGGTGGAGCTGAAGCGCGGCGGCGGCGGCAGCGGCGGCGGCGGCAGCGAGGTGAAGC
TGCTGGAGAGCGGCGGCGGCCTGGTGCAGCCCGGCGGCAGCCTGAAGCTGAGCTGCGCCGTGAGCGGCTTCAGCC
TGACCGACTACGGCGTGAACTGGGTGCGCCAGGCCCCCGGCCGCGGCCTGGAGTGGATCGGCGTGATCTGGGGCG
ACGGCATCACCGACTACAACAGCGCCCTGAAGGACCGCTTCATCATCAGCAAGGACAACGGCAAGAACACCGTGT
ACCTGCAGATGAGCAAGGTGCGCAGCGACGACACCGCCCTGTACTACTGCGTGACCGGCCTGTTCGACTACTGGG
GCCAGGGCACCCTGGTGACCGTGAGCAGCTACCCATACGATGTTCCAGATTACGCTGGTGGAGGCGGAGGTTCTG
GGGGAGGAGGTAGTGGCGGTGGTGGTTCAGGAGGCGGCGGAAGCTTGGATCCAGGTGGAGGTGGAAGCGGTAGCA
AAGGAGAAGAACTTTTCACTGGAGTTGTCCCAATTCTTGTTGAATTAGATGGTGATGTTAATGGGCACAAATTTT
CTGTCCGTGGAGAGGGTGAAGGTGATGCTACAAACGGAAAACTCACCCTTAAATTTATTTGCACTACTGGAAAAC
TACCTGTTCCGTGGCCAACACTTGTCACTACTCTGACCTATGGTGTTCAATGCTTTTCCCGTTATCCGGATCACA
TGAAACGGCATGACTTTTTCAAGAGTGCCATGCCCGAAGGTTATGTACAGGAACGCACTATATCTTTCAAAGATG
ACGGGACCTACAAGACGCGTGCTGAAGTCAAGTTTGAAGGTGATACCCTTGTTAATCGTATCGAGTTAAAGGGTA
TTGATTTTAAAGAAGATGGAAACATTCTTGGACACAAACTCGAGTACAACTTTAACTCACACAATGTATACATCA
CGGCAGACAAACAAAAGAATGGAATCAAAGCTAACTTCAAAATTCGCCACAACGTTGAAGATGGTTCCGTTCAAC
TAGCAGACCATTATCAACAAAATACTCCAATTGGCGATGGCCCTGTCCTTTTACCAGACAACCATTACCTGTCGA
CACAATCTGTCCTTTCGAAAGATCCCAACGAAAAGCGTGACCACATGGTCCTTCTTGAGTTTGTAACTGCTGCTG
GGATTACACATGGCATGGATGAGCTCTACAAAGGTGGTTCGGGTTCTGGAGGAGGTGGATCTGCAAGCAAGGACA
ATACTGTGCCTTTGAAGCTAATAGCTTTGCTTGCGAATGGAGAATTCCATTCCGGGGAACAACTTGGCGAAACAT
TAGGGATGTCACGTGCTGCTATCAACAAGCACATTCAGACTCTAAGGGATTGGGGTGTTGATGTGTTCACTGTCC

CTGGCAAAGGATACTCTCTCCCTGAACCTATTCCCCTTCTTAACGCTAAGCAGATACTGGGGCAACTCGATGGTG
GTAGTGTGGCGGTTTTGCCAGTTGTGGATTCAACGAACCAGTATCTTCTCGATCGTATCGGAGAGCTTAAAAGTG
GAGATGCGTGTATAGCAGAGTACCAACAAGCTGGTAGAGGTTCACGAGGACGGAAATGGTTTTCACCCTTTGGCG
CCAATTTGTACTTGAGCATGTTTTGGCGACTCAAAAGAGGTCCAGCTGCAATAGGTCTTGGACCTGTCATCGGCA
TCGTTATGGCAGAAGCGTTAAGGAAACTTGGTGCCGATAAGGTACGTGTAAAATGGCCAAATGATCTCTATTTAC
AGGATCGTAAGCTAGCAGGAATTCTGGTTGAACTAGCAGGCATTACCGGAGATGCTGCTCAGATTGTCATTGGAG
CTGGAATTAACGTTGCCATGAGGAGAGTCGAGGAATCGGTTGTAAATCAAGGTTGGATCACACTTCAAGAAGCTG
GGATTAACCTAGACAGAAATACGCTTGCTGCAACCCTCATTAGGGAGTTACGGGCTGCTTTGGAACTCTTTGAGC
AGGAAGGTTTGGCTCCATATCTGCCTAGATGGGAGAAACTCGATAACTTCATCAATAGACCGGTTAAGCTGATAA
TTGGAGACAAGGAGATCTTTGGGATATCTAGAGGTATCGACAAACAAGGTGCATTGCTTTTAGAGCAAGATGGGG
TGATCAAGCCATGGATGGGCGGAGAGATAAGCTTGAGATCGGCCGAAAAGTTGCAAGGGAGTGGTAGAGGATCCC
ATCATCATCACCATCACGACTATGATATTCCGACTACAGCCTCTGAGAACCTGTATTTCCAGGGTGAGTTAGATT
ACAAAGACCATGATGGAGATTATAAGGATCACGATATCGACTACAAAGATGACGACGATAAATAAGCTTCGAACT
CTAGCTAGAGTCGATCGACAAGCTCGAGTTTCTCCATAATAATGTGTGAGTAGTTCCCAGATAAGGGAATTAGGG
TTCCTATAGGGTTTCGCTCATGTGTTGAGCATATAAGAAACCCTTAGTATGTATTTGTATTTGTAAAATACTTCT
ATCAATAAAATTTCTAATTCCTAAAACCAAAATCCAGTACTAAAATCCAGATCGCT
